# A viral ADP-ribosyltransferase attaches RNA chains to host proteins

**DOI:** 10.1101/2021.06.04.446905

**Authors:** Maik Wolfram-Schauerte, Nadiia Pozhydaieva, Julia Grawenhoff, Luisa M. Welp, Ivan Silbern, Alexander Wulf, Franziska A. Billau, Timo Glatter, Henning Urlaub, Andres Jäschke, Katharina Höfer

## Abstract

The mechanisms by which viruses hijack their host’s genetic machinery are of current interest. When bacteriophage T4 infects *Escherichia coli*, three different ARTs (ADP-ribosyltransferases) reprogram the host’s transcriptional and translational apparatus through ADP-ribosylation using nicotinamide adenine dinucleotide (NAD) as substrate ^1,2^. Recently, NAD was identified as a 5’-modification of cellular RNAs ^3–5^. Here, we report that T4 ART ModB accepts not only NAD but also NAD-capped RNA (NAD-RNA) as substrate and attaches entire RNA chains to acceptor proteins in an “RNAylation” reaction. ModB specifically RNAylates ribosomal proteins rS1 and rL2 at defined arginine residues, and a specific group of *E. coli* and T4 phage RNAs is linked to rS1 *in vivo*. T4 phages that express an inactive mutant of ModB show a decreased burst size and slowed lysis of *E. coli*. Our findings reveal a distinct biological role of NAD-RNA, namely activation of the RNA for enzymatic transfer to proteins. The attachment of specific RNAs to ribosomal proteins might provide a strategy for the phage to modulate the host’s translation machinery. This work exemplifies the first direct connection between RNA modification and post-translational protein modification. As ARTs play important roles far beyond viral infections ^6^, RNAylation may have far-reaching implications.

---

ARTs catalyse the transfer of one or multiple ADP-ribose (ADPr) units from NAD to target proteins ^7^. In bacteria and archaea, they act as toxins and are involved in host defence or drug resistance mechanisms ^8^, while in eukaryotes, they play roles in distinct processes ranging from DNA damage repair to macrophage activation and stress response ^9^. Viruses use ARTs as weapons to reprogram the host’s gene expression system ^6^. Mechanistically, a nucleophilic residue of the target protein (mostly Arg, Glu, Asp, Ser, Cys) attacks the glycosidic carbon atom in the nicotinamide riboside moiety of NAD, forming a covalent bond as N-, O-, or S-glycoside (Fig. 1a) ^7^. As the adenosine moiety of NAD is not involved in this reaction, we speculated that elongation of the adenosine to long RNA chains (via regular 5’-3’-phosphodiester bonds) might be tolerated by ARTs, potentially leading to the formation of covalent RNA-protein conjugates (Fig. 1b). RNAs harbouring a 5’-NAD-cap were recently found in bacteria, including *E. coli* ^3,10,11^, archaea ^12,13^, and higher organisms ^5,14–19^, where NAD-RNA concentrations range from 1.9 to 7.4 fmol/μg RNA ^16^. The modification was observed in different RNA types, including messenger RNAs (mRNAs) and small regulatory RNAs (sRNAs) ^20^. However, very little is known about the biological functions of this RNA cap ^21^.

**Fig. 1.**
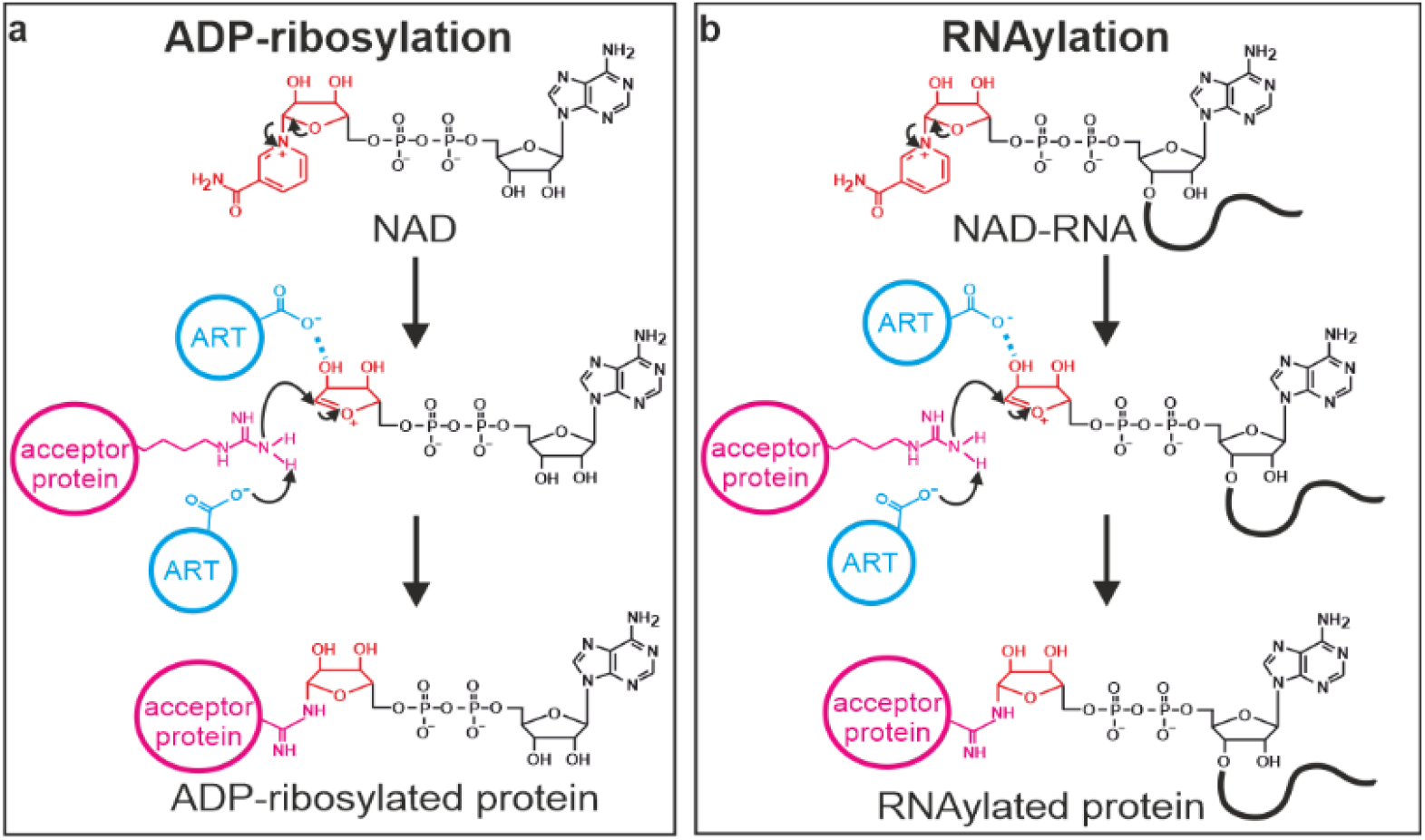
Mechanisms of ADP-ribosylation and proposed “RNAylation”. **a**, The mechanism of ADP-ribosylation is shown exemplarily for arginine. Initially, the N-glycosidic bond between the ribose and nicotinamide is destabilised by a glutamate residue of an ART. This leads to the formation of an oxocarbenium ion of ADP-ribose. Nicotinamide serves as the leaving group. This electrophilic ion is attacked by a nucleophilic arginine residue of the acceptor protein after glutamate-mediated proton abstraction. This leads to the formation of an N-glycosidic bond ^49^. **b**, Proposed RNAylation reaction mechanism. Analogous to ADP-ribosylation in the presence of NAD, we propose that ARTs might use NAD-RNA to catalyse an “RNAylation” reaction, thereby covalently attaching an RNA to an acceptor protein.

The infection cycle of bacteriophage T4 relies on the sequential expression of early, middle and late phage genes that are transcribed by *E. coli* RNA polymerase (RNAP) ^22^. For the specific temporal reprogramming of the *E. coli* transcriptional and translational apparatus, the T4 phage uses three ARTs that modify over 30 host proteins: upon infection, Alt is injected into the bacterium together with the phage DNA and immediately ADP-ribosylates *E. coli* RNAP at different residues, thought to result in the preferential transcription of phage genes from “early” promoters ^23,24^. Two early phage genes code for the ARTs ModA ^25^ and ModB ^1,26^. The former completes ADP-ribosylation of RNAP, while the latter is described to modify the host protein rS1 ^1,26^. However, it is still unknown how ADP-ribosylation changes the target proteins’ properties and whether additional proteins are modified upon T4 phage infection.

## RESULTS

### T4 ART ModB catalyses RNAylation *in vitro*

To test our hypothesis that ARTs may accept NAD-RNAs as substrates, we purified the three T4 ARTs Alt, ModA and ModB. We incubated them with a synthetic, site-specifically ^32^P-labelled 5’-NAD-RNA 8mer or with a 3’-fluorophore-labelled 5’-NAD-RNA 10mer to test for either self-modification or modification of target proteins. While both Alt and ModA showed only a low extent of target RNAylation (Extended Data Fig. 1a), ModB rapidly RNAylated its known ADP-ribosylation target protein rS1, without detectable self-RNAylation (Fig. 2a and Extended Data Fig. 1b). In contrast, ModB-mediated ADP-ribosylation in the presence of ^32^P-NAD resulted in the modification of both proteins (ModB and rS1) with similar intensities (Fig. 2b and Extended Data Fig. 1c). No signal appeared when either ModB or rS1 were missing or when a 5’-^32^P-monophosphate-RNA (5’-^32^P-RNA) of the same sequence was used as a substrate for ModB (Extended Data Fig. 1d). Moreover, mutation of a catalytically important residue (R73A, G74A), located in the active site of ModB, also prevented RNAylation of rS1 (Extended Data Fig. 1e,f) ^1^.

**Fig. 2.**
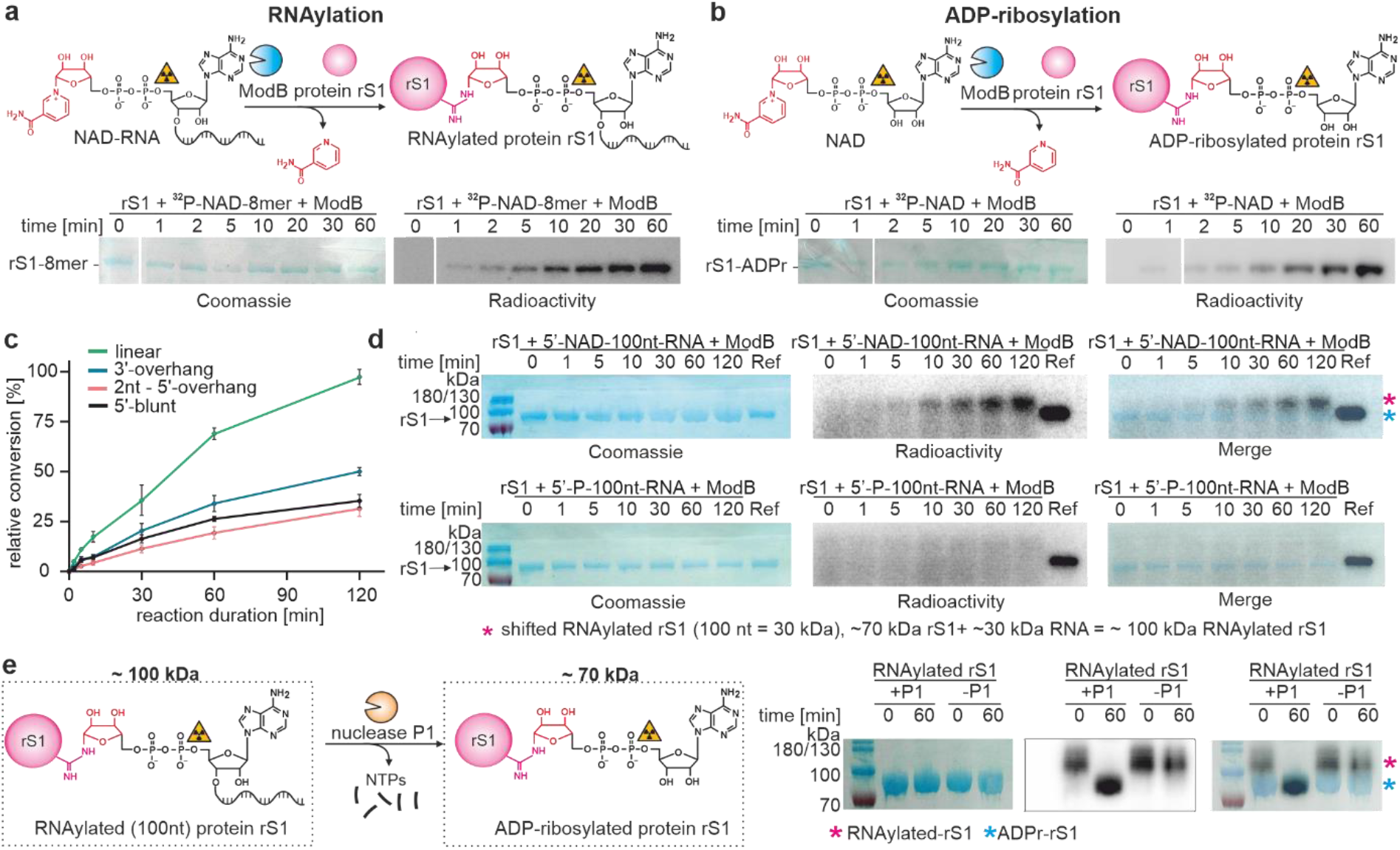
Post-translational protein modification of rS1 by the ART ModB *in vitro*. **a**, Time course of the RNAylation of rS1 by ModB (complete SDS-PAGE gels are shown in Extended Data Fig. 1b) (n=3). **b**, Time course of the ADP-ribosylation of rS1 by ModB (complete SDS-PAGE gels are shown in Extended Data Fig. 1c) (n=3). rS1 RNAylation and ADP-ribosylation is indicated by the acquisition of a radioactive signal overlapping with the Coomassie stain. **c**, Analysis of the role of RNA secondary structure on RNAylation reaction. Four different 3’-Cy5 labelled NAD-capped RNAs were tested including a linear (green) NAD-capped RNA (10mer) and three structured NAD-capped RNAs with either a 3’-overhang (blue), a 5’-overhang (red) or a blunt end (black). SDS-PAGE analysis is shown in Extended Data Fig. 3a. Data points represent mean ± standard deviation (s.d.), based on quantification of fluorescence Cy5 signals (n=3). **d**, *in vitro* kinetics of the RNAylation of rS1 by ModB using 5’-NAD-100nt-RNA as substrate (top panel), analysed by SDS-PAGE. Shifted RNAylated rS1 is highlighted with a pink asterisk. 5’-P-100nt-RNA is used as a negative control (bottom panel) (n=2). **e**, Nuclease P1 digest of RNAylated protein rS1. The covalently attached 100 nt long RNA results in a shift of the RNAylated protein rS1 (~100 kDa) in SDS-PAGE. Treatment of the RNAylated protein rS1 with nuclease P1, which cleaves the phosphodiester bond, resulting in degradation of the attached RNA into mononucleotides. Nuclease P1 coverts RNAylated rS1 into ADP-ribosylated rS1 (~70 kDa), which can be visualised by a downshifted protein band on the SDS-PAGE gel (n=1).

### RNAylation by ModB proceeds by an ADP-ribosylation-like mechanism

ModB-catalysed RNAylation of rS1 was strongly inhibited by the ART inhibitor 3-methoxybenzamide (3-MB) ^27^, which is thought to mimic the nicotinamide moiety (Extended Data Fig. 2a), confirming an ADP-ribosylation-like mechanism. Moreover, RNAylated rS1 proteins that carry a ^32^P-labelled ADP-ribose moiety were treated with RNase T1 to verify if the RNA and the protein are covalently linked (Extended Data Fig. 2b). This treatment would remove the ^32^P-label if the RNA was non-covalently bound to rS1 or was covalently linked via other than 5’-terminal positions. The ^32^P-rS1 signal did not disappear upon treatment with RNase T1. The signal, however, disappeared entirely upon treatment with trypsin, which digests rS1 (Extended Data Fig. 2c). Collectively, these data strongly support the covalent linkage of the RNA to rS1 via its 5’-end as shown in Fig. 1b.

RNAylation assays using short linear or hairpin-forming NAD-RNAs (Fig. 2c, Extended Data Fig. 3a) revealed a substrate preference of ModB for unstructured NAD-RNAs. ModB also accepted longer, biologically relevant NAD-capped RNAs as substrates (e.g., an NAD-capped Qβ-RNA fragment of ~ 100nt ^28^, Fig. 2d and Extended Data Fig. 3b). RNAylation with NAD-capped 100nt-RNA shifted protein rS1 (~ 70 kDa) to 100 kDa MW protein (Fig. 2e). Treatment of the RNAylated protein with nuclease P1, which hydrolyses 3’-5’ phosphodiester bonds but does not attack the pyrophosphate bond of the 5’-ADP-ribose, reverted this shift, and the ^32^P-labelled product migrated like unmodified rS1 or ADPr-rS1 (Fig. 2e), again confirming the proposed nature of the covalent linkage.

To exclude the possibility that ModB just removes the nicotinamide moiety from the NAD-RNA by hydrolysis, thus generating a highly reactive ribosyl moiety that could (via its masked aldehyde group) spontaneously react with nucleophiles in its vicinity ^29^, we prepared ADP-ribose-modified RNA and tested it as a substrate for ModB. No modification could be detected (Extended Data Fig. 3c), providing no support for spontaneous RNAylation.

To exclude RNA degradation during RNAylation, we supplied ModB with a NAD-RNA 10mer that carried a fluorescent dye (Cy5) at the 3’-terminus (Extended Data Fig. 1e, Extended Data Fig. 3a). The time course analysis of the RNAylation indicated the attachment of intact oligonucleotide chains to rS1 for a variety of NAD-capped RNAs (Extended Data Fig. 3a).

### ModB modifies specific arginine residues in rS1

To identify the amino acid residues in protein rS1 to which RNA chains are covalently linked during RNAylation, we took advantage of tools developed to analyse protein ADP-ribosylation.

The radioactive signal of ^32^P-RNAylated protein rS1 or ^32^P–ADP-ribosylated rS1 did not change upon treatment with HgCl_2_ (which cleaves S-glycosides at Cys), NH_2_OH (which hydrolyses O-glycosides at Asp and Glu) (Extended Data Fig. 4a) and recombinant enzyme ARH3 (which hydrolyses O-ADPr glycosides specifically at serine residues) (Extended Data Fig. 4b), while it was efficiently removed by treatment with human ARH1 (Fig. 3a-d). These findings indicate that the major product(s) of the ModB-catalysed RNAylation reaction are linked as N-glycosides via arginine residues (as shown in Fig. 3a,b).

**Fig. 3.**
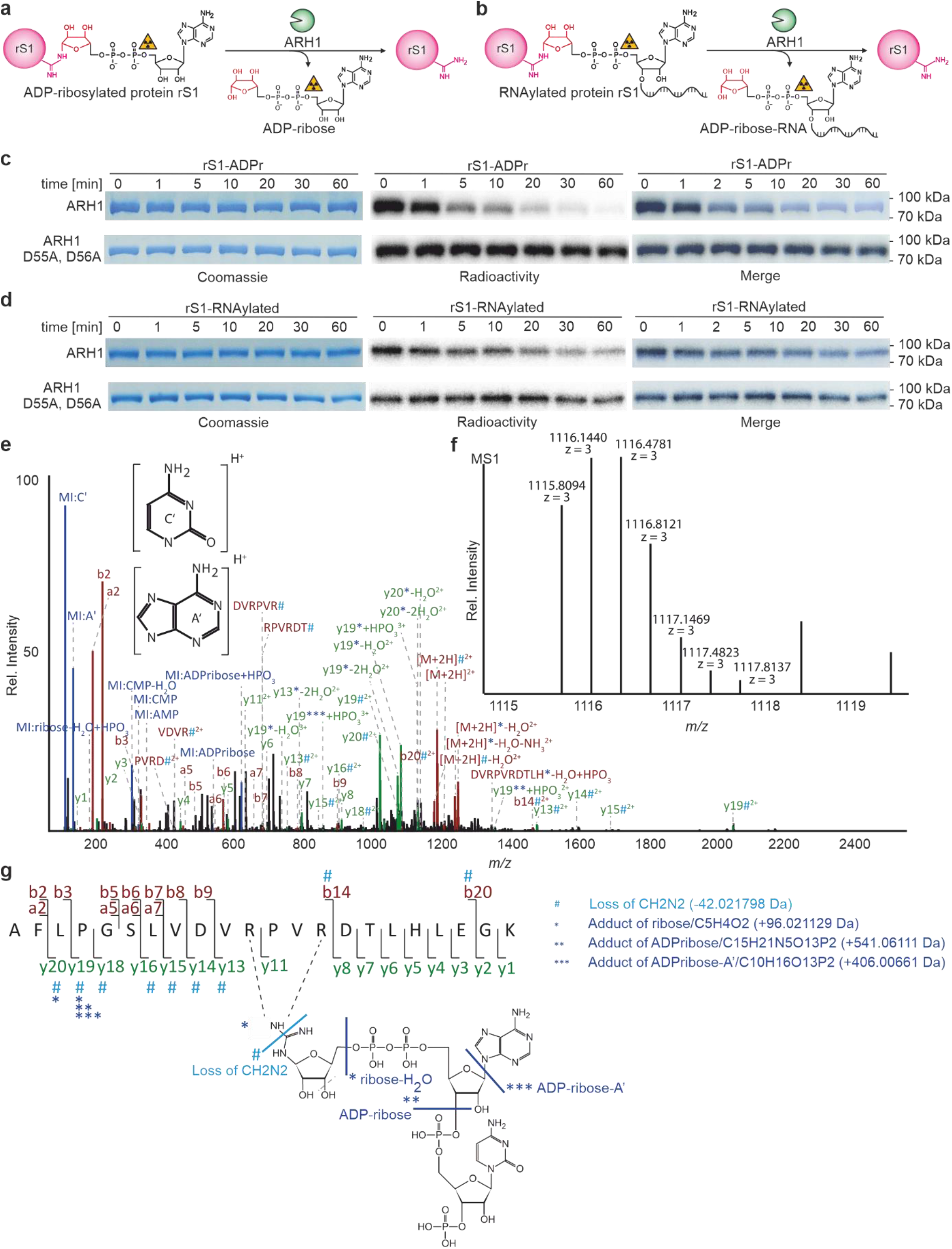
Identification of RNAylation sites of rS1. **a-d**, Specific removal of ADP-ribosylation and RNAylation by ARH1 (n=3). Enzyme kinetics of ARH1 in the presence of ADP-ribosylated or RNAylated protein rS1 analysed by SDS-PAGE. Mutation of catalytically important residues D55 and D56 abolished the removal of ADP-ribosylation and RNAylation. **e-g**, Tandem MS-based identification of RNAylated rS1 peptide. **e**, MS/MS fragment ion spectrum (spectrum ID: 23723) of RNAylated rS1 peptide AFLPGSLVDVRPVRDTLHLEGK carrying ADP-ribose plus cytidine-monophosphate and a 3’-phosphate group. The spectrum shows marker ions of adenine (A’) and cytosine (C’), AMP, CMP as well as ribose-H_2_O and ADP-ribose. The precursor ion ([M+2H]^2+^) and fragment ions y13 to y16, y18 to y20, b14 and b20 show a specific loss of 42.021798 Da (#), which can be explained by a loss of CH_2_N_2_ at the modified arginine ^31^. Precursor ions y13, y19 and y20 are detected as shifted by the mass of a ribose-H_2_O (*). The spectrum further contains precursor ions and y19 shifted by ADP-ribose with and without adenine loss (**,***). **f**, Isotopic peak pattern of the precursor ion as detected in the MS precursor ion scan for the MS/MS spectrum shown in **e**. **g**, Schematic sequence and RNA adduct representation of the RNAylated peptide shown in **e** and **f** including annotations of unshifted fragment ions and fragment ions showing arginine-losses (#) as well as ribose-H_2_O (*), ADP-ribose (**) and ADP-ribose-adenine (***). The fragmentation products of the ADP-ribose+CMP+3’-phosphate adduct observed in the MS/MS spectrum, shown in **e**, are indicated in the structure by light blue (mass loss) and dark blue (mass adducts) lines.

To establish that the ModB-mediated ADP-ribosylation or RNAylation also occurs at arginine residues *in vivo*, we isolated genomically His-tagged rS1 from non-infected or T4-infected *E. coli*. LC-MS/MS analysis confirmed specific modification of rS1 arginine residues with ADP-ribose. These ADP-ribose modifications were present only in the T4-infected sample (Extended Data Table 1, Supplementary Table 1). R139 was identified as a modified residue, which was confirmed by site-directed mutagenesis to lysine and alanine: rS1 R139K and R139A mutants were expressed in T4-infected *E. coli*, purified and analysed, revealing that these mutations prevent modification at these positions (Extended Data Table 2, Supplementary Table 2).

### LC-MS/MS analysis verifies the covalent nature of the RNAylation

LC-MS/MS analysis above could not unambiguously show that the modification of rS1 is derived from RNAylated or ADP-ribosylated rS1. Therefore, LC-MS/MS was optimized to detect the covalent attachment of an RNA to rS1. For this, *in vitro* RNAylated, truncated rS1 protein was subjected to RNase A/T1 and tryptic digest. The obtained mixture was directly subjected, to LC-MS/MS analysis and MS data were evaluated with RNPXL software tool ^30^ assuming that RNAylated rS1 peptide contains a tri-nucleotide (ADPr-cytidine) still attached. LC-MS/MS analysis showed indeed the covalent attachment of a tri-nucleotide (ADPr-cytidine) to an rS1 peptide encompassing amino acid positions 129 – 150. Strikingly, the precursor mass ([M+3H]^3+^ *m/z* = 1115.81, MW_exp_ = 3344.41 Da) plus the gas phase b- and y-type fragmentation pattern, which shows characteristic neutral loss of CH_2_N_2_ (derived from a modified arginine ^31^), or ribose, ADP-ribose, ADP-ribose-A’ adducts, revealed that the RNA is attached via an N-glycosidic bond to R139 and/or R142 (Fig. 3 e-g, Extended Data Fig. 5 and Supplementary Table 3). We note that we cannot unambiguously assign the modified arginine because of low intensities of respective fragment ions and the occurrence of mixed spectra harbouring fragment ions of the same peptide species modified at different sites (Fig. 3e-g).

### 30 % of modified rS1 are RNAylated *in vivo*

To quantitatively distinguish between ADP-ribosylation and RNAylation *in vivo*, we applied immune blotting with an antibody-like ADP-ribose binding reagent (“pan-ADPr”), which specifically recognises ADP-ribosylated rS1 and ModB, however, detects RNAylated rS1 and ModB only after nuclease P1 treatment (Fig. 4a and Extended Data Fig. 6a,b). rS1 was expressed in non-infected or T4-infected *E. coli*, affinity-purified, and its ADP-ribosylation was analysed with pan-ADPr. We found extensive ADP-ribosylation of rS1 only in the T4-infected sample. After nuclease P1 treatment, the pan-ADPr signal intensity of the rS1 band increased (Fig. 4b), indicating that ~30 % of modified rS1 was RNAylated *in vivo*. Moreover, the signal for ADP-ribose disappeared upon ARH1 treatment, again confirming the nature of the RNA-protein linkage (Extended Data Fig. 6b). We note that ADP-ribosylation and RNAylation of rS1 occur in parallel *in vivo*.

**Fig. 4.**
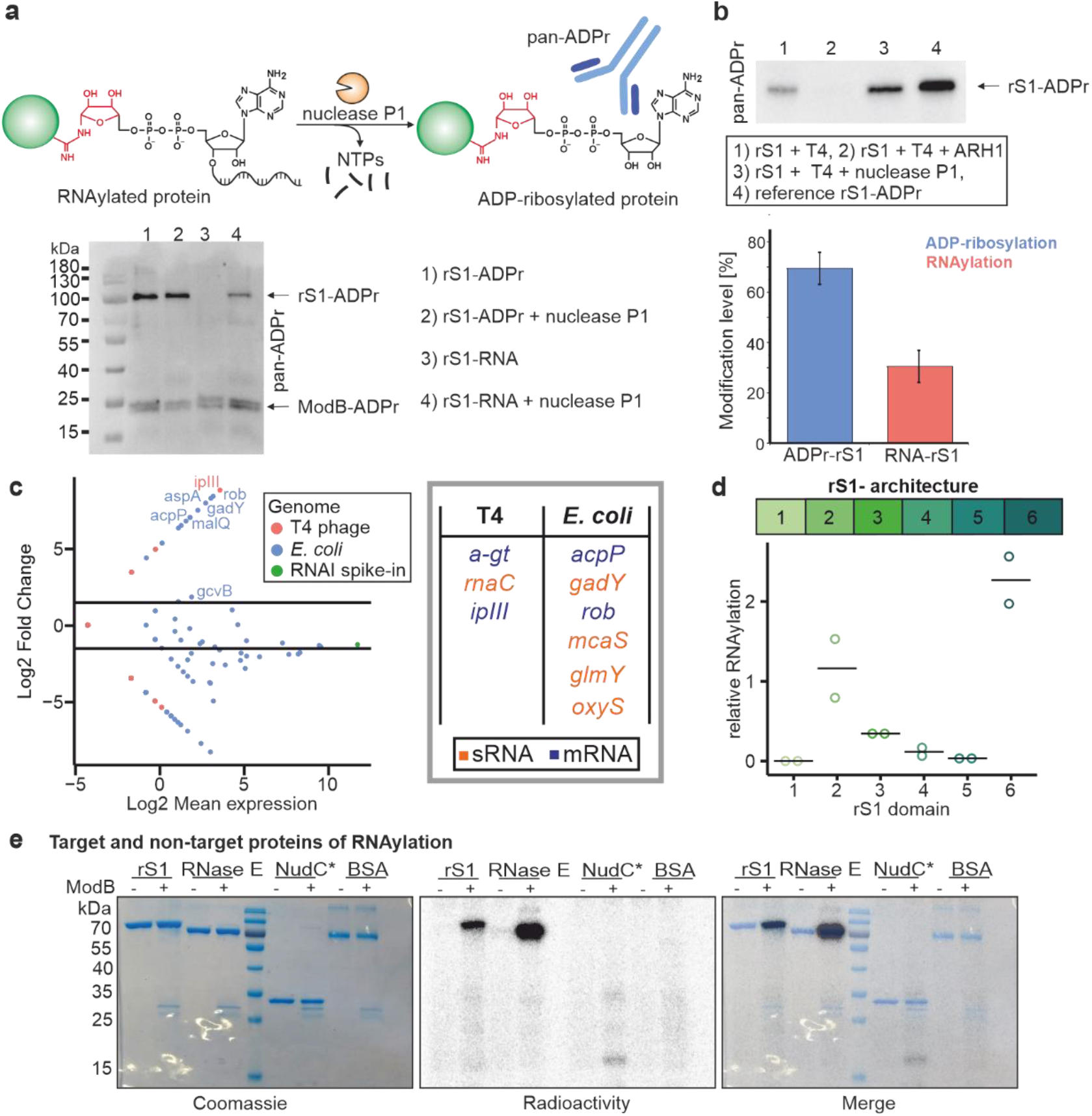
*in vivo* characterisation of ADP-ribosylation and RNAylation. **a**, Illustration of the quantification of protein rS1 RNAylation using a nuclease P1 digest and Western blot analysis. **b**, Quantification of rS1 RNAylation *in vivo* based on biological triplicates (n=3). Data is represented as mean ± s.d.. The complete corresponding blot is shown in Extended Data Fig. 6b. **c**, Identification of RNA substrates of ModB using RNAylomeSeq. MA plot shows data for replicate 1 of three biological replicates in total (n=3). **d**, Quantification of RNAylation or rS1. Modification of rS1 domains 1 – 6. n = 2 of biologically independent replicates, black lines indicate mean. **e**, SDS-PAGE analysis of the RNAylation of protein rS1, RNase E, inactive NudC mutant (NudC*: V157A, E174A, E177A, E178A) and BSA by ModB. n = 2 of biologically independent replicates.

### ModB RNAylates proteins with host and phage transcripts

To identify the RNAs linked to rS1 by ModB during T4 phage infection, we developed an RNAylomeSeq approach (Extended Data Fig. 6c), in which genomically His-tagged rS1 was isolated from T4-infected *E. coli* and captured on Ni-NTA beads. Similar to NAD captureSeq ^32^, RNA was reverse-transcribed “on-bead” and the resulting cDNA PCR-amplified and analysed by next-generation sequencing (NGS).

We applied this workflow to *E. coli* treated with T4 phage wild-type (WT). As a negative control, we generated a T4 phage that expresses the catalytically inactive mutant ModB R73A, G74A using the CRISPR-Cas9 technology ^33^. We compared the abundance of reads mapped to individual RNA species and identified specific *E. coli* and T4 phage RNAs enriched in T4 phage WT samples (Fig. 4c, Extended Data Fig. 6d,e, Supplementary Table 4). Several of the *E. coli* transcripts (mRNAs and sRNAs) have been reported to be 5’-NAD-capped in *E. coli*^3,34^; e. g., RNAs of genes *acpP, glmY, mcaS, oxyS, aspA* and *rob*, which makes them suitable substrates for ModB. We also identified phage transcripts, such as *ipIII* (internal head protein III), as enriched in our data sets (Fig. 4c, Extended Data Fig. 6d,e, Supplementary Table 4). The enriched RNAs do not share any common features except for adenosine (+1A) at the transcription start site, which is crucial for the biosynthesis of NAD-capped RNAs *in vivo* ^35^.

### ModB RNAylates OB-fold proteins

To understand how ModB identifies its target proteins we analysed the structural features of known target proteins. rS1 contains oligonucleotide-binding (OB)-fold domains ^28^. One structural variant of OB-folds is the S1 domain, present in rS1 in six copies that vary in sequence (Extended Data Fig. 7a); the RNAylated R139 and R142 are located in the rS1 domain 2. We speculated that the S1 domain might be important for substrate recognition by ModB. To characterise ModB’s specificity for different S1 domains, we cloned, expressed and purified each S1 domain of rS1 (D1-D6) and tested them in the RNAylation assay (Fig. 4d and Extended Data Fig. 7b). In agreement with the MS data (Extended Data Table 1, Supplementary Table 1) we detected strong RNAylation signals for rS1 D2 and D6, whereas rS1 D1, D3, D4 and D5 were modified to a much lesser extent. Multiple sequence alignment of rS1 D2, D6 and the S1 domain of *E. coli* PNPase revealed that these S1 domains share an arginine residue as part of the loop connecting strands 3 and 4 of the β-barrel ^36^ (Extended Data Fig. 7c). This loop is packed on the top of the β-barrel, thereby possibly accessible for ModB. For rS1 D2, particular residues are R139 and R142 which are the actual sites of RNAylation as identified by MS (Fig. 3e-g, Supplementary Table 1-3). Mutation analysis confirmed that the RNAylation level of D2 is significantly reduced if R139 is replaced by alanine or lysine (Extended Data Fig. 8a,b). *E. coli* RNase E also harbours an S1 domain in its active site with an arginine in the loop between strands 3 and 4. In the RNAylation *in vitro* assays, RNase E was modified by ModB, while control proteins without S1 domain (BSA, NudC inactive mutant) were not. These data suggest OB-folds such as S1 domains with an embedded arginine as RNAylation target motifs (Fig. 4e).

### Ribosomal protein L2 (rL2) is a novel target for RNAylation by ModB

To discover additional RNAylation target proteins of ModB, a cell lysate, prepared from exponentially growing *E. coli* was incubated with purified ModB and an NAD-10mer-RNA with a fluorescent 3’-Cy5 label (Fig. 5a, Extended Data Fig. 8c). Thereby, we thought to mimic the cellular conditions with respect to the presence of proteins, nucleic acids, and various small molecules, including NAD ^37^.

**Fig. 5.**
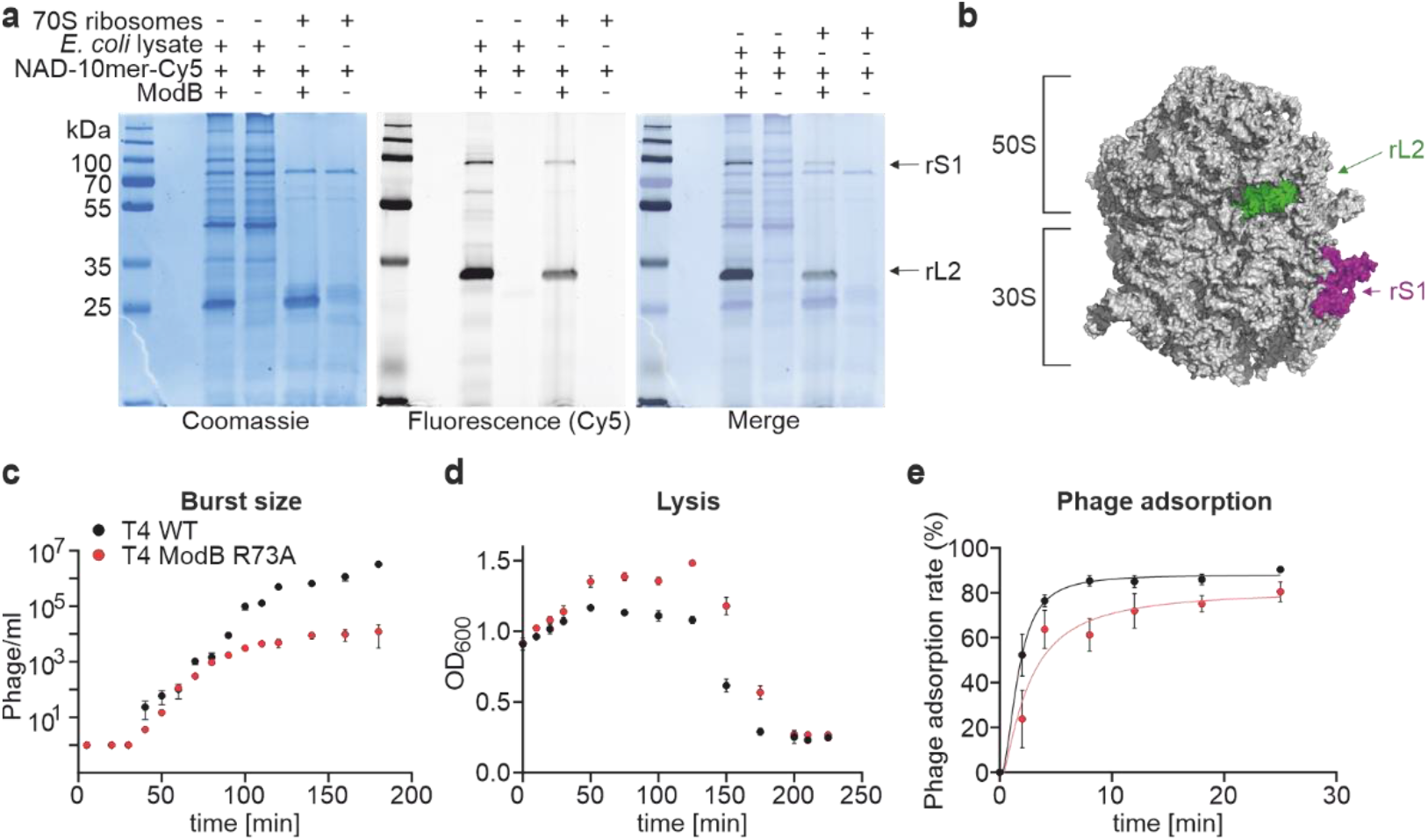
RNAylation of the ribosome and phenotype of a ModB mutant T4 phage. **a**, Characterisation of ModB substrate specificity. RNAylation of ribosomal proteins (rS1 and rL2) in cell lysates or the 70S ribosome assemblies (n=3). **b**, Illustration of the RNAylated proteins rS1 and rL2 in the context of the 70S ribosome based on the cryo-EM structure of the hibernating 70S *E. coli* ribosome (PDB: 6H4N) ^50^. **c-e**, Characterisation of the T4 ModB R73A, G74A mutant phenotype. Calculation of the burst size (**c**), *E. coli* lysis (**d**) and phage absorption (**e**) of T4 phage WT and T4 ModB R73A, G74A (n=3).

The kinetic analysis of the ModB activity in these lysates showed several *E. coli* proteins to be RNAylated (Extended Data Fig. 8c, 9a) including rS1 (which migrates similar to an RNAylated rS1 added as marker) and a protein of ~ 35 kDa size. Importantly, this pattern was not observed in the presence of 5’-monophosphorylated-RNA-Cy5. In addition, we characterized the ADP-ribosylation in the same lysates showing different patterns of ADP-ribosylation targets and RNAylation targets of ModB (Extended Data Fig. 9b). These results indicate that under cellular conditions where NAD is much more abundant than NAD-RNA, ModB RNAylates specific target proteins (Extended Data Fig. 8c, 9b). As ModB was previously assumed to preferentially ADP-ribosylate proteins involved in translation ^1^, we monitored RNAylation patterns of isolated *E. coli* ribosomes (Fig. 5a) and observed a similar pattern as in the lysates (Extended Data Fig. 8c, 9).

To identify the RNAylated proteins, we RNAylated the *E. coli* ribosome with a 40nt long NAD-RNA, resulting in a gel shift of RNAylated ribosomal proteins. Mass-spectrometric analysis of the isolated gel band identified rL2 as novel target for RNAylation by ModB (Extended Data Fig. 10a,b). rL2 is evolutionarily highly conserved, described as absolutely required for the association of 30S and 50S subunits, involved in tRNA binding to both A and P sites and important for the peptidyltransferase activity ^38^. Similar to rS1, PNPase, and RNase E, rL2 contains an RNA-binding domain which is homologous to the OB-fold ^39^. In *in vitro* RNAylation assays, ^~^80 % of rL2 were RNAylated by ModB in the presence of NAD-RNA (Extended Data Fig. 10c). *In vitro* RNAylation sites of rL2 were identified using the LC-MS/MS approach including MS data search with RNPxl as described above. Trinucleotides (ADPr-C) were found to be attached to R217 and R221 (Extended Data Fig. 10d-g, Supplementary Table 5). R221 is in close distance (11 Å) to H229, which is indispensable for the ribosomal peptidyltransferase activity ^38^. Future studies will reveal if the RNAylation of rL2 and rS1 influences the translation efficiency of the ribosome (Fig. 5b).

### ModB is important for efficient phage infection

To investigate the functional role of ModB during phage infection, we compared the phenotypes of T4 WT and T4 ModB R73A, G74A. We observed that the burst size of T4 ModB R73A, G74A was 4-fold reduced (15 progeny/cell, 50 min post-infection) in comparison to the T4 WT (60 progeny/cell, 50 min post-infection) (Fig. 5c). T4 WT produced 3.2 x 10^6^ progenies after 180 min of infection. In contrast, the infection of *E. coli* by T4 ModB R73A, G74A phage resulted in 2.4 x 10^4^ phages after 180 min. This is a 134-fold decrease in the progeny number compared to the T4 phage WT. Thus, ModB inactivation significantly affects phage propagation properties.

We also observed a delay in the lysis of approximately 20 min for the *E. coli* culture grown in the presence of the mutant phage (Fig. 5d). To determine whether ModB affects the infection cycle at the intra- or extracellular stage of infection, we measured the kinetics of phage adsorption to the cell (Fig. 5e). Here, we observed a significantly lower adsorption rate for the mutant phage. At 5 min post-infection, around 20 % of the T4 ModB R73A, G74A mutants successfully entered *E. coli*, compared to 50 % for WT T4 phage. As a result, ModB inactivation restrains the phage by hindering the first stages of the infection, namely the attachment to the host or its penetration. This finding is consistent with the delayed host lysis.

## DISCUSSION

Whereas the majority of RNA-protein interactions in biology is non-covalent ^40^, a few relevant exceptions exist ^41^, most notably the peptidyl-tRNA intermediates in protein biosynthesis ^42^ (which are esters), and the adenoviral VPg proteins that form a phosphodiester bond (via a tyrosine OH group) with a nucleotide that is then used to initiate transcription ^43,44^. Here, we show that an ADP-ribosyltransferase can attach NAD-capped RNAs to target proteins post-transcriptionally by glycosidic bond formation. This finding represents a distinct biological function of the NAD-cap on RNAs in bacteria, namely activation of the RNA for enzymatic transfer to an acceptor protein. We discovered that RNAylation of target proteins, a novel post-translational protein modification, plays a role in the infection of the bacterium *E. coli* by bacteriophage T4. We discovered that ModB is a target-specific ART that RNAylates proteins that are part of the translational apparatus. We identified rS1 and rL2 to be RNAylated at specific arginine residues in their RNA-binding regions. Moreover, we identified predominantly *E. coli* transcripts to be linked to rS1 during T4 phage infection.

These findings demonstrate the importance of the RNAylation reaction for bacteriophage pathogenicity and introduce a novel molecular mechanism by which the T4 phage controls its host’s translational machinery. Inactivation of ModB caused a significant delay in bacterial lysis upon phage infection and decreased the amount of released progeny.

ModB was known as an enzyme that uses NAD as substrate to ADP-ribosylate host proteins during T4 infection. During this study, it became evident that in addition to NAD, ModB accepts NAD-RNA as a substrate. In general, enzymes have evolved high specificity towards their substrates and tolerate only limited chemical modifications. Surprisingly, ModB tolerates the attachment of a bulky RNA chain to the 3’-OH-group of NAD (NAD-RNA) for modification of a specific subset of target proteins. Remarkably, all four proteins (rS1, rL2, RNase E, PNPase) identified here as RNAylation targets of ModB are well-known to interact with RNA. We therefore assume that both the ability of ModB to accept NAD-RNA as a substrate and the RNA-affinity of the target protein determine RNAylation specificity. Why does a phage ART attach specific RNAs to proteins involved in translation? Upon infection of *E. coli*, T4 phage aims to reprogram the host ribosome to translate its mRNAs ^45^. One possibility to achieve this may be a controlled shutdown of ribosomes that do not participate in the translation of T4 mRNAs. The discovery of crucial ribosomal proteins, rS1 and rL2, as RNAylation targets let us speculate that RNAylation impairs their functionality, e.g., modulating peptidyltransferase activity. The fact that mostly *E. coli* transcripts are linked to rS1 *in vivo* suggests that undesired host gene expression events are stopped by RNAylation. Thus the phage might exploit RNAylation to inactivate distinct ribosomes. Future studies will show if ribosomes that translate *E. coli* transcripts are blocked by RNAylation. This proposed mechanism would enable the phage to regulate the activity of the ribosome throughout the infection cycle and to stop the translation of host proteins.

Why only one out of three known T4 ARTs carries out efficient RNAylations is not understood. ModA and ModB both contain characteristic features of arginine-specific ADP-ribosyltransferases, such as the active site motif R-S-EXE ^1^. Differences in substrate specificity are therefore likely due to sequence differences (25% identical, 47% homologous amino acids between ModA and ModB) ^1^.

ARTs are known to occur not only in bacteriophages, and ADP-ribosylated proteins have been detected in hosts upon infections by various viruses, including influenza, corona and HIV. In addition to viruses using ARTs as weapons, the mammalian antiviral defence system applies host ARTs to inactivate viral proteins. Moreover, mammalian ARTs and poly-(ADP-ribose) polymerases (PARPs) are regulators of critical cellular pathways and are known to interact with RNA ^46^. Thus, ARTs in different organisms might catalyse RNAylation reactions, and RNAylation may be a phenomenon of broad biological relevance.

Finally, RNAylation may be considered as both a post-translational protein modification and a post-transcriptional RNA modification. Our findings challenge the established views of how RNAs and proteins can interact with each other. The discovery of these new RNA-protein conjugates comes at a time when the structural and functional boundaries between the different classes of biopolymers become increasingly blurry ^47,48^.

## ACKNOWLEDGEMENTS

We thank N. Beumer, J. Hoff, S. J. Keding, J. Kahnt, J. Koch, P. Mann, N. Moskalchuk, M. Raabe, E. Tamerler, M. Viering, and M. Weber for experimental assistance. This project has received funding from the European Research Council (ERC) under the European Union’s Horizon 2020 research and innovation programme (Grant agreement No. 882789 RNACoenzyme, to A.J.) and by the German Research Council (DFG, Project-ID 439669440, TRR319, project A02, to A.J.). M.W. is supported by the Studienstiftung des Deutschen Volkes e.V. and the Joachim Herz Stiftung. K.H. is supported by the Max Planck Society, Baden-Württemberg Stiftung, Carl-Zeiss-Stiftung and a grant of the German Research Council (DFG-SPP2330). H.U. is supported by the Max Planck Institute for Multidisciplinary Sciences and by grants of the Deutsche Forschungsgemeinschaft (DFG-SPP1935, DFG-SFB-1286, DFG-SFB-1565).

## AUTHOR CONTRIBUTIONS

K.H. and A.J. designed the study. K.H., M.W., J.G., F.A.B., N.P. cloned, expressed, purified and analysed ARTs and their target proteins. K.H., I.S., L.M.W., A.W. and M.W. prepared samples for mass spectrometry. I.S., L.M.W., A.W. and H.U. developed an LC-MS/MS pipeline to study ADP-ribosylation/RNAylation and analysed the data. T.G. performed mass spectrometric analysis of rS1. M.W. developed the RNAylomeSeq pipeline and analysed the data. N.P. created and characterised the ModB mutant phage. K.H., H.U. and A.J. supervised the work. K.H. and A.J. wrote the first draft, and all authors contributed to reviewing, editing and providing additional text for the manuscript.

## MATERIALS & CORRESPONDENCE

Supplementary Information is available for this paper.

The datasets generated during and/or analysed during the current study are available from the corresponding author on reasonable request. NGS data is accessible via GEO record GSE214431 (reviewer token: evedsmayhvyxfen). LC-MS/MS raw data for measurements of rS1 ADP-ribosylation *in vivo* have been deposited in PRIDE under the accession PXD038075 (reviewer access via username; password: Uz3xrs6b). LC-MS/MS raw data for measurements of *in vitro* ADP-ribosylated and RNAylated rS1 and rL2 have been deposited in PRIDE under the accession PXD038910 (reviewer access via username; password: Vt8nPXs3).

Correspondence should be addressed to A.J. and K.H.

## METHODS

### General

Reagents were purchased from Sigma-Aldrich and used without further purification. Oligonucleotides, DNA and RNA, were purchased from Integrated DNA Technologies, Inc. (Supplementary Table 6 - 9). DNA and RNA concentrations were determined by measurements with the NanoDrop ND-1000 spectrophotometer. Radioactively labelled proteins or nucleic acids were visualised using storage phosphor screens (GE Healthcare) and a Typhoon 9400 imager (GE Healthcare).

### Preparation of 5’PPP-/5’P-/5’-NAD-RNA by *in vitro* transcription

DNA templates for Qβ-RNA (100nt-RNA) and RNAI were amplified by PCR (primer sequences are listed Supplementary Table 8) and PCR products were analysed by 2 % agarose gel electrophoresis and purified using the QIAquick PCR purification kit (QIAGEN). 5’-triphosphate (PPP) Qβ-RNA and RNAI were synthesised by *in vitro* transcription (IVT) in the presence of 1 x transcription buffer (40 mM Tris, pH 8.1, 1 mM spermidine, 10 mM MgCl_2_, 0.01 % Triton-X-100), 5 % DMSO, 10 mM DTT, 4 mM of each NTP, 20 μg of T7 RNA polymerase (2 mg/ml, self-purified) and 200 nM DNA template. NAD-RNAI was generated under similar conditions using only 2 mM ATP and 4 mM NAD. The same conditions were applied for the synthesis of a mixture of α-^32^P-labelled 5’-NAD- and PPP-Qβ-RNAs except for the presence of 2 mM ATP, 80 μCi ^32^P-α-ATP and 4 mM NAD instead of 4 mM ATP. The IVT reactions were incubated at 37 °C for 4 h and digested with DNase I (Roche). RNA was purified by denaturing PAGE, isopropanol-precipitated and resuspended in Millipore water. RNA sequences are listed in Supplementary Table 6.

To convert 5’-PPP-RNAs into 5’-monophosphate RNAs (5’-P-RNAs), 250 pmol of Qβ-RNA were treated with 60 U of RNA 5’-polyphosphatase (Epicentre) in 1 x polyphosphatase reaction buffer at 37 °C for 70 min. Protein was removed from 5’-P-RNAs by phenol-chloroform extraction and residual phenol-chloroform removed by three rounds of diethyl ether extraction. 5’-P-RNAs were isopropanol precipitated and resuspended in Millipore water.

### 5’-radiolabelling of 5’-monophosphate and NAD-capped RNAs

120 pmol of 5’-P-Qβ-RNA or 6.25 nmol of 5’-P-RNA 8mer (Supplementary Table 6) were treated with 50 U of T4 polynucleotide kinase (PNK) in 1 x reaction buffer B and 1,250 μCi ^32^P-γ-ATP. The reaction was incubated at 37 °C for 2 h. The resulting 5’-^32^P-RNA 8mer/ 5’-^32^P-Qβ-RNA were separated from residual protein by phenol-chloroform extraction. The remaining ^32^P-γ-ATP was removed by washing with 3 column volumes of Millipore water and centrifugation in 10 kDa (Qβ-RNA) or 3 kDa (8mer) Amicon filters (Merck Millipore) at 14,000 rpm at 4 °C for four times. RNA sequences are listed in Supplementary Table 6. To convert the purified 5’-^32^P-RNAs into 5’-^32^P-NAD-capped RNAs, 800 pmol of 5’-^32^P-RNA 8mer or 30 pmol of 5’-^32^P-Qβ-RNA were each incubated in 50 mM MgCl_2_ in the presence of a spatula tip of nicotinamide mononucleotide phosphorimidazolide (Im-NMN), synthesised as described in ^1^, at 50 °C for 2 hours. RNAs were purified by washing with Millipore water and centrifugation in 10 kDa (Qβ-RNAs) or 3 kDa (8mer) Amicon filters at 14,000 rpm at 4 °C for four times. Concentrations of the 5’-^32^P-RNAs were measured on the NanoDrop ND-1000 spectrophotometer and used to calculate the approximate concentrations of yielded 5’-NAD-capped ^32^P-RNAs assuming an approximate yield of the imidazolide reaction of 50 % ^1^. 5’-^32^P-ADP-ribose-RNA 8mer (ADPr-8mer) was synthesised by incubation of 4.8 μM 5’-^32^P-NAD-RNA 8mer and 0.08 μM ADP-ribosyl cyclase CD38 (R & D Systems) in 1 x degradation buffer at 37 °C for 4 h. The reaction was purified by P/C/I-diethyl ether extraction and filtration through 3 kDa filters washing with 4 column volumes of Millipore water.

### Cloning of ADP-ribosyltransferases, ADP-ribose hydrolases and target proteins

To amplify bacteriophage T4 genes *modA, modB* and *alt*, a single plaque from bacteriophage T4 revitalisation was resuspended in Millipore water and used in a “plaque” PCR, analogous to bacterial colony PCR. The gene encoding for the ADP-ribosylhydrolase ARH1 was purchased from IDT as gblocks and amplified by PCR. *E. coli* genes coding for rS1 and PNPase were PCR-amplified from genomic DNA of *E. coli* K12, which was isolated via GenElute Bacterial Genomic DNA Kit (Sigma-Aldrich). Nucleotide sequences are listed in Supplementary Table 7. XhoI and NcoI restriction sites were introduced during amplification using appropriate primers (Supplementary Table 8). The resulting PCR product was digested with XhoI and NcoI (Thermo Fisher Scientific) and cloned into the pET-28c vector (Merck Millipore). After Sanger sequencing, the resulting plasmids were transformed into *E*. *coli* One Shot BL21 (DE3) (Life Technologies). The ARH1 D55,56A, ModB R73A and rS1 mutants were generated by site-directed mutagenesis using a procedure based on the Phusion Site-Directed Mutagenesis Kit (Thermo Scientific). The resulting plasmids were sequenced and transformed into *E*. *coli* One Shot BL21 (DE3). All strains used and generated in this work are summarised in Supplementary Table 9.

### Purification of rS1, rS1 domains and variants, rL2, PNPase S1 domain, RNase E (1-529), Alt, NudC, NudC* (V157A, E174A, E177A, E178A) and NudC E178Q

IPTG-induced *E. coli* One Shot BL21 (DE3) containing the respective plasmid (Supplementary Table 9) were cultured in LB medium at 37 °C. Protein expression was induced at OD_600_ = 0.8, bacteria were harvested by centrifugation after 3 hours at 37 °C and lysed by sonication (30 s, 50 % power, five times) in HisTrap buffer A (50 mM Tris-HCl pH 7.8, 1 M NaCl, 1 M urea, 5 mM MgSO_4_, 5 mM β-mercaptoethanol, 5 % glycerol, 5 mM imidazole, 1 tablet per 500 ml complete EDTA-free protease inhibitor cocktail (Roche)). The lysate was clarified by centrifugation (37,500 g, 30 min, 4 °C) and the supernatant was applied to a 1 ml Ni-NTA HisTrap column (GE Healthcare). The protein was eluted with an imidazole gradient using an analogous gradient of HisTrap buffer B (HisTrap buffer A with 500 mM imidazole) and analysed by SDS–PAGE.

Further protein purification was achieved by size exclusion chromatography (SEC) through a Superdex™ 200 10/300 GL column (GE Healthcare) using SEC buffer containing 0.5 M NaCl and 25 mM Tris-HCl, pH 8. Fractions of interest were analysed by SDS–PAGE, pooled and concentrated in Amicon Ultra-4 centrifugal filters (MWCO 10 kDa, centrifugation at 2,000 rpm, 4 °C). Protein concentration was measured with the NanoDrop ND-1000 spectrophotometer. Proteins were finally stored in SEC buffer supplemented with 50 % glycerol at −20 °C.

### Purification of ARH1 and ARH1 D55A, D56A

*E. coli* BL21 DE3 pET28-ARH1 and BL21-pET28-ARH1 D55A, D56A (Supplementary Table 9) were grown to an OD_600_ = 0.6 at 37 °C, 175 rpm. Afterwards, bacteria were allowed to cool to room temperature for 30 minutes. Expression was induced with 1 mM IPTG, and bacteria were finally grown o/n at room temperature, 175 rpm. Bacteria were harvested by centrifugation, and proteins were purified analogously to rS1 variants.

### Purification of ModA

*E. coli* BL21 DE3 pET28-ModA (Supplementary Table 9) were grown to an OD_600_ = 1 at 37 °C, 175 rpm. Protein expression was induced with 0.5 mM IPTG and bacteria were harvested by centrifugation after 3 hours at 37 °C. Pelleted bacteria were resuspended in 50 mM NaH_2_PO_4_, pH 8, 300 mM NaCl, 1 mM DTT, 1 tablet per 500 ml complete EDTA-free protease inhibitor cocktail (Roche) and lysed by sonication (3 x 1 min, 50 % power). Lysates were centrifuged at 3,000 *g*, 4 °C for 20 min. Sediments were washed by resuspension in 30 ml 50 mM Tris-HCl, pH 7.5, 2 mM EDTA, 100 mM NaCl, 1 M urea, 1 mM DTT, 1 tablet EDTA-free protease inhibitor (Roche), and centrifuged at 10,000 *g*, 4 °C for 20 min. Pellets, containing inclusion bodies, were resuspended in 40 ml 100 mM Tris pH 11.6, 8 M urea, transferred to 12-14 kDa MWCO dialysis bags (Roth), and dialysed o/n against 50 mM NaH_2_PO_4_, 300 mM NaCl. Protein solutions were centrifuged at 20,000 *g*, 4 °C for 30 min. Supernatants were batch-purified using disposable 10 ml columns (Thermo Fisher Scientific) packed with 2 ml Ni-NTA agarose (Jena Bioscience) and equilibrated with 10 column volumes (CV) of 50 mM NaH_2_PO_4_ (pH 8), 300 mM NaCl. Proteins were purified by washing the columns with 30 CV 50 mM NaH_2_PO_4_, 300 mM NaCl, 15 mM imidazole, eluted with 5 ml 50 mM NaH_2_PO_4_, 300 mM NaCl, 300 mM imidazole, and concentrated in Amicon (Merck Millipore) filters (MWCO 10 kDa, centrifugation at 2,000 rpm, 4 °C). Proteins were finally purified by SEC as described for rS1.

### Purification of ModB and ModB R73A G74A

*E. coli* BL21 DE3 pET28-ModB or *E. coli* BL21 DE3 pET28-ModB R73A G74A (Supplementary Table 9) were grown to an OD_600_ = 2.0 at 37 °C, 185 rpm and cooled down to 4 °C while shaking at 160 rpm for at least 30 min. Protein expression was induced by the addition of 1 mM IPTG. The cultures were then incubated for 120 min at 4 °C, 160 rpm and bacteria harvested by centrifugation (4,000 rpm, 4 °C, 25 min). The ModB protein was purified from the supernatant as described for rS1 variants.

### *In vitro* ADP-ribosylation and RNAylation of rS1 and rL2 with ^32^P-labelled NAD, NAD-8mer, NAD-Qβ-RNA or NAD-10mer-Cy5

0.3 μM protein rS1 were ADP-ribosylated in the presence of 0.25 μCi/μl ^32^P-NAD or RNAylated in the presence of either 0.6 μM ^32^P-NAD-8mer, 0.03 μM ^32^P-NAD-Qβ-RNA or 0.8 μM NAD-10mer-Cy5 (Supplementary Table 6) by 1.4 μM ModB and in 1 x transferase buffer (10 mM Mg(OAc)_2_, 22 mM NH_4_Cl, 50 mM Tris-acetate pH 7.5, 1 mM EDTA, 10 mM ß-mercaptoethanol, 1 % glycerol) at 15 °C for at least 120 minutes. 5 μl samples were taken after 0 (before the addition of ModB), 1, 2, 5, 10, 30, 60 and 120 minutes and mixed with 5 μl 2 x Laemmli buffer to stop the reaction. Reactions were assessed by 12 % SDS-PAGE and gels stained in Instant Blue solution (Sigma-Aldrich) for 10 min. Radioactive signals were visualised using storage phosphor screens and a Typhoon 9400 imager. Intensities of radioactive bands were quantified using ImageQuant 5.2 (GE Healthcare). The RNAylation with NAD-capped Cy5-labelled RNA was visualised with the ChemiDoc (Bio-Rad), Cy5 channel. Afterwards, gels were stained in Coomassie solution and imaged with the same system.

rL2 was ADP-ribosylated or RNAylated at the same settings using either 6.4 μM NAD or 6.4 μM NAD-8mer as substrate to modify 4.6 μM rL2 in the presence of 1.57 μM ModB for either 4 h for LC-MS/MS measurements. For shift assays, 538 nM rL2 were RNAylated by 2.61 μM ModB in the presence of 6 μM NAD-8mer. 12 % SDS-PA gels were fixed with fixation solution (40 % ethanol, 10 % acetic acid) o/n and stained in Flamingo fluorescent protein dye (Bio-Rad) for up to 6 h and imaged with the ChemiDoc (Bio-Rad). Signal intensities were quantified in ImageLab (Bio-Rad).

### *In vitro* RNAylation of *E. coli* RNA polymerase with NAD-10mer-Cy5

0.8 μM NAD-10mer-Cy5 (Supplementary Table 6) were incubated with 0.5 μM of protein *E. coli* RNA polymerase (NEB) and 3 μM of Alt or ModA in the presence of 1 x transferase buffer at 15 °C for 60 minutes. Samples were taken before the addition of Alt or ModA (t0) and after 60 minutes (t60) of incubation. The reactions were stopped by the addition of 1 volume 2 x Laemmli buffer. Reactions were analysed by 10 % SDS-PAGE applying as a reference protein rS1 RNAylated by ModB with NAD-10mer-Cy5. RNAylated proteins were visualised with the ChemiDoc (Bio-Rad), Cy5 channel. Afterwards, gels were stained in Coomassie solution and imaged with the same system.

### Analysis of protein rS1 self-RNAylation

In scales of 20 μl reactions, 3.6 μM of ^32^P-ADPr-8mer (Supplementary Table 6) were either incubated with 2.6 μM of protein rS1, 3.9 μM of ModB or both 2.59 μM of protein rS1 and 3.9 μM of ModB in 1 x transferase buffer. As a positive control, equal amounts of protein rS1 and ModB were incubated with 0.6 μM of ^32^P-NAD-8mer. All reactions were incubated at 15 °C for 60 minutes. Samples were taken before the addition of ModB (0 minutes), after 60 minutes of incubation and reactions were stopped by the addition of 1 volume 2 x Laemmli buffer each. Reactions were analysed by 12 % SDS-PAGE and autoradiography imaging.

### RNAylation of protein rS1 with Qβ-RNA (100nt-RNA) and specificity for the 5’-NAD-cap

0.05 μM ^32^P-NAD-Qβ-RNA, 0.15 μM 5’-^32^P-Qβ-RNA or 0.15 μM 5’-^32^PPP-Qβ-RNA (Supplementary Table 6) were incubated with 2.3 μM of protein rS1 and 1.4 μM of ModB each in the presence of 1 x transferase buffer at 15 °C for 60 minutes. Samples were taken before the addition of ModB (0 minutes) and after 60 minutes of incubation, and reactions were stopped by adding 1 volume 2 x Laemmli buffer. Reactions were analysed by 10 % SDS-PAGE, applying rS1-^32^P-ADPr in 1 x Laemmli buffer as a reference, and subsequent autoradiography imaging.

### Preparation of RNAylated and ADP-ribosylated rS1 for enzymatic treatments

ADP-ribosylation or RNAylation reactions performed with radio-labelled substrates were washed and equilibrated in 1 x transferase or 1 x degradation buffer for further enzymatic treatments. Therefore, the reactions were washed with 4 column volumes of the corresponding buffer via centrifugation at 10,000 x g, 4 °C in 10 kDa Amicon (Merck Millipore) filters. Proteins RNAylated with Cy5-labelled RNA were equilibrated in the same buffers using Zeba Spin desalting columns (7 MWCO, 0.5 ml) (Thermo Fisher Scientific) according to manufacturer’s instructions.

### Nuclease P1 digest of protein rS1 RNAylated with 100nt-RNA (rS1-100nt-RNA)

19 μl of rS1-100nt-RNA (^32^P) mixture equilibrated in 1 x transferase buffer were incubated with either 1 μl of nuclease P1 or 1 μl of Millipore water at 37 °C for 60 minutes. Samples were taken before the addition of enzyme (t0) and after 60 minutes of incubation (t60) and reactions stopped by addition of 1 volume 2 x Laemmli buffer. Reactions were analysed by 10 % SDS-PAGE, applying rS1-^32^P-ADPr in 1 x Laemmli buffer as a reference, and subsequent autoradiography imaging.

### Tryptic digest of ^32^P-labelled rS1-8mer and rS1-ADPr

19 μl of both rS1/rS1-8mer (^32^P) mixture and rS1/rS1-ADPr (^32^P) mixture in 1 x degradation buffer were incubated with either 0.2 μg Trypsin (Sigma, EMS0004, mass spectrometry grade) or Millipore water as a negative control at 37 °C. Samples were taken before the addition of Trypsin/Millipore water (t0) and after 120 minutes of incubation (t120). Reactions were stopped by adding 1 volume 2 x Laemmli buffer to samples and were analysed by 12 % SDS-PAGE and autoradiography imaging.

### Chemical removal of ADP-ribosylation and RNAylation *in vitro*

Aliquots from washed and equilibrated ADP-ribosylated (1 μl) and RNAylated (2 μl) (^32^P) rS1 were treated with either 10 mM HgCl_2_ or 500 mM NH_2_OH ^2,3^ at 37 °C for 1h. Reactions were stopped by addition of 2 x Laemmli-Buffer and analysed by 12 % SDS-PAGE.

### Enzymatic removal of ADP-ribosylation and RNAylation *in vitro*

Aliquots from washed and equilibrated (in 1 x degradation buffer) ADP-ribosylated (1 μl) and RNAylated (2 μl) rS1 (^32^P) were treated with 0.5 U endonuclease P1 (Sigma-Aldrich) ^4^ or 0.95 μM ARH1 or ARH3 (human, recombinant, Enzo Life Science) ^5^ in the presence of 10 mM Mg(OAc)_2_, 22 mM NH_4_Cl, 50 mM HEPES, 1 mM EDTA, 10 mM ß-mercaptoethanol and 1 % (v/v) glycerol in a total volume of 20 μl at 37 °C for 1 h. Enzymatic reactions were stopped by the addition of 2 x Laemmli-Buffer and analysed by 12 % SDS-PAGE.

### Inhibition of RNAylation and ADP-ribosylation with 3-methoxybenzamide (3-MB)

20 μl reactions of 1.4 μM ModB and 2.3 μM protein rS1 with either 1 μM of ^32^P-NAD-8mer or 3 μM 5’-^32^P-8mer (Supplementary Table 6) were incubated in the presence of 2 mM 3-MB (50 mM stock in DMSO) or the absence of the inhibitor (DMSO only) at 15 °C ^6^. Samples were taken before the addition of ModB (t0) and after 60 minutes of incubation (t60) with ModB. Reactions were stopped by the addition of 1 volume 2 x Laemmli buffer and analysed by 12 % SDS-PAGE.

### Effect of RNA secondary structure on RNAylation efficiency

1.1 μM NAD-RNA-Cy5 (linear, 5’-overhang, 3’-overhang, and blunt ends, Supplementary Table 6) were incubated with 0.9 μM rS1 and 0.4 μM ModB in 1 x transferase buffer. 5 μl samples were taken before addition of ModB protein (0 min) and 2, 5, 10, 30, 60, and 120 min after the reaction start. The sample was directly mixed with 1 volume of 2 x Laemmli buffer to stop the reaction. The conversion of the substrates was analysed via 12 % SDS-PAGE, following visualisation on the ChemiDoc (Bio-Rad) in the Cy5 channel. The maximal observed signal intensity of RNAylated rS1 protein was used to determine the relative conversion for each of the analysed substrates at distinct time points.

### Cultivation of *E. coli* B strain and T4 phage infection

Pre-cultures of *E. coli* B strain pTAC-rS1 (Supplementary Table 9) were incubated in LB medium with 100 μg/ml ampicillin at 37 °C, 185 rpm o/n. For main cultures, 150 ml LB medium with 100 μg/ml ampicillin were inoculated with pre-culture to an OD_600_ = 0.1. At OD_600_ = 0.4 protein expression was induced by the addition of 1 mM IPTG. At an OD_600_ = 0.8, cultures were either infected with bacteriophage T4 at an MOI 10 (20 ml phage solution) (DSM 4505, Leibniz Institute DSMZ) or not infected by adding 20 ml LB medium instead (negative control). Cultures were incubated for 20 min at 37 °C, 240 rpm. Bacteria were harvested by centrifugation at 4,000 x g at room temperature for 15 min. Pellets were stored at −80 °C.

### Purification of His-tagged rS1 from infected *E. coli* strain B pTAC-rS1

Bacterial pellets were resuspended in 10 ml buffer A and lysed via sonication (1 x 5 min, cycle 2, 50 % power). Lysates were centrifuged at 37,500 g, 4 °C for 30 min. The supernatant was filtered through 0.45 μm filters (Sarstedt). rS1 from bacteriophage T4-infected or non-infected *E. coli* B strain was purified from the supernatant by gravity Ni-NTA affinity chromatography. 1 ml of Ni-NTA agarose slurry (Thermo Fisher Scientific) was added to a 10 ml propylene column and equilibrated in buffer A. The supernatant was loaded onto the column twice. The column was washed with a mixture of 95 % buffer A and 5 % buffer B containing 29.75 mM imidazole. Protein was eluted from the column with 10 ml buffer B.

His-tagged-protein rS1 from T4-infected or uninfected *E. coli* B strain pTAC-rS1 was washed with two filter volumes of 1 x degradation buffer (12.5 mM Tris-HCl, pH 7.5, 25 mM NaCl, 25 mM KCl, 5 mM MgCl_2_) by centrifugation in 10 kDa Amicon filters at 5,000 g, 4 °C and concentrated to a final volume of 120 μL. The fractions were analysed by 12 % SDS-PAGE analysis, and the gel was stained in Instant Blue solution for 10 min and imaged immediately.

### Purification of His-tagged rS1 and rL2 for LC-MS/MS analysis

*E. coli* B strain with endogenously His-tagged rS1 or *E. coli* B strain expressing His-tagged rS1 WT, R139A or R139K were infected with T4 an MOI of 5.0 as described above for 8 minutes. 100 ml culture were harvested and the pellet resuspended in 1.5 ml Ni-NTA buffer A with 15 mM imidazole (50 mM Tris-HCl pH 7.8, 1 M NaCl, 1 M urea, 5 mM MgSO_4_, 5 mM β-mercaptoethanol, 5 % glycerol, 15 mM imidazole, 1 tablet per 500 ml complete EDTA-free protease inhibitor cocktail (Roche)). Cells were lysed by sonication (3 times, 2 min, 80 % power) and supernatant cleared by centrifugation at 17,000 x g, 4 °C, 30 min. The supernatant was incubated with 75 μl Ni-NTA magnetic beads (Jena Bioscience) equilibrated in Ni-NTA buffer A with 15 mM imidazole for 1 h at 4 °C. Magnetic beads were washed 7 times with 1 ml Ni-NTA buffer A with 15 mM imidazole and 3 times with Ni-NTA buffer without imidazole but with 4 M urea. Finally, protein was eluted by addition of Ni-NTA elution buffer (50 mM Tris-HCl pH 7.8, 1 M NaCl, 1 M Urea, 5 mM MgSO_4_, 5 mM β-mercaptoethanol, 5 % glycerol, 300 mM imidazole, 1 tablet per 500 ml complete EDTA-free protease inhibitor cocktail (Roche)). Protein was equilibrated in 1 x transferase buffer with Zeba columns (7 MWCO, 0.5 ml) according to manufacturer’s instructions and protein was digested with trypsin in a 1:20 ratio (w/w) at 37 °C for 3 h. Peptides were C18-purified using 50 mM triethylamine-acetate (pH 7.0) buffer in combination with 0 – 90 % acetonitrile and Chromabond C18 WP spin columns (20 mg, Macherey Nagel, Germany). Purified peptides were dissolved in HPLC-grade H_2_O and subjected to LC-MS/MS analysis (see below).

*In vitro* RNAylated rS1 (D2) reactions in 1 x transferase buffer were directly digested (without further purification) with 1 μg RNase A (Thermo Fisher Scientific) and 100 U RNase T1 (Thermo Fisher Scientific) at 37 °C for 1 h, following tryptic digest at 37 °C for 3 h in the same buffer with trypsin (Promega) in a 1:30 ratio (w/w) relative to total protein content per sample. Peptides were purified with Chromabond C18 WP spin columns as described above and subjected to LC-MS/MS analysis (see below).

*In vitro* RNAylation reactions of rL2 with NAD-8mer and ADP-ribosylation reactions were purified at similar settings as for proteins from T4 phage-infected *E. coli*. Here, 200 μl reactions were incubated with 100 μl Ni-NTA beads equilibrated in 800 μl Ni-NTA buffer A with 10 mM imidazole and 40 U murine RNase inhibitor (NEB) at 4 °C for 1 h. Beads were washed 8 times with 1 ml streptavidin wash buffer (50 mM Tris-HCl pH 7.4, 8 M urea) at room temperature and protein was eluted with 130 μl Ni-NTA elution buffer. Purified proteins were re-buffered in 100 mM NH_4_OAc using Zeba spin desalting columns (7 MWCO, 0.5 ml) according to manufacturer’s instructions. rL2 samples were dissolved in 4 M urea in 50 mM Tris-HCl (pH 7.5) and incubated for 30 min at room temperature followed by dilution to 1 M urea with 50 mM Tris-HCl (pH 7.5). 10 μg RNase A (Thermo Fisher Scientific) and 1kU RNase T1 (Thermo Fisher Scientific) were added, following incubation for 4 h at 37 °C. For protein digestion, 0.5 μg of trypsin (Promega) were added to each sample and digestion was performed o/n at 37 °C. Samples were adjusted to 1 % acetonitrile (ACN) and to pH 3 using formic acid (FA). Sample cleanup was performed using C18 columns (Harvard Apparatus) according to the manufacturer’s instructions.

### LC-MS/MS analysis of His-tagged, *in vitro* RNAylated rS1 and rL2

Cleaned-up rS1 and rL2 peptide samples were dissolved in 2 % ACN, 0.05% trifluoroacetic acid (TFA) and subjected to LC-MS/MS analysis using an Orbitrap Exploris 480 Mass Spectrometer (Thermo Fisher Scientific) coupled to a Dionex Ultimate 3000 RSLCnano system. Peptides were loaded on a Pepmap 300 C18 trap column (Thermo Fisher Scientific) (flow rate, 10 μl/min) in buffer A (0.1 % [v/v] FA) and washed for 3 min with buffer A. Peptide separation was performed on an in-house packed C18 column (30 cm; ReproSil-Pur 120Å, 1.9 μm, C18-AQ; inner diameter, 75 μm; flow rate: 300 nl/min) by applying a linear gradient of buffer B (80% [v/v] ACN, 0.08% [v/v] FA). The main column was equilibrated with 5 % buffer B for 18 s, sample was applied and column was washed for 3 min with 5% buffer B.

A linear gradient from 10-45 % buffer B over 44 min was applied to elute peptides, followed by 4.8 min washing at 90 % buffer B and 6 min at 5% buffer B. Eluting rS1 and rL2 peptides were analysed for 58 min in positive mode using a data-dependent top 20 acquisition method. MS1 and MS2 resolution were set to 120,000 and 30,000 FWHM, respectively, and AGC targets were set to 10^6^ (MS1) and 10^5^(MS2). MS1 scan range was set to m/z 350-1600. Precursors were fragmented using 28% normalised, higher-energy collision-induced dissociation (HCD) fragmentation. Other analysis parameters were set as follows: isolation width, 1.6 m/z; dynamic exclusion, 9 s; max. injection times (MS1/MS2), 60 ms/120 ms.

For all measurements, the lock mass option (m/z 445.120025) was used for internal calibration.

### Analysis of *in vitro* RNAylated rS1 and rL2 MS data

MS data were analysed and manually validated using the OpenMS pipeline RNPxl and OpenMS TOPPASViewer ^7^. Precursor mass tolerance was set to 6 ppm. MS/MS mass tolerance was set to 20 ppm. A neutral loss of 42.021798 Da (C1H2N2) at arginine residues was defined as well as adducts of ribose minus H_2_O (78.010565 Da; C5H2O), ADP-ribose (541.06111 Da; C15H21N5O13P2) and ADP-ribose minus adenine base (485.97295 Da; C10H17O16P3) ^8^. Results were filtered for 1 % false discovery rate (FDR) on peptide spectrum match (PSM) level. Ion chromatograms for rS1 peptides were extracted and visualized using Skyline (v. 21.2.0.369) ^9^.

### LC-MS/MS analysis of His-tagged rS1 isolated from T4 phage-infected *E. coli*

LC-MS/MS analysis of protein digests was performed on an Exploris 480 mass spectrometer connected to an electrospray ion source (Thermo Fisher Scientific). Peptide separation was carried out using Ultimate 3000 nanoLC-system (Thermo Fisher Scientific), equipped with packed in-house C18 resin column (Magic C18 AQ 2.4 μm, Dr. Maisch). The peptides were eluted from a pre-column in backflush mode with a gradient from 98 % solvent A (0.15 % formic acid) and 2 % solvent B (99.85 % acetonitrile, 0.15 % formic acid) to 35 % solvent B over 40 min and 90 min, respectively. The flow rate was set to 300 nl/min. The data dependent acquisition mode for label-free quantification was set to obtain one high-resolution MS scan at a resolution of 60000 (m/z 200) with scanning range from 350 to 1650 m/z. MS/MS scans were acquired either of the 20 most intense ions (90 min gradient) or for the most intense ions detected within 2s (cycle 1s, 40 min gradient). To increase the efficiency of MS/MS attempts, the charged state screening modus was adjusted to exclude unassigned and singly charged ions. The ion accumulation time was set to 25 ms for MS and “auto” for MS/MS scans. The automatic gain control (AGC) was set to 300 % for MS survey scans and 200 % for MS/MS scans.

MS raw spectra were analysed using MaxQuant (version 1.6.17.0 and 2.0.3.0) using a fasta database of the targets proteins and a set of common contaminant proteins. The following search parameters were used: full tryptic specificity required (cleavage after lysine or arginine residues); three missed cleavages allowed; carbamidomethylation (C) set as a fixed modification; and oxidation (M; +16 Da), deamidation (N, Q; +1 Da) and ADP-ribosylation (K; +541 Da) set as a variable modifications. MaxQuant was executed in defaults setting. All MaxQuant parameters are listed in Supplementary Tables 1 and 2. The MS proteomics data have been deposited with the ProteomeXchange Consortium via the PRIDE partner repository under the data set identifier PXD038075.

### Generation of *E. coli* B strain with endogenously His-tagged rS1

*E. coli* B strain with endogenously His-tagged rS1 was created by homologous recombination of linear transforming DNA (tDNA) using the pRET/ET plasmid in *E. coli* B strain. The linear tDNA was generated by fusion PCR aligning four fragments: 156 bp of the rpsA gene with an additional His-tag amplified from the pET28 rS1 vector (serving as the left homologous flank), a 70 bp fragment of the native rpsA terminator, the Flp-flanked kanamycin cassette from pKD4 and 140bp of the 3’-flanking region of the *rpsA* gene (right homologous flank). The utilised primers are indicated in Supplementary Table 8. The subsequent procedure for recombination is based on the protocol for the *E. coli* Gene Deletion Kit by RET/ET Recombination (Gene Bridges). Briefly, *E. coli* B strain containing the pRED/ET plasmid was grown in LB medium supplemented with 100 μg/ml ampicillin at 30 °C. At OD_600_ = 0.35, L-arabinose was added to 0.33 % (w/v) to induce expression of the RED/ET recombination system during growth at 37 °C for 1 h. 1.4 ml culture were harvested by centrifugation at 3,000 x g, 4 °C, 1 min, cells washed twice with 1 ml cold 10 % glycerol and finally resuspended in 50 μl 10 % glycerol. Cells were electroporated with 1 μg tDNA using a MicroPulser Electroporator (Bio-Rad) at standard settings (Ec1). Electroporated cells were immediately resuspended in 1 ml pre-warmed LB medium and incubated at 37 °C, 600 rpm for 3 h. Cells were finally plated on kanamycin (30 μg/ml) LB-agar plates. Cells took two days to recover and grow. Successful recombination was evaluated by Sanger sequencing and correct protein expression was validated by pull-down and proteomics.

### RNAylomeSeq

100 ml cultures of *E. coli* B strain with endogenously His-tagged rS1 (Supplementary Table 9) in LB medium supplied with 1 mM CaCl_2_, 1 mM MgCl_2_ and 30 μg/ml kanamycin were grown at 37 °C in 250 ml baffled Erlenmeyer flasks to an OD_600_ = ~ 0.8. T4 phage WT or T4 phage ModB R73A, G74A were added to an MOI of 5.0. For the uninfected negative control, same volumes of LB medium were added to the cultures. Cultures were then incubated at 37 °C for 8 min and *E. coli* harvested by centrifugation at 3,000 x g for 13 min. Dried pellets were stored at – 80 °C.

Pellets from 100 ml culture infected with either T4 phage WT, T4 phage ModB R73A G74A or the uninfected control (LB) were resuspended in 2 ml Ni-NTA wash buffer (10 mM imidazole, 50 mM Tris-HCl pH 7.5, 1 M NaCl, 1 M urea, 5 mM MgSO_4_, 5 mM β-mercaptoethanol, 5 % glycerol, pH 8.0, EDTA-free protease inhibitor (Roche, 1 tablet per 500 ml)) on ice and lysed by sonication (6 min, 50 % power, 0.5 s pulse). The lysate was cleared from cell debris by centrifugation at 21,000 x g, 4 °C, 30 min. 1.9 ml supernatant, 50 μl Ni-NTA agarose beads (Jena Bioscience, equilibrated in Ni-NTA wash buffer), 80 U murine RNase inhibitor (NEB) and 4.72 μg of rS1 D2 RNAylated with NAD-capped RNAI were combined and incubated at 4 °C in a Rotary Mixer for 30 min. Entire samples were transferred to Mobicol mini spin columns (MoBiTec). Beads were washed four times with 200 μl Ni-NTA wash buffer and subsequently eight times with 200 μl Strep wash buffer (50 mM Tris-HCl pH 7.5, 8 M urea). Beads were equilibrated in standard ligation buffer (10 mM MgCl_2_, 50 mM Tris-HCl pH 7.4) and blocked with BSA prior to 3’-RNA-adapter ligation at 4 °C o/n in the presence of standard ligation buffer, 50 mM β-mercaptoethanol, 0.05 μg/μl BSA, 15 % (v/v) DMSO, 5 μM adenylated RNA-3’-adapter, 0.5 U/μl T4 RNL1 (NEB) and 10 U/μl T4 RNL2, tr. K227Q (NEB). Protein was rebound by the addition of NaCl to 1.5 M and incubation at 20 °C, 400 rpm for 20 min. Beads were subsequently washed six times with strep wash buffer and equilibrated in first strand buffer (50 mM Tris-HCl pH 8.3, 3 mM MgCl_2_, 75 mM KCl) and blocked with BSA. Reverse transcription of protein-bound RNA was performed in a 30 μl scale for 1 h at 40 °C using 10 U/μl Superscript IV Reverse Transcriptase (Invitrogen) in the presence of 5 μM RT primer, first strand buffer, 25 mM β-mercaptoethanol, 0.05 μg/μl BSA and 0.5 mM dNTPs. After incubation, NaCl was added to 1.5 M and the solution was incubated at 20 °C, 400 rpm for 1 h in order to rebind RNA-cDNA hybrids. Beads were subsequently washed five times with 0.25 x strep wash buffer (2 M urea, 50 mM Tris-HCl pH 7.5), equilibrated in ExoI buffer (10 mM Tris-HCl pH 7.9, 5 mM β-mercaptoethanol, 10 mM MgCl_2_, 50 mM NaCl) and blocked with BSA. Residual RT primer was removed by ExoI digest with 1 U/μl *E. coli* ExoI (NEB) in ExoI buffer at 37 °C for at least 30 min. Beads were finally washed with 200 μl 0.25 x strep wash buffer five times and subsequently with 200 μl immobilisation buffer (10 mM Na-HEPES pH 7.2, 1 M NaCl) three times. cDNA was eluted by incubation of beads in 100 μl 150 mM NaOH at 55 °C for 25 min and by washing with 100 μl MQ water. Eluate pH was neutralised by addition of 0.05 volumes 3 M NaOAc pH 5.5. cDNA was removed from residual protein by phenol-chloroform extraction and precipitated with 2.5 volumes ethanol in the presence of 0.3 M NaOAc pH 5.5 o/n. Precipitated cDNA was C-tailed using 1 U/μl TdT (Thermo Fisher) in the presence of 1.25 mM CTP and 1 x TdT buffer at 37 °C for 30 min and subsequently inactivated at 70 °C for 10 min. 5 μM cDNA anchor (hybridisation of fwd and rev anchor) were ligated to C-tailed cDNA in standard ligation buffer in the presence of 10 μM ATP and 1.5 U/μl T4 DNA Ligase (Thermo Fisher Scientific) at 4 °C o/n. Ligation reactions were inactivated at 70 °C for 10 min and cDNA was ethanol-precipitated.

For the preparation of the Illumina RNAylomeSeq library, cDNA was amplified by PCR using 2 U Phusion Polymerase (Thermo Fisher Scientific) in the presence of 5 % (v/v) DMSO, 200 μM dNTPs and 2500 nM NEBNext Universal and Index Primer each (Primer Set 1, NEB). PCR products were purified by native PAGE and ethanol-precipitated. dsDNA concentration was determined via Quantus fluorometer (Promega) and library size was determined with the Bioanalyzer (Agilent). Equimolar amounts of each library were sequenced on a MiniSeq system (Illumina) using the MiniSeq High-Output Kit (150 cycles, Illumina) generating 20 million 151 bp single-end reads.

### Analysis of NGS Data

NGS data were demultiplexed using bcl2fastq (version 2.20.0, Illumina). Fastq files were assessed using FastQC (version 0.11.9) and Illumina sequencing adapters were trimmed from reads using cutadapt (version 1.18). Reads were aligned to a reference genome composed of an *E. coli* K12 (U00096.3), bacteriophage T4 (NC_000866.4) and RNAI (self-designed) with hisat2 (version 2.2.1). Primary alignments were selected using samtools (version 1.7) and reads per genomic feature were counted with featureCounts (version 2.0.1 from Subread package). The resulting counts table was subjected to further analysis and data visualisation in R (version 4.1.2). Read counts were normalised to the overall number of mapped reads per sample and to the respective read counts for the RNAI spike-in as follows:

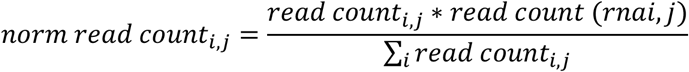

Data visualisation was performed with a custom R script and alignments were manually inspected in Integrative Genomics Viewer (IGV, version 2.4.9). Hits were identified based on the following criteria: log2 fold change (LFC) ≥ 1.5 comparing T4 WT and T4 R73A, G74A mutant and a log2-normalised mean expression among WT and R73A, G74A sample of one replicate ≥ −0.5.

### qPCR validation of NGS data

cDNAs from RNAylomeSeq were diluted 1:30 in Millipore water. qPCRs were performed on 1 μl diluted cDNAs in 10 μl scale in technical duplicates amplifying regions of 100 – 150 bp with the iTaq Universal SYBR Green Supermix (Bio-Rad) according to manufacturer’s instructions using primers indicated in Table 8. The log2 of the difference in ct-values for T4 WT and T4 R73A, G74A infected samples from corresponding replicates was computed and an LFC ≥ 1 was set as a threshold for cDNA enrichment.

### Ribosome RNAylation and proteomic analysis of RNAylated proteins

4.3 μg/μl 70S ribosomes were RNAylated in transferase buffer in the presence of either 1 μM NAD-10mer-Cy5 or 1 μM NAD-40mer-Cy5 (Supplementary Table 6) by 0.05 μg/μl ModB at 15 °C for 90 min. RNAylated and non-RNAylated control samples were subjected to 12 % SDS-PAGE analysis. To identify RNAylated proteins, SDS-PAGE-separated protein bands were excised and proteins were in-gel digested as described previously ^10^. LC-MS was carried out on an Exploris 480 instrument connected to an Ultimate 3000 RSLC nano with a Prowflow upgrade and a nanospray flex ion source (all Thermo Scientific). Peptide mixtures were then analysed on the LC-MS system described above with a peptide separating gradient of 30 min from 2 % to 35 % buffer B. Peptide separation was performed on a reverse phase HPLC column (75 μm x 42 cm) packed in-house with C18 resin (2.4 μm, Dr. Maisch). Peptides were ionised at 2.3 kV spray voltage with a heated capillary temperature at 275 °C and funnel RF level at 40. MS survey scans were acquired with a resolution of 120.000 at m/z 200 and full MS AGC target of 300 % with a maximal IT of 50 ms. The mass range was set to 350–1650. The fragment spectra were acquired in data-dependent acquisition mode with a quadrupole isolation window of m/z 1.5, an AGC target value of 200 % and a resolution of 15.000, and fragmentation was induced with a normalised HCD collision energy of 27 %. MS raw data was searched with SEQUEST embedded in Proteome Discoverer 2.2 (Thermo Scientific) against an Uniprot *E. coli* protein database containing the bacteriophage T4 protein ModB.

### RNAylation of proteins in *E. coli* lysates

A fresh pellet from 40 ml *E. coli* B strain culture at OD_600_ = ~ 0.8 was resuspended in 2 ml transferase buffer (10 mM Mg(OAc)_2_, 22 mM NH_4_Cl, 50 mM Tris-acetate pH 7.5, 1 mM EDTA, 10 mM 2-mercaptoethanol, 1 % glycerol). Cells were lysed by sonication (3 x 2 min at 50 % power, 0.5 s pulse) and the lysate was cleared from cell debris by centrifugation at 27,670 x g, 4 °C for 30 min. The supernatant was used in RNAylation assays.

100 μl lysate was incubated in the presence of 0.93 μM NAD-10mer-Cy5 or 0.93 μM P-10mer-Cy5 (Supplementary Table 6), 0.37 U murine RNase inhibitor (NEB) and 0.69 μM ModB at 15 °C. 10 μl samples were taken before the addition of ModB (t0), 2, 5, 10, 20, 30 and 60 minutes and immediately resuspended in 1 volume 2 x Laemmli buffer. Samples were analysed by 12 % SDS-PAGE applying the same reference (rS1 RNAylated with NAD-10mer-Cy5) to each gel. The Cy5 signal was recorded using the Cy5 blot option of the ChemiDoc Imaging System at 90 s manual exposure. Afterwards, gels were stained in Coomassie solution and imaged with the same system.

### Western Blotting

Proteins were separated by 10 % SDS-PAGE and gels were equilibrated in transfer buffer (25 mM Tris, pH 8.3, 192 mM glycine, 20 % [v/v] methanol). 0.2 μm polyvinylidene difluoride (PVDF) membranes (GE Healthcare) were activated in methanol for 1 min and equilibrated in transfer buffer. Proteins were transferred from gels to PVDF membranes in a semi-dry manner at 300 mA for 1.5 h, if not indicated differently. After the transfer, membranes were dehydrated by soaking in methanol and washed 2 x with TBS-Tween (TBS-T; 10 mM Tris-HCl, pH 7.5, 150 mM NaCl, 0.05 % [v/v] Tween^®^ 20). Afterwards, 10 ml blocking buffer (5 % [w/v] milk powder in TBS-T) were added to the membranes and incubated at room temperature for 1 h. To detect ADP-ribosylated proteins, membranes were incubated with a 1:10,000 dilution of primary antibody MABE1016 (Merck) in 10 ml washing buffer (1 % [w/v] milk powder in TBS-T) at 4 °C o/n ^11^. Membranes were washed and incubated with 10 ml of a 1:10,000 dilution of the horseradish-peroxidase-(HRP)-goat-anti-rabbit-IgG secondary antibody (Advansta) in washing buffer at room temperature for 1 h. Afterwards, membranes were washed with PBS. ADP-ribosylated proteins were visualised by chemiluminescence using the SignalFire ECL Reagent or the SignalFire Elite ECL Reagent (Cell Signaling Technology) according to the manufacturer’s instructions. If proteins in SDS-PAGE gels needed to be visualised before blotting, a 2,2,2-trichloroethanol (TCE) staining method ^12^ was used. Resolving gels were supplemented with 0.5 % (v/v) TCE. For visualisation, gels were activated by ultraviolet transillumination (300 nm) for 60 s. Proteins then showed fluorescence in the visible spectrum.

### Quantification of RNAylation

rS1 proteins were isolated from *E. coli* strain B pTAC rS1 (Supplementary Table 9) infected or non-infected with bacteriophage T4. 1.5 μM rS1 were digested with 1 μM ARH1 in the presence of 12.5 mM Tris-HCl pH 7.5, 25 mM NaCl, 25 mM KCl and 5 mM MgCl_2_. 1.5 μM rS1 were digested with 0.5 U endonuclease P1 in 100 mM Mg(OAc)_2_, 220 mM NH_4_Cl, 500 mM HEPES pH 7.5, 10 mM EDTA, 100 mM β-mercaptoethanol and 10 % glycerol. Digests were incubated at 37 °C for 2 h. Afterwards, digests were precipitated by the addition of 9 volumes of ethanol and precipitated by centrifugation (14,000 rpm) at 4 °C for 1 h. Protein pellets were resuspended in 10 μl 1 x Laemmli buffer and analysed via Western blot. ADPr-modifications were detected by the primary antibody MABE1016 (Merck) as described above.

### T4 phage mutagenesis

The CRISPR-Cas9 spacer plasmids were generated by introducing the *modB* spacer sequence into the DS-SPCas plasmid (Addgene, no.48645) (Supplementary Table 9). The *modB* carrying vector pET28_ModB was used as a donor DNA for homologous recombination in CRISPR-Cas9 mediated mutagenesis. The pET28_ModB plasmid was modified by site-directed mutagenesis, during which point mutations R73A and G74A were introduced to *modB*. The R73A mutation led to the inactivation of ADP-ribosyltransferase activity. The G74A mutation was located in the protospacer adjacent motif (PAM) and was required to prevent the cleavage of donor DNA by Cas9 nuclease. The applied primers are listed in Supplementary Table 8. The resulting plasmids were sequenced and transformed into *E*. *coli* BL21 (DE3).

The CRISPR-Cas9-mediated mutagenesis was based on the publication of Tao et al^13^. The DS_SPCas_ModB plasmid with the target spacer sequence and the donor plasmid pET28a_ModB_R73A/G74A were co-transformed into *E. coli* DH5α. The cells were further infected by bacteriophage T4 (1:10000 phages/cells), and the plaque assay was performed. The plates were incubated o/n at 37 °C, and the resulting plaques were screened for mutants. Single plaques were picked by sterile pipet tips and transferred into 200 μl Pi-Mg buffer (26 mM Na_2_HPO_4_, 68 mM NaCl, 22 mM KH_2_PO_4_, 1 mM MgSO_4_, pH7.5) supplied with 2 μl CCl_3_H. The samples were incubated at room temperature for 1 h. Next, 2 μl were transferred to a new PCR tube and heated up to 95 °C for 10 min. The sample was further used for DNA amplification via PCR (used primers are listed in Supplementary Table 8). The amplified DNA was purified via agarose gel electrophoresis and submitted for Sanger sequencing.

### Plaque assay

*E. coli* culture of interest was grown to OD_600_ = ~ 0.8-1.0. Next, 300 μl of the culture were infected with 100 μl of T4 WT phage or T4 ModB R73A, G74A (Supplementary Table 9) mutant with either defined or unknown MOI. The bacteria-phage suspension was incubated at 37 °C for 7 min and subsequently transferred to 4 ml of LB-soft-agar (0.75 %), properly mixed, and poured onto an LB-agar plate (1.5 % LB-agar). The plates were incubated at 37 °C o/n and validated on the following day.

### Time course of T4-mediated lysis of *E. coli*

100 ml of LB medium (in 500 ml baffled flasks) were inoculated with *E. coli* B o/n culture to the start OD_600_ = 0.1 and incubated at 37 °C and 180 rpm until OD_600_ = 0.8 was reached. The culture was cooled down to room temperature and infected either by T4 WT phage or T4 ModB R73A, G74A mutant (Supplementary Table 9) to an MOI of 5. The culture was further incubated at room temperature and 150 rpm. The cell lysis was tracked by measurement of OD_600_ at different time points of infection (0 – 200 min post-infection). The experiment was run in biological triplicates.

### Burst size determination

100 ml of LB medium (500 ml baffled flasks) were inoculated with *E. coli* B o/n culture to the start OD_600_ = 0.1 and incubated at 37 °C and 180 rpm till OD_600_ = 0.8 was reached. Afterwards, the culture was infected either by T4 WT phage or T4 ModB R73A, G74A mutant (Supplementary Table 9) to an MOI of 0.01 and further incubated at 37 °C without shaking.

To determine the number of total infective centers T_0_ (unadsorbed and already adsorbed phages), at 5 min post-infection, 100 μl of infected culture were used for re-infection of 300 μl of *E. coli* B cells (OD_600_ = 1.0) and subsequential plaque assay. The number of unabsorbed phages (U) was determined by transferring 1 ml of infected culture to 50 μl CCl_3_H. In this way, the disruption of *E. coli* cells took place, whereupon the unadsorbed phages remained intact and were used for plaque assay. The T_0_-U difference represented the number of initially infected centers. The number of unadsorbed phages (U_xmin_) was continuously traced during infection and used for the calculation of T4 phage progeny (T4 phage progeny=U_xmin_/(T_0_-U_5min_)). The time point at which the first increase in phage number was observed was treated as the first burst time point and used to calculate the phage burst size (burst size=U_1st burst_/(T_0_-U_5min_)).

### Bacteriophage adsorption kinetics

100 ml of LB medium (in 500 ml baffled flasks) were inoculated with *E. coli* B o/n culture to an OD_600_ = 0.1 and incubated at 37 °C and 180 rpm until OD_600_ = 0.8 was reached. The culture was cooled down to room temperature and infected either by T4 WT phage or T4 ModB R73A, G74A mutant (Supplementary Table 9) to an MOI of 0.1. Directly after infection, 100 μl of the culture were used to determine the number of total infective centers T_0_ via plaque assay. Further, 100 μl of the culture were taken at different time points of infection (0-25 min post-infection) and were supplied with 5 μl of CCl_3_H to disrupt *E. coli* cells. This suspension was subsequently used to determine the number of unadsorbed phages (U_xmin_) via plaque assay. The calculation of the adsorption rate was performed as follows:

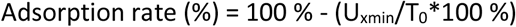

## REFERENCES

1 Tiemann, B. et al. ModA and ModB, two ADP-ribosyltransferases encoded by bacteriophage T4: catalytic properties and mutation analysis. J Bacteriol 186, 7262–7272 2004). https://doi.org:10.1128/JB.186.21.7262-7272.2004

2 Koch, T., Raudonikiene, A., Wilkens, K. & Rüger, W. Overexpression, purification, and characterization of the ADP-ribosyltransferase (gpAlt) of bacteriophage T4: ADP-ribosylation of E. coli RNA polymerase modulates T4 “early” transcription. Gene Expr 4, 253–264 (1995).

3 Cahova, H., Winz, M. L., Höfer, K., Nübel, G. & Jäschke, A. NAD captureSeq indicates NAD as a bacterial cap for a subset of regulatory RNAs. Nature 519, 374–377 (2015). https://doi.org:10.1038/nature14020

4 Chen, Y. G., Kowtoniuk, W. E., Agarwal, I., Shen, Y. & Liu, D. R. LC/MS analysis of cellular RNA reveals NAD-linked RNA. Nat Chem Biol 5, 879–881 (2009). https://doi.org:10.1038/nchembio.235

5 Jiao, X. et al. 5’ End Nicotinamide Adenine Dinucleotide Cap in Human Cells Promotes RNA Decay through DXO-Mediated deNADding. Cell 168, 1015–1027 e1010 (2017). https://doi.org:10.1016/j.cell.2017.02.019

6 Fehr, A. R. et al. The impact of PARPs and ADP-ribosylation on inflammation and host-pathogen interactions. Genes Dev 34, 341–359 (2020). https://doi.org:10.1101/gad.334425.119

7 Cohen, M. S. & Chang, P. Insights into the biogenesis, function, and regulation of ADP-ribosylation. Nat Chem Biol 14, 236–243 (2018). https://doi.org:10.1038/nchembio.2568

8 Simon, N. C., Aktories, K. & Barbieri, J. T. Novel bacterial ADP-ribosylating toxins: structure and function. Nat Rev Microbiol 12, 599–611 (2014). https://doi.org:10.1038/nrmicro3310

9 Luscher, B. et al. ADP-Ribosylation, a Multifaceted Posttranslational Modification Involved in the Control of Cell Physiology in Health and Disease. Chem Rev 118, 1092–1136 (2018). https://doi.org:10.1021/acs.chemrev.7b00122

10 Morales-Filloy, H. G. et al. The 5’ NAD Cap of RNAIII Modulates Toxin Production in Staphylococcus aureus Isolates. J Bacteriol 202 (2020). https://doi.org:10.1128/JB.00591-19

11 Frindert, J. et al. Identification, Biosynthesis, and Decapping of NAD-Capped RNAs in B. subtilis. Cell Rep 24, 1890–1901 e1898 (2018). https://doi.org:10.1016/j.celrep.2018.07.047

12 Ruiz-Larrabeiti, O. et al. NAD+capping of RNA in Archaea and Mycobacteria. bioRxiv, 2021.2012.2014.472595 (2021). https://doi.org:10.1101/2021.12.14.472595

13 Gomes-Filho, J. V. et al. Identification of NAD-RNAs and ADPR-RNA decapping in the archaeal model organisms Sulfolobus acidocaldarius and Haloferax volcanii. bioRxiv, 2022.2011.2002.514978 (2022). https://doi.org:10.1101/2022.11.02.514978

14 Walters, R. W. et al. Identification of NAD+ capped mRNAs in Saccharomyces cerevisiae. Proc Natl Acad Sci U S A 114, 480–485 (2017). https://doi.org:10.1073/pnas.1619369114

15 Zhang, Y. et al. Extensive 5’-surveillance guards against non-canonical NAD-caps of nuclear mRNAs in yeast. Nat Commun 11, 5508 (2020). https://doi.org:10.1038/s41467-020-19326-3

16 Wang, J. et al. Quantifying the RNA cap epitranscriptome reveals novel caps in cellular and viral RNA. Nucleic Acids Res 47, e130 (2019). https://doi.org:10.1093/nar/gkz751

17 Zhang, H. et al. NAD tagSeq reveals that NAD(+)-capped RNAs are mostly produced from a large number of protein-coding genes in Arabidopsis. Proc Natl Acad Sci U S A 116, 12072–12077 (2019). https://doi.org:10.1073/pnas.1903683116

18 Wang, Y. et al. NAD(+)-capped RNAs are widespread in the Arabidopsis transcriptome and can probably be translated. Proc Natl Acad Sci U S A 116, 12094–12102 (2019). https://doi.org:10.1073/pnas.1903682116

19 Dong, H. et al. NAD(+)-capped RNAs are widespread in rice (Oryza sativa) and spatiotemporally modulated during development. Sci China Life Sci 65, 2121–2124 (2022). https://doi.org:10.1007/s11427-021-2113-7

20 Wolfram-Schauerte, M. & Höfer, K. NAD-capped RNAs - a redox cofactor meets RNA. Trends Biochem Sci (2022). https://doi.org:10.1016/j.tibs.2022.08.004

21 Höfer, K. & Jäschke, A. Epitranscriptomics: RNA Modifications in Bacteria and Archaea. Microbiol Spectr 6 (2018). https://doi.org:10.1128/microbiolspec.RWR-0015-2017

22 Miller, E. S. et al. Bacteriophage T4 genome. Microbiol Mol Biol Rev 67, 86–156, table of contents (2003). https://doi.org:10.1128/MMBR.67.1.86-156.2003

23 Rohrer, H., Zillig, W. & Mailhammer, R. ADP-ribosylation of DNA-dependent RNA polymerase of Escherichia coli by an NAD+: protein ADP-ribosyltransferase from bacteriophage T4. Eur J Biochem 60, 227–238 (1975). https://doi.org:10.1111/j.1432-1033.1975.tb20995.x

24 Depping, R., Lohaus, C., Meyer, H. E. & Rüger, W. The mono-ADP-ribosyltransferases Alt and ModB of bacteriophage T4: target proteins identified. Biochem Biophys Res Commun 335, 1217–1223 (2005). https://doi.org:10.1016/j.bbrc.2005.08.023

25 Skorko, R., Zillig, W., Rohrer, H., Fujiki, H. & Mailhammer, R. Purification and properties of the NAD+: protein ADP-ribosyltransferase responsible for the T4-phage-induced modification of the alpha subunit of DNA-dependent RNA polymerase of Escherichia coli. Eur J Biochem 79, 55–66 (1977). https://doi.org:10.1111/j.1432-1033.1977.tb11783.x

26 Tiemann, B., Depping, R. & Rüger, W. Overexpression, purification, and partial characterization of ADP-ribosyltransferases modA and modB of bacteriophage T4. Gene Expr 8, 187–196 (1999).

27 Rankin, P. W., Jacobson, E. L., Benjamin, R. C., Moss, J. & Jacobson, M. K. Quantitative studies of inhibitors of ADP-ribosylation in vitro and in vivo. J Biol Chem 264, 4312–4317 (1989).

28 Goelz, S. & Steitz, J. A. Escherichia coli ribosomal protein S1 recognizes two sites in bacteriophage Qbeta RNA. J Biol Chem 252, 5177–5179 (1977).

29 Cervantes-Laurean, D., Jacobson, E. L. & Jacobson, M. K. Glycation and glycoxidation of histones by ADP-ribose. J Biol Chem 271, 10461–10469 (1996). https://doi.org:10.1074/jbc.271.18.10461

30 Kramer, K. et al. Photo-cross-linking and high-resolution mass spectrometry for assignment of RNA-binding sites in RNA-binding proteins. Nat Methods 11, 1064–1070 (2014). https://doi.org:10.1038/nmeth.3092

31 Gehrig, P. M. et al. Gas-Phase Fragmentation of ADP-Ribosylated Peptides: Arginine-Specific Side-Chain Losses and Their Implication in Database Searches. J Am Soc Mass Spectrom 32, 157–168 (2021). https://doi.org:10.1021/jasms.0c00040

32 Winz, M. L. et al. Capture and sequencing of NAD-capped RNA sequences with NAD captureSeq. Nat Protoc 12, 122–149 (2017). https://doi.org:10.1038/nprot.2016.163

33 Tao, P., Wu, X., Tang, W. C., Zhu, J. & Rao, V. Engineering of Bacteriophage T4 Genome Using CRISPR-Cas9. Acs Synth Biol 6, 1952–1961 (2017). https://doi.org:10.1021/acssynbio.7b00179

34 Zhang, H. et al. Use of NAD tagSeq II to identify growth phase-dependent alterations in E. coli RNA NAD(+) capping. Proc Natl Acad Sci U S A 118 (2021). https://doi.org:10.1073/pnas.2026183118

35 Bird, J. G. et al. The mechanism of RNA 5’ capping with NAD+, NADH and desphospho-CoA. Nature 535, 444–447 (2016). https://doi.org:10.1038/nature186228

36 Bycroft, M., Hubbard, T. J., Proctor, M., Freund, S. M. & Murzin, A. G. The solution structure of the S1 RNA binding domain: a member of an ancient nucleic acid-binding fold. Cell 88, 235–242 (1997). https://doi.org:10.1016/s0092-8674(00)81844-9

37 Bennett, B. D. et al. Absolute metabolite concentrations and implied enzyme active site occupancy in Escherichia coli. Nat Chem Biol 5, 593–599 (2009). https://doi.org:10.1038/nchembio.186

38 Diedrich, G. et al. Ribosomal protein L2 is involved in the association of the ribosomal subunits, tRNA binding to A and P sites and peptidyl transfer. EMBO J 19, 5241–5250 (2000). https://doi.org:10.1093/emboj/19.19.5241

39 Nakagawa, A. et al. The three-dimensional structure of the RNA-binding domain of ribosomal protein L2; a protein at the peptidyl transferase center of the ribosome. EMBO J 18, 1459–1467 (1999). https://doi.org:10.1093/emboj/18.6.1459

40 Hentze, M. W., Castello, A., Schwarzl, T. & Preiss, T. A brave new world of RNA-binding proteins. Nat Rev Mol Cell Biol 19, 327–341 (2018). https://doi.org:10.1038/nrm.2017.130

41 Drygin, Y. F. Natural covalent complexes of nucleic acids and proteins: some comments on practice and theoryon the path from well-known complexes to new ones. Nucleic Acids Res 26, 4791–4796 (1998). https://doi.org:10.1093/nar/26.21.4791

42 Ibba, M. & Soll, D. Aminoacyl-tRNA synthesis. Annu Rev Biochem 69, 617–650 (2000). https://doi.org:10.1146/annurev.biochem.69.1.617

43 Rekosh, D. M., Russell, W. C., Bellet, A. J. & Robinson, A. J. Identification of a protein linked to the ends of adenovirus DNA. Cell 11, 283–295 (1977). https://doi.org:10.1016/0092-8674(77)90045-9

44 Rothberg, P. G., Harris, T. J., Nomoto, A. & Wimmer, E. O4-(5’-uridylyl)tyrosine is the bond between the genome-linked protein and the RNA of poliovirus. Proc Natl Acad Sci U S A 75, 4868–4872 (1978). https://doi.org:10.1073/pnas.75.10.4868

45 Uzan, M. & Miller, E. S. Post-transcriptional control by bacteriophage T4: mRNA decay and inhibition of translation initiation. Virol J 7, 360 (2010). https://doi.org:10.1186/1743-422X-7-360

46 Ke, Y., Zhang, J., Lv, X., Zeng, X. & Ba, X. Novel insights into PARPs in gene expression: regulation of RNA metabolism. Cell Mol Life Sci 76, 3283–3299 (2019). https://doi.org:10.1007/s00018-019-03120-6

47 Otten, E. G. et al. Ubiquitylation of lipopolysaccharide by RNF213 during bacterial infection. Nature 594, 111–116 (2021). https://doi.org:10.1038/s41586-021-03566-4

48 Flynn, R. A. et al. Small RNAs are modified with N-glycans and displayed on the surface of living cells. Cell 184, 3109–3124 e3122 (2021). https://doi.org:10.1016/j.cell.2021.04.023

49 Tsuge, H. et al. Structural basis of actin recognition and arginine ADP-ribosylation by Clostridium perfringens iota-toxin. Proc Natl Acad Sci U S A 105, 7399–7404 (2008). https://doi.org:10.1073/pnas.0801215105

50 Beckert, B. et al. Structure of a hibernating 100S ribosome reveals an inactive conformation of the ribosomal protein S1. Nat Microbiol 3, 1115–1121 (2018). https://doi.org:10.1038/s41564-018-0237-0

## REFERENCES (Methods only)

1 Höfer, K., Abele, F., Schlotthauer, J. & Jäschke, A. Synthesis of 5’-NAD-Capped RNA. Bioconjug Chem 27, 874–877 (2016). https://doi.org:10.1021/acs.bioconjchem.6b00072

2 Hsia, J. A. et al. Amino acid-specific ADP-ribosylation. Sensitivity to hydroxylamine of [cysteine(ADP-ribose)]protein and [arginine(ADP-ribose)]protein linkages. J Biol Chem 260, 16187–16191 (1985).

3 McDonald, L. J., Wainschel, L. A., Oppenheimer, N. J. & Moss, J. Amino acid-specific ADP-ribosylation: structural characterization and chemical differentiation of ADP-ribose-cysteine adducts formed nonenzymatically and in a pertussis toxin-catalyzed reaction. Biochemistry 31, 11881–11887 (1992). https://doi.org:10.1021/bi00162a029

4 Silberklang, M., Gillum, A. M. & RajBhandary, U. L. The use of nuclease P1 in sequence analysis of end group labeled RNA. Nucleic Acids Res 4, 4091–4108 (1977). https://doi.org:10.1093/nar/4.12.4091

5 Abplanalp, J. et al. Proteomic analyses identify ARH3 as a serine mono-ADP-ribosylhydrolase. Nat Commun 8, 2055 (2017). https://doi.org:10.1038/s41467-017-02253-1

6 Banasik, M., Komura, H., Shimoyama, M. & Ueda, K. Specific inhibitors of poly(ADP-ribose) synthetase and mono(ADP-ribosyl)transferase. J Biol Chem 267, 1569–1575 (1992).

7 Kramer, K. et al. Photo-cross-linking and high-resolution mass spectrometry for assignment of RNA-binding sites in RNA-binding proteins. Nat Methods 11, 1064–1070 (2014). https://doi.org:10.1038/nmeth.3092

8 Gehrig, P. M. et al. Gas-Phase Fragmentation of ADP-Ribosylated Peptides: Arginine-Specific Side-Chain Losses and Their Implication in Database Searches. J Am Soc Mass Spectrom 32, 157–168 (2021). https://doi.org:10.1021/jasms.0c00040

9 MacLean, B. et al. Skyline: an open source document editor for creating and analyzing targeted proteomics experiments. Bioinformatics 26, 966–968(2010). https://doi.org:10.1093/bioinformatics/btq054

10 Dell’Aquila, G. et al. Mobilization and Cellular Distribution of Phosphate in the Diatom Phaeodactylum tricornutum. Front Plant Sci 11, 579 (2020). https://doi.org:10.3389/fpls.2020.00579

11 Gibson, B. A., Conrad, L. B., Huang, D. & Kraus, W. L. Generation and Characterization of Recombinant Antibody-like ADP-Ribose Binding Proteins. Biochemistry 56, 6305–6316 (2017). https://doi.org:10.1021/acs.biochem.7b00670

12 Ladner, C. L., Yang, J., Turner, R. J. & Edwards, R. A. Visible fluorescent detection of proteins in polyacrylamide gels without staining. Anal Biochem 326, 13–20 (2004). https://doi.org:10.1016/j.ab.2003.10.047

13 Tao, P., Wu, X., Tang, W. C., Zhu, J. & Rao, V. Engineering of Bacteriophage T4 Genome Using CRISPR-Cas9. ACS Synth Biol 6, 1952–1961 (2017). https://doi.org:10.1021/acssynbio.7b00179

