## supplemental Files for "A viral ADP-ribosyltransferase attaches RNA chains to host proteins"

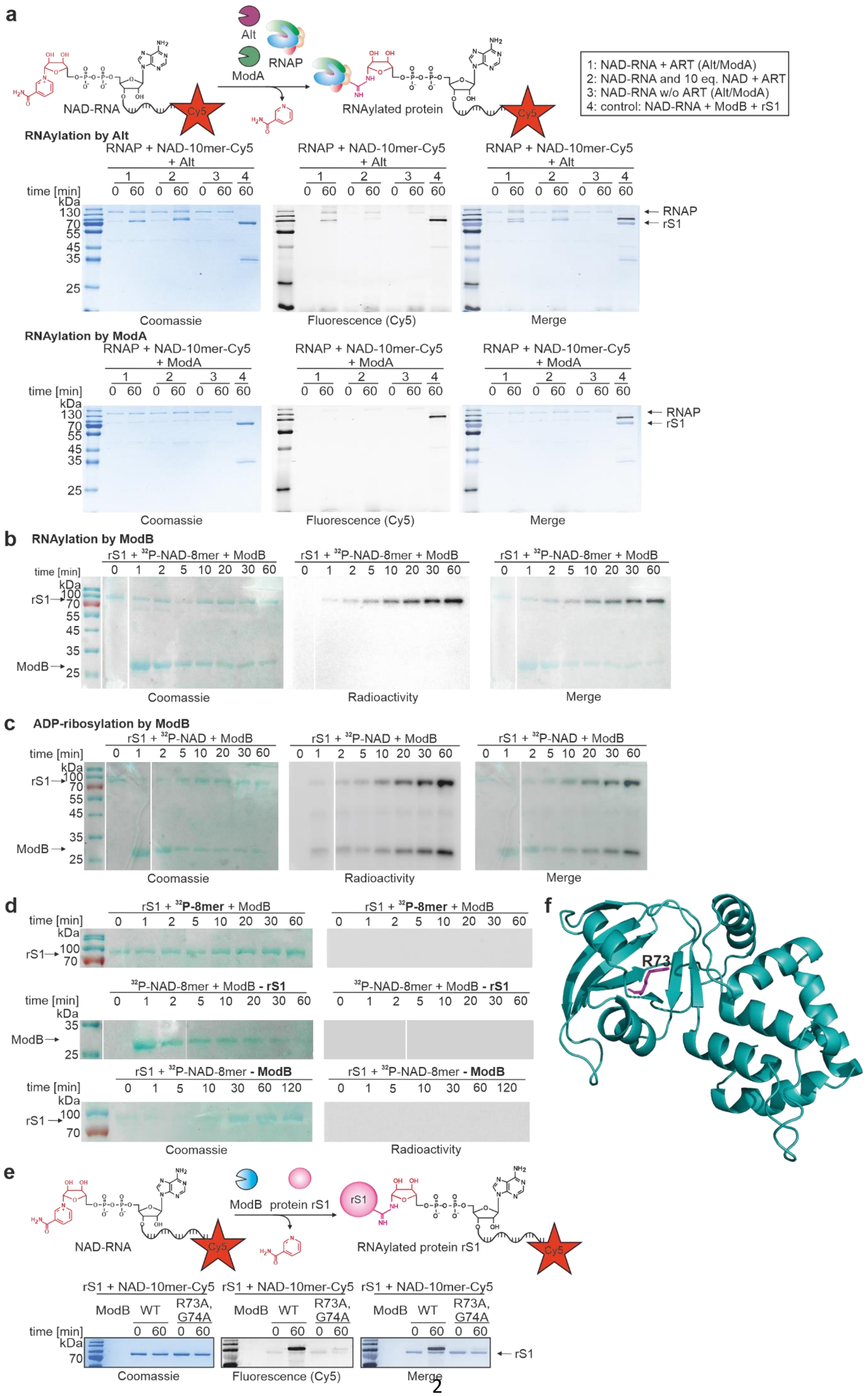
a
Alt
RNAP
ModA
NAD-RNA
Cy5
RNAylated protein
1: NAD-RNA + ART (Alt/ModA)
2: NAD-RNA and 10 eq. NAD + ART
3: NAD-RNA w/o ART (Alt/ModA)
4: control: NAD-RNA + ModB + rS1
RNAylation by Alt
RNAP + NAD-10mer-Cy5 + Alt
time [min]
kDa
RNAP
rS1
Coomassie
Fluorescence (Cy5)
Merge
RNAylation by ModA
RNAP + NAD-10mer-Cy5 + ModA
b
RNAylation by ModB
rS1 + 32P-NAD-8mer + ModB
ModB
Radioactivity
c
ADP-ribosylation by ModB
rS1 + 32P-NAD + ModB
d
rS1 + 32P-8mer + ModB
32P-NAD-8mer + ModB - rS1
rS1 + 32P-NAD-8mer - ModB
f
R73
e
ModB
protein rS1
RNAylated protein rS1
rS1 + NAD-10mer-Cy5
WT
R73A, G74A

**Extended Data Fig. 1: ADP-ribosylation and RNAylation by T4 ARTs. a**, Characterisation of RNAylation of the RNA polymerase (RNAP) by the ARTs Alt or ModA in the presence of NAD-10mer-Cy5 (1), additional 10 equivalents of NAD (2) or in the absence of the respective ART (3) (n=3). rS1 RNAylated with NAD-10mer-Cy5 by ModB serves as a reference (4). The RNAP is a well-established target protein of Alt and ModA and was thus chosen to assess RNAylation by Alt and ModA. Alt slightly RNAylates the RNAP *in vitro* which is abolished in the presence of 10 equivalents of NAD relative to NAD-10mer-Cy5. Protein load is visualised by Coomassie staining and RNAylated protein is visualised by the fluorescent Cy5 channel. **b**, Time course of the ADP-ribosylation of rS1 by ModB analysed by SDS-PAGE (n=3). ADP-ribosylated protein is visualised by radioactivity scan and protein load confirmed by Coomassie staining. **c**, Time course analysis of the ModB-mediated RNAylation of rS1 analysed by SDS-PAGE (n=3). RNAylated protein is visualised by radioactivity scan and protein load confirmed by Coomassie staining. **d**, Negative controls for RNAylation of rS1 with ModB analysed by SDS-PAGE. RNAylation assay was performed in the presence of $^{32}$P-RNA and in the absence of either rS1 (- rS1) or ModB (-ModB) (n=3). In these experiments, no RNAylation was detected in the radioactive scan of the SDS-PAGE gel. **e**, RNAylation of rS1 in the presence of catalytically active ModB and catalytically inactive ModB R73A, G74A with NAD-10mer-Cy5 (n=3). In addition to the catalytically important residue R73, we mutated G74 as well. Mutation of G74 results in an altered PAM region, which is important for CRISPR-Cas9 gene editing of the T4 phage genome. **f**, AlphaFold prediction [1] of the structure of ModB. Residue R73 is highlighted in magenta.

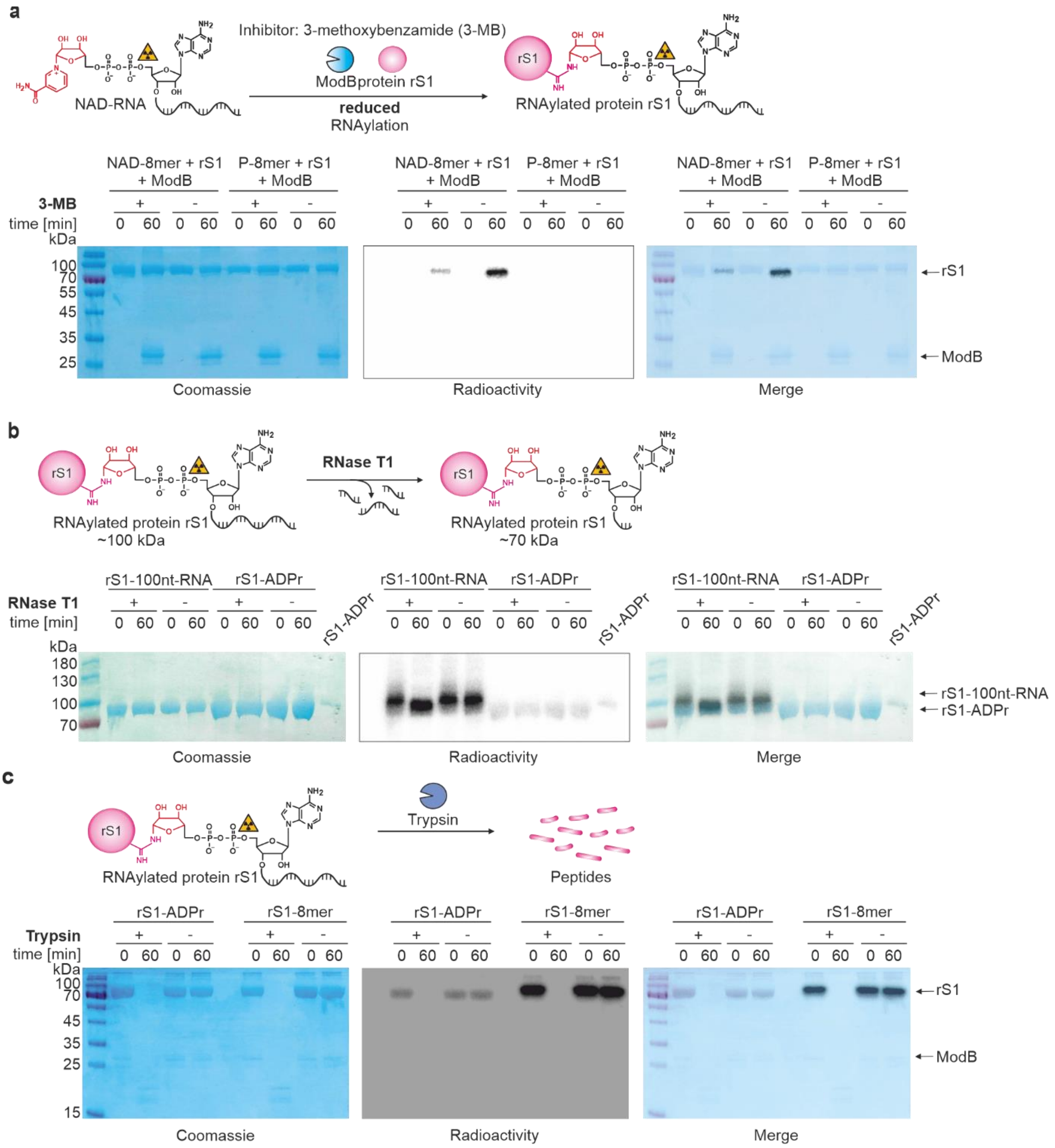


**Extended Data Fig. 2: Characterisation of the RNAylation of protein rS1 by ModB. a**, Inhibition of *in vitro* RNAylation of protein rS1 by ModB via ART inhibitor 3-MB. Reactions were performed with $^{32}$P-NAD-RNA 8mer ($^{32}$P-NAD-8mer) as well as $^{32}$P-RNA 8mer (negative control) (n=3). 3-MB reduces the yield of RNAylated rS1. **b**, *in vitro* digest of RNAylated and ADP-ribosylated protein rS1 by RNase T1. Reactions performed in the absence of RNase T1 (-) serve as negative controls. Protein rS1 ADP-ribosylated in the presence of $^{32}$P-NAD was applied as a reference (S1-ADPr) (n=2). Upon T1 digest, the 100nt-RNA at rS1 is shortened, and the molecular weight of RNAylated rS1 is reduced. This leads to similar electrophoretic mobility as for ADP-ribosylated rS1. **c**, Tryptic digest of ADP-ribosylated and RNAylated protein rS1 (n=2). The protein is degraded in the presence of trypsin, and RNAylation and ADP-ribosylation signals are lost. All samples were analysed by 12 % SDS-PAGE, protein was visualised by Coomassie staining and RNAylation was assessed via a radioactivity scan.

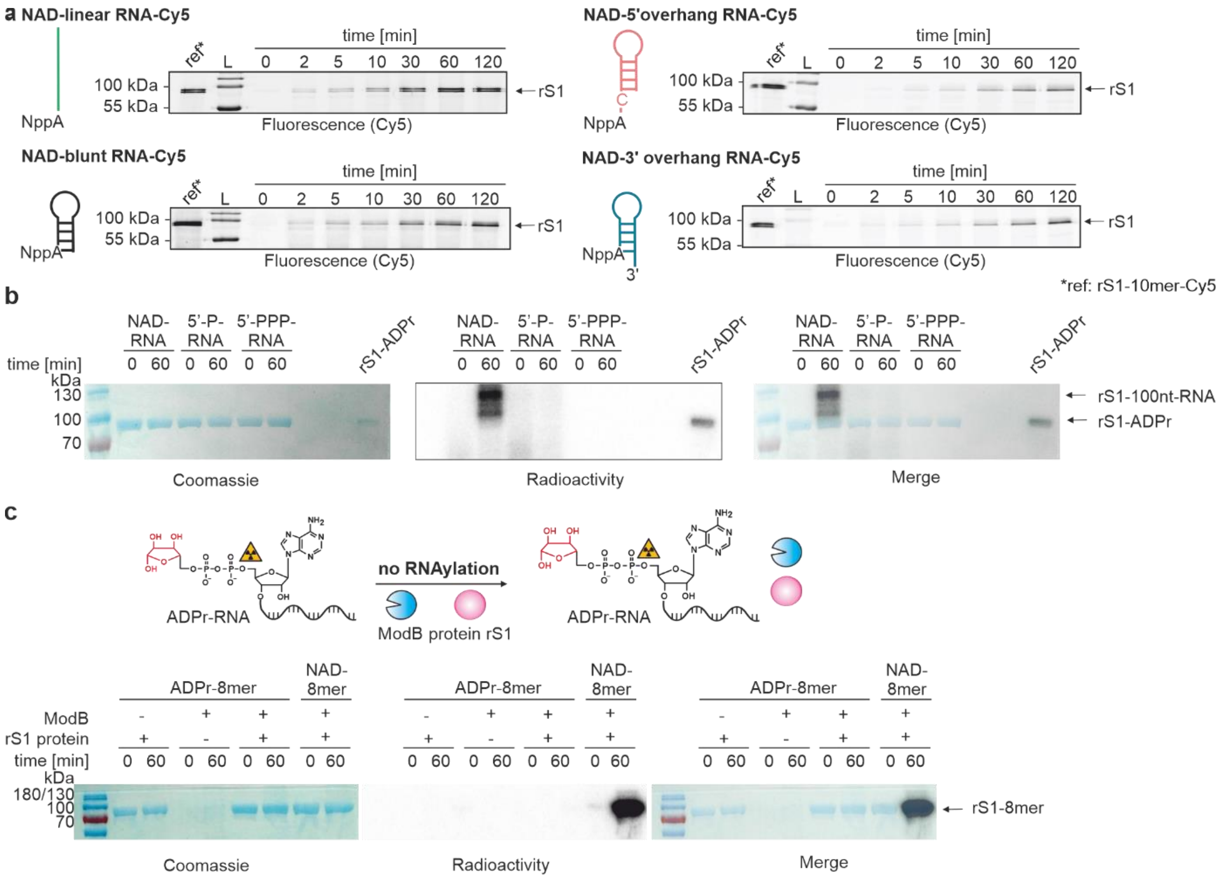


**Extended Data Fig. 3: Characterisation of ModB mediated RNAylation *in vitro*. a**, Analysis of the role of RNA secondary structure on RNAylation reaction. Four different 3'-Cy5-labelled NAD (NppA)-capped RNAs were tested including a linear (green) NAD-capped RNA (10mer) and three structured RNAs with either a 3'-overhang (blue), a 5'-overhang (red) or a blunt end (black) (n=3). SDS-PAGE analyses of the time course of RNAylation are shown. Quantification of the signal intensities (Cy5 scan) relative to the reference is shown in Fig. 2c. ModB prefers linear 5'-ends of NAD-capped RNAs. L = ladder. **b**, Analysis of the RNAylation dependency on the presence of a 5'-NAD-cap of the RNA. 10 % SDS-PAGE analysis of *in vitro* RNAylation of the protein rS1 by ModB in the presence of either 5'-NAD-capped (NAD-RNA), 5'-monophosphate- (5'-P-RNA) or 5'-triphosphate-100nt-RNA (5'-PPP-RNA) (n=2). RNAylation with radiolabelled RNA is detected by radioactivity scan and protein load visualised by Coomassie staining. *In vitro* RNAylation of rS1 is only observed in the presence of NAD-RNA. RNAylated rS1 cannot be detected by Coomassie staining due to low sensitivity. **c**, Characterisation of ADPr-RNA (which is lacking the nicotinamide moiety compared to NAD-RNA) as a substrate for ModB (n=2). As a positive control, NAD-8mer was applied. All reactions were analysed by 12 % SDS-PAGE. RNAylation with radiolabelled RNA is detected by radioactivity scan and protein load visualised by Coomassie staining. ADPr-RNA is not accepted as a substrate for ModB-catalysed RNAylation *in vitro*.

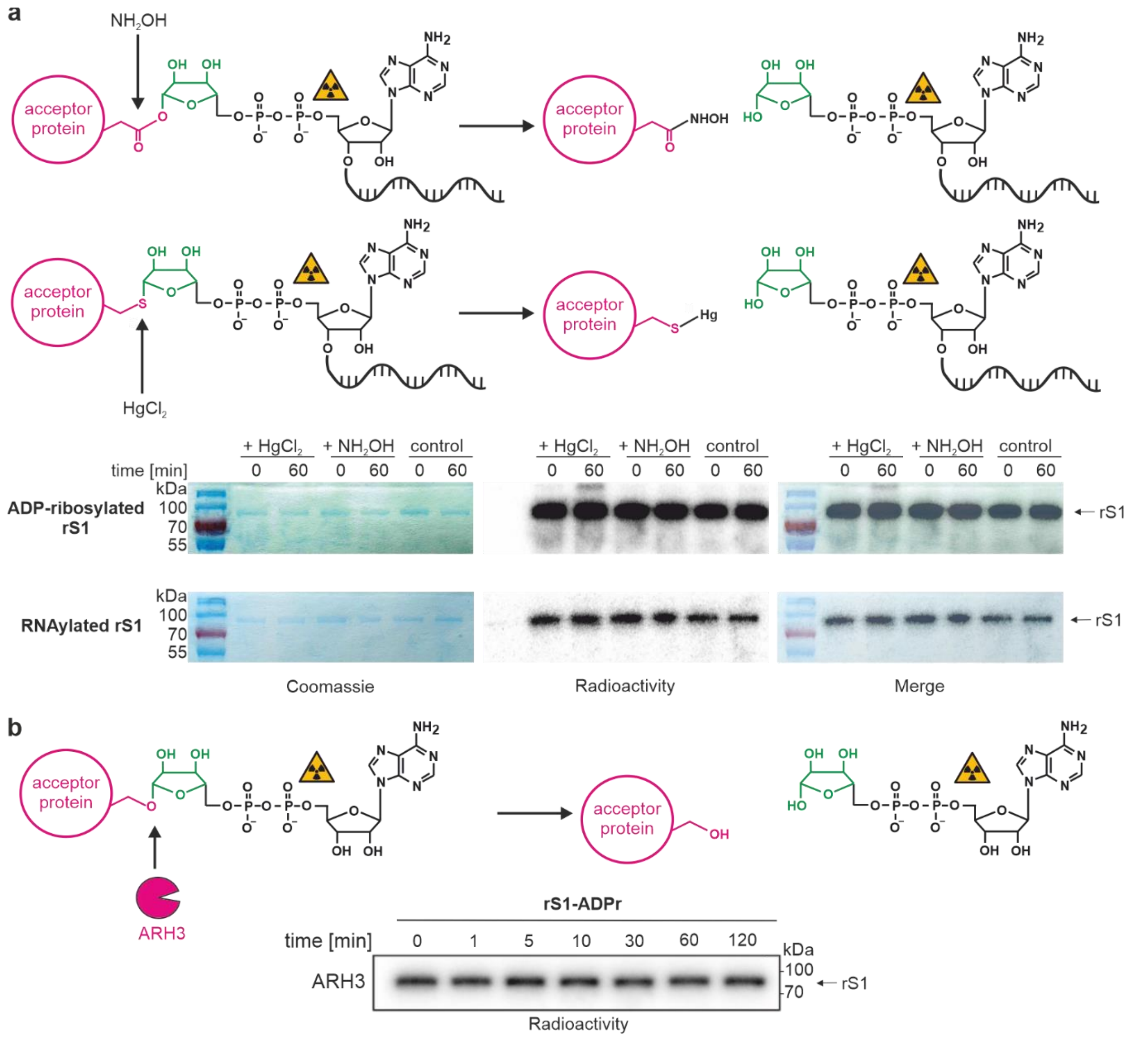


**Extended Data Fig. 4: Specific removal of RNAylation using chemical and enzymatic treatments. a**, Different ADP-ribose-protein linkages have been shown to be either stable or unstable in the presence of $HgCl_2$ and neutral hydroxylamine ($NH_2OH$), which represents a relatively straightforward and fast approach to identify ADP-ribosylation sites. Treatment with $NH_2OH$ hydrolyses linkages between glutamate/aspartate and ADP-ribose. $HgCl_2$ specifically cleaves thiol-glycosidic bonds. ADP-ribosylated and RNAylated protein rS1 were treated with $NH_2OH$ or $HgCl_2$. The removal of ADPr or RNA by these chemicals would result in a decrease of the radioactive signal of protein rS1. All samples were analysed by 12 % SDS-PAGE, stained in Coomassie (protein loading control) and RNAylation assessed as radioactivity. A decrease of the radioactive signal in comparison to the control (untreated) was not determined (n=1). **b**, *in vitro* time course of the stability of rS1 ADP-ribosylation in the presence of ARH3 analysed by 12 % SDS-PAGE (n=1). The autoradiography scan is presented. ARH3 did not remove the ADP-ribosylation.

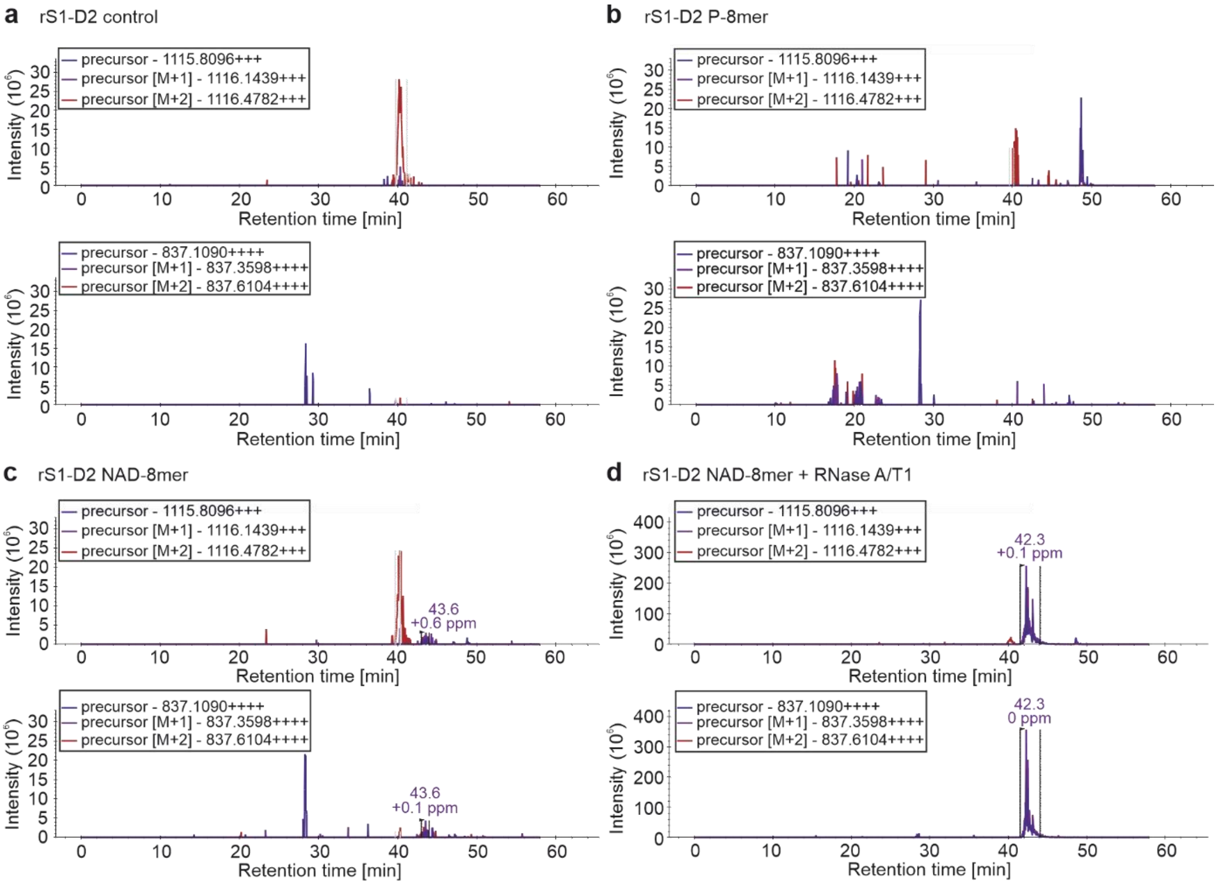


**Extended Data Fig. 5: Ion chromatograms of unmodified rS1 and *in vitro* RNAylated rS1 extracted from LC-MS/MS data.** Extracted ion chromatograms (XICs) for triply and quadruply charged precursor ions (monoisotopic masses 1115.8096 and 837.1090, respectively). XICs were extracted using Skyline [2], an open source document editor for creating and analysing targeted proteomics experiments. The masses correspond to an rS1 peptide AFLPGSLVDVRPVRDTLHLEGK with an attached ADPr-cytidine. Recombinant S1 domain 2 was *in vitro* incubated with ModB and one of the following components: **a**, no other supplements, **b**, uncapped RNA-8mer, **c**, NAD-RNA-8mer, **d**, NAD-RNA-8mer treated with RNase A and T1 (results in ADPr-cytidine adducts). An elution peak at 42.3 min is observable in d and corresponds to the peptide modified with ADPr-cytosine. Spurious intensities can be observed in c and might represent a degradation product. **a** and **b** show only background/contaminant peaks. A contaminant peak at 40 min can be also observed in d (consider the difference in the intensity scale between **d** and **a**-**c**).

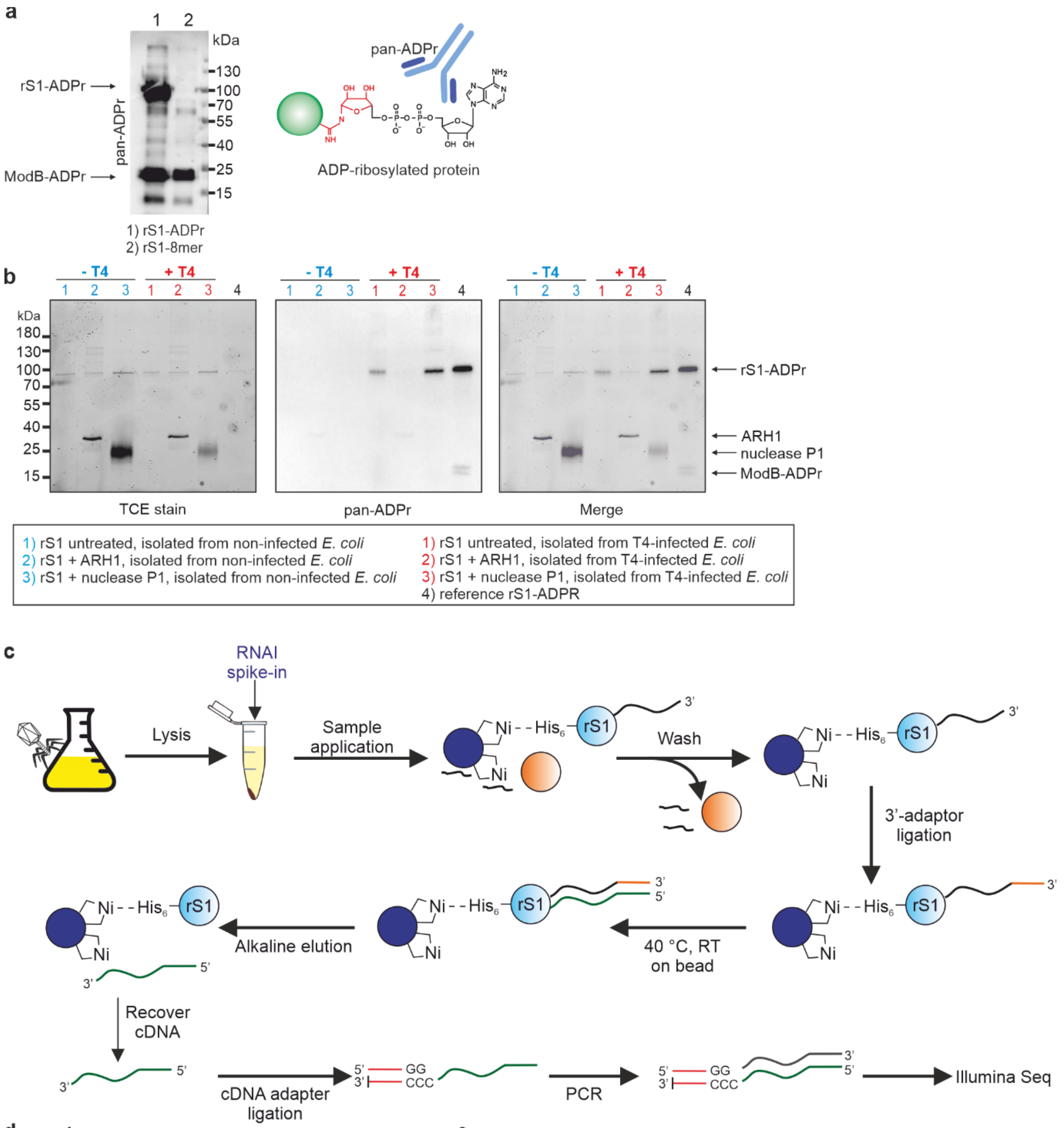


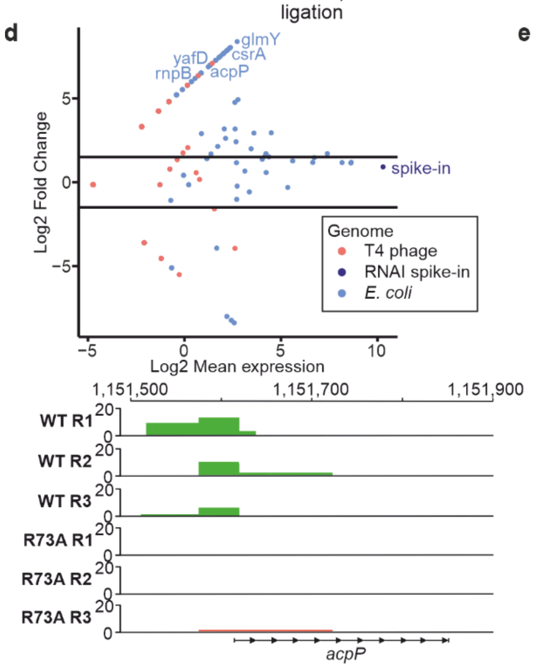


**e**

| Gene | RNAylomeSeq | | | qPCR | | | Genome | Read distribution | NAD-RNA |
|---|---|---|---|---|---|---|---|---|---|
| | R1 | R2 | R3 | R1 | R2 | R3 | | | |
| *acpP* | + | + | + | + | + | + | *E. coli* | 5'-UTR and 5'-fragment | Cahova et al., 2015 Zhang et al., 2021 |
| *gadY* | + | - | - | + | - | + | *E. coli* | full coverage | Cahova et al., 2015 |
| *rob* | + | - | - | n.d. | n.d. | n.d. | *E. coli* | 5'-UTR and 5'-fragment | Zhang et al., 2021 |
| *ipIII* | + | - | - | n.d. | n.d. | n.d. | T4 phage | within gene body | n.d. |
| *mcaS* | - | + | - | - | + | - | *E. coli* | full coverage | Cahova et al., 2015 |
| *glmY* | - | + | - | n.d. | n.d. | n.d. | *E. coli* | full coverage | Cahova et al., 2015 Zhang et al., 2021 |
| *oxyS* | - | + | - | - | + | - | *E. coli* | full coverage | Cahova et al., 2015 |
| *a-gt* | - | - | + | n.d. | n.d. | n.d. | T4 phage | within gene body | n.d. |
| *rnaC* | - | - | + | - | - | (+) | T4 phage | full coverage and 3'-UTR | n.d. |

**Extended Data Fig. 6: *In vivo* characterisation of the RNAylation by Western blot and RNAylomeSeq. a**, Analysis of the substrate specificity of the pan-ADPr antibody. *In vitro* prepared ADP-ribosylated or RNAylated protein rS1 was applied to evaluate the specificity of the antibody (n=3). The pan-ADPr antibody detected ADP-ribosylated proteins rS1 and ModB (lane 1). In contrast, RNAylated rS1 is not detected by pan-ADPr (lane 2). However, a signal for ADP-ribosylated ModB was observed due to self-ADP-ribosylation in its expression host *E. coli* (lane 2). **b**, Quantification of RNAylation using the combination of nuclease P1 digest and detection of protein-linked ADP-ribose by Western blot. Visualisation of protein load by TCE stain. Removal of the ADP-ribose signal by ARH1 treatment. pan-ADPr signals for ADP-ribosylated rS1 were normalised to corresponding band intensities in the TCE stain. The corresponding bar chart is shown in Fig. 4 (n=3). **c,** Schematic illustration of the RNAylomeSeq protocol: Identification of RNAylated RNAs which are covalently attached to rS1 *in vivo*. Briefly, endogenously His-tagged rS1 is isolated from T4 phage infected *E. coli* with Ni-NTA beads. A spike-in - rS1 domain 2 RNAylated with NAD-RNAI - (RNAI spike-in) is added to the lysate which is meant to be enriched via the RNAylomeSeq workflow. rS1 captured on Ni-NTA beads is intensively washed with 8 M urea in order to remove RNA non-covalently bound to rS1. Similar to NAD captureSeq [3], an RNA 3'-adapter is ligated to covalently linked RNAs and RNA is reverse transcribed "on-bead". cDNA is then eluted by alkaline digest of RNA and an additional adapter is ligated to the 3'-terminus of the cDNA. cDNA is amplified by PCR and sequenced by next-generation sequencing (Illumina). Importantly, the RNAI spike-in is not meant to be enriched in any sample but rather to be found in each sample in similar amounts. Thereby, read counts can be normalised to the RNAI counts in each sample allowing for their comparison. **d,** MA plot showing RNAs enriched in the T4 phage WT infected sample compared to T4 phage ModB R73A, G74A control identified by RNAylomeSeq for replicate 2 (total of n=3 biological replicates). Read counts per sample have been normalised to RNAI spike-in read counts which serves as an enrichment control for each sample. Thus, RNAI is not found enriched comparing T4 WT and T4 ModB R73A, G74A. Mean expression values (T4 WT and T4 ModB R73A, G74A condition) have been normalised by Log2 (x-axis). Read coverage on identified RNAylated RNAs as analysed in IGV is exemplarily shown for *acpP* in the lower panel depicting reads in T4 WT samples (green) vs. T4 ModB R73A, G74A samples (red). **e,** Selected hits of RNAs identified by RNAylomeSeq comparing T4 phage WT and T4 ModB R73A, G74A. Enrichments have been further validated on cDNA level by qPCR. +: enriched; -: not enriched; (+): enriched, but Log2 fold change <= 1; n.d.: not defined.

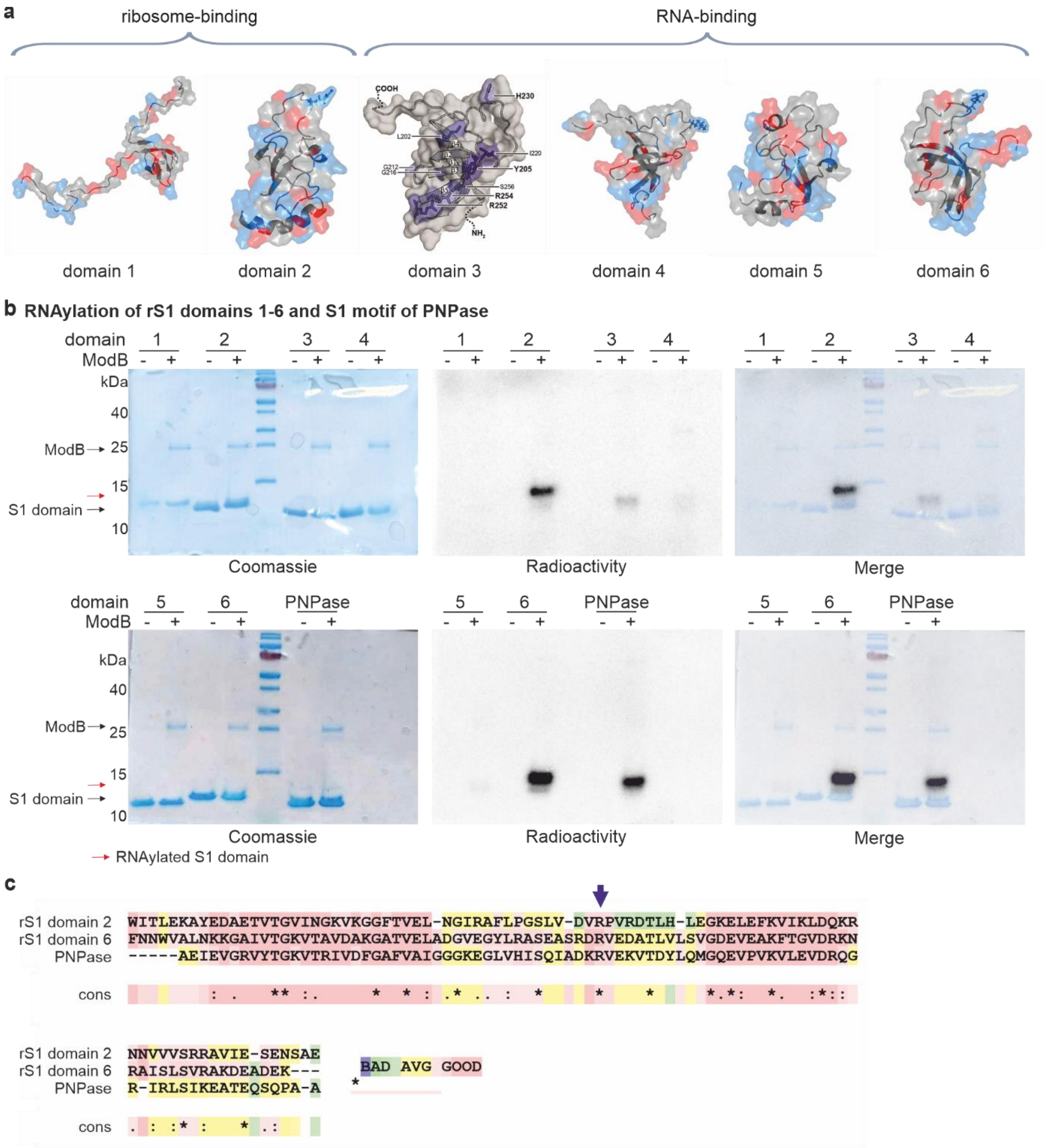


**Extended Data Fig. 7: RNAylation of rS1 domains D1 – D6 and S1 motif of PNPase by ModB *in vitro*. a**, Illustration of the rS1-motifs of rS1 based on crystal structures (PDB) of domains 1 (2MFI), 2 (2MFL), 4 (2KHI), 5 (5XQ5) and 6 (2KHJ) as well as an NMR structure of domain 3 [4]. **b**, *in vitro* RNAylation of S1 domains and PNPase by ModB using a $^{32}$P-NAD-8mer. ModB and S1 domains (D1-6) are marked with black arrows. RNAylated rS1 domains, characterised by a significant shift compared to the non-modified proteins, are highlighted with red arrows. n=2 of biologically independent replicates. Reactions were analysed by 16 % Tricine-SDS-PAGE, stained in Coomassie and RNAylation recorded by autoradiography imaging (Radioactivity). **c,** Local alignment of rS1 D2 and D6 as well as the S1 domain of PNPase using T-coffee expresso [5]. R139 of D2 (highlighted with an arrow) is conserved in PNPase and D6.

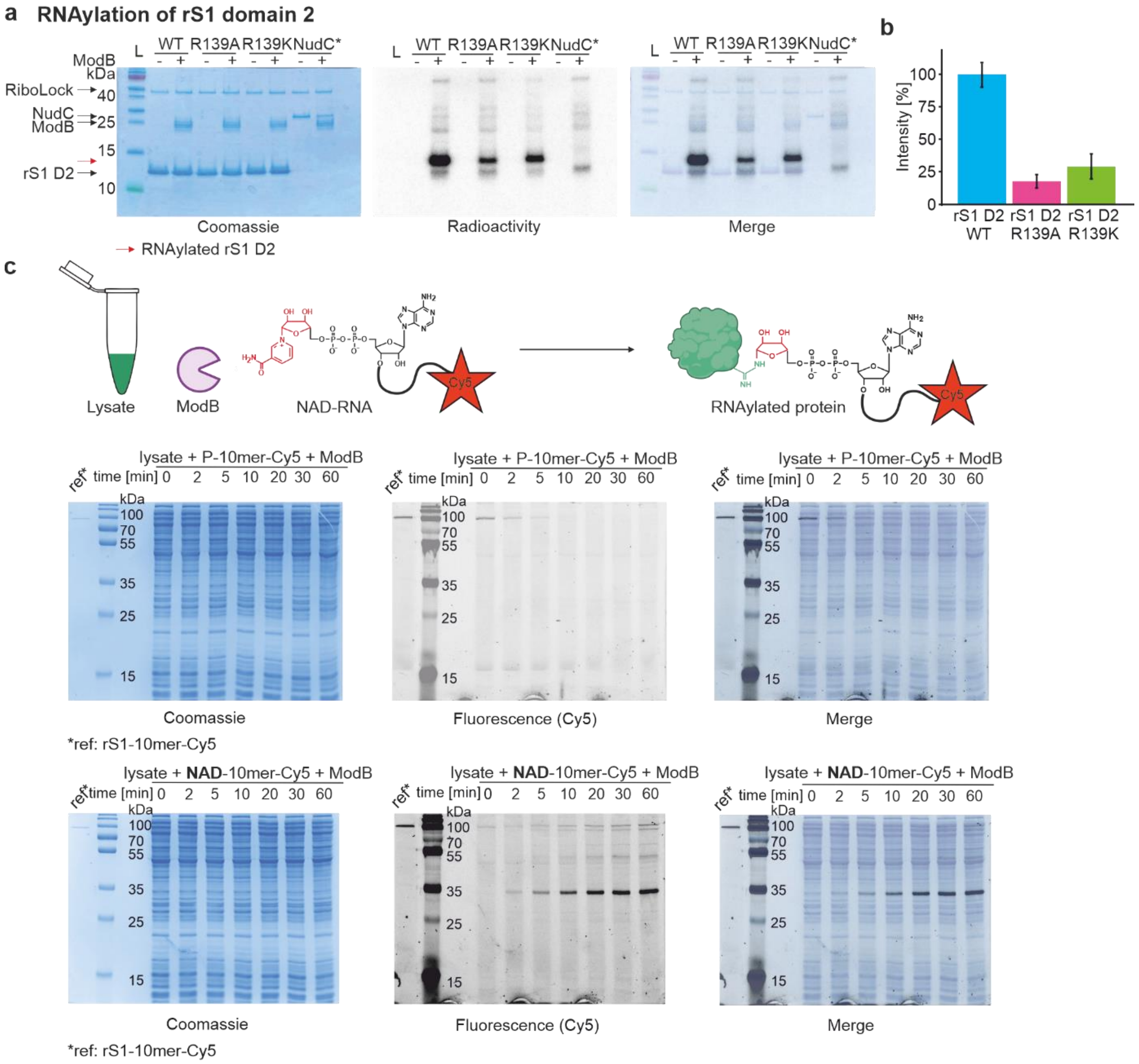


**Extended Data Fig. 8: Characterisation and identification of RNAylation target proteins of ModB. a**, Analysis of the *in vitro* RNAylation of rS1 domain 2 and its mutants R139A and R139K by 16 % Tricine-SDS-PAGE. An inactive NudC mutant (NudC*; V157A, E174A, E177A, E178A) was used as a negative control (n=2). Radioactivity indicates RNAylation, Coomassie scan visualises protein load. **b**, Quantification of relative intensities of RNAylation of rS1 domain 2 and its mutants R139A and R139K based on radioactivity in 16 % Tricine-SDS-PAGE analysis. n=2 of biologically independent replicates. **c**, RNAylation of *E. coli* cell lysate by ModB using 3'-Cy5-labelled NAD-RNA (schematically shown in upper panel). A time course of *E. coli* cell lysate RNAylation by ModB in the presence of either a 5'-monophosphorylated RNA 10mer (P-10mer-Cy5, middle panel) or 5'-NAD-capped RNA 10mer (NAD-10mer-Cy5, lower panel), each with a 3'-fluorescent (Cy5) label. rS1 RNAylated with an NAD-10mer-Cy5 is applied as a reference (ref). The time course of lysate RNAylation was analysed by 12 % SDS-PAGE, protein visualised by Coomassie staining and RNAylation recorded via fluorescence (Cy5). n=3 of biologically independent replicates.

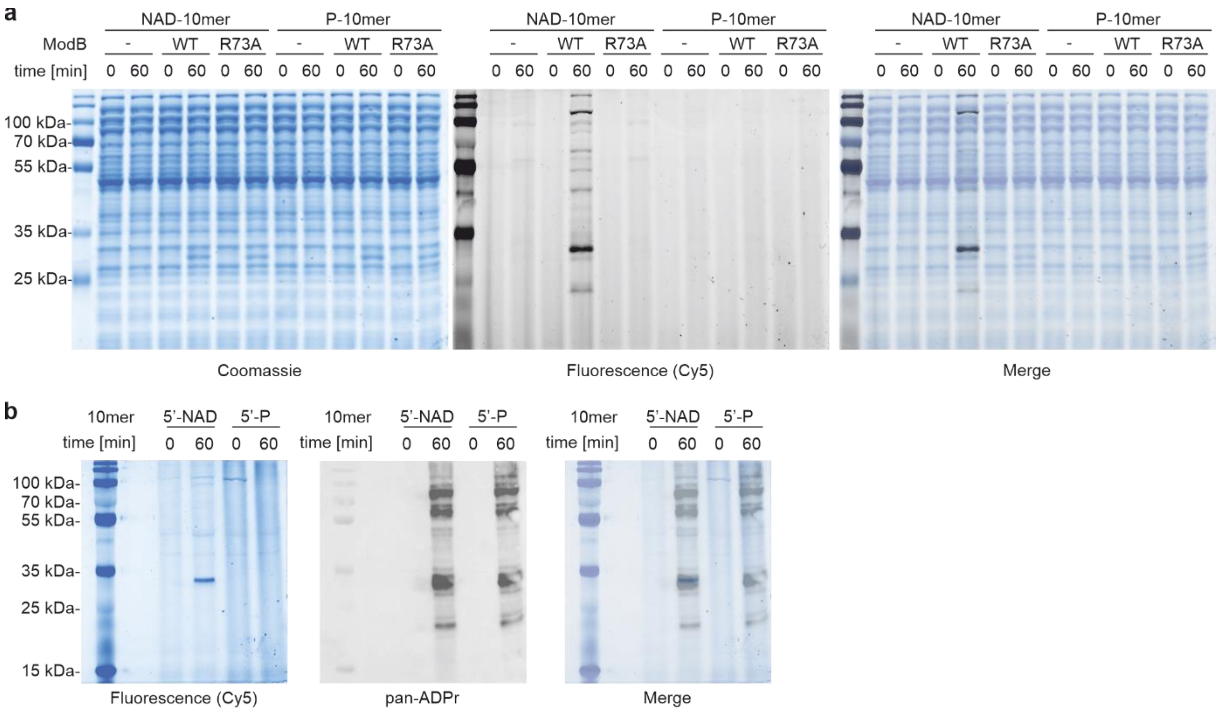


**Extended Data Fig. 9: Characterisation of the specificity of ModB-mediated RNAylation in *E. coli* lysates. a**, RNAylation of *E. coli* cell lysate in the presence of ModB WT or inactive ModB R73A, G73A or the absence of ModB using 3'-Cy5-labelled NAD-10mer or P-10mer. Time point 0 shows lysate before addition of ModB, 60 min shows RNAylation after 60 minutes incubation with ModB. Reactions were analysed by 12 % SDS-PAGE, protein visualised by Coomassie staining and RNAylation recorded via fluorescence (Cy5). n=2 of biologically independent replicates. **b**, Samples from lysate RNAylation with Cy5-labelled 5'-NAD- or 5'-P-10mer (as presented in Extended Data Fig. 8c) before addition of ModB (0 min) and after 60 minutes incubation in the presence of ModB (60 min) were analysed by 10 % SDS-PAGE and RNAylation monitored by fluorescence (Cy5). Subsequently, Western blotting was performed and ADP-ribosylation was detected using pan-ADPr binding reagent (MABE1016). n=2 of biologically independent replicates. Different band patterns were observed for ModB-mediated RNAylation and ADP-ribosylation in *E. coli* lysates indicating a distinct target specificity of ModB for RNAylation.

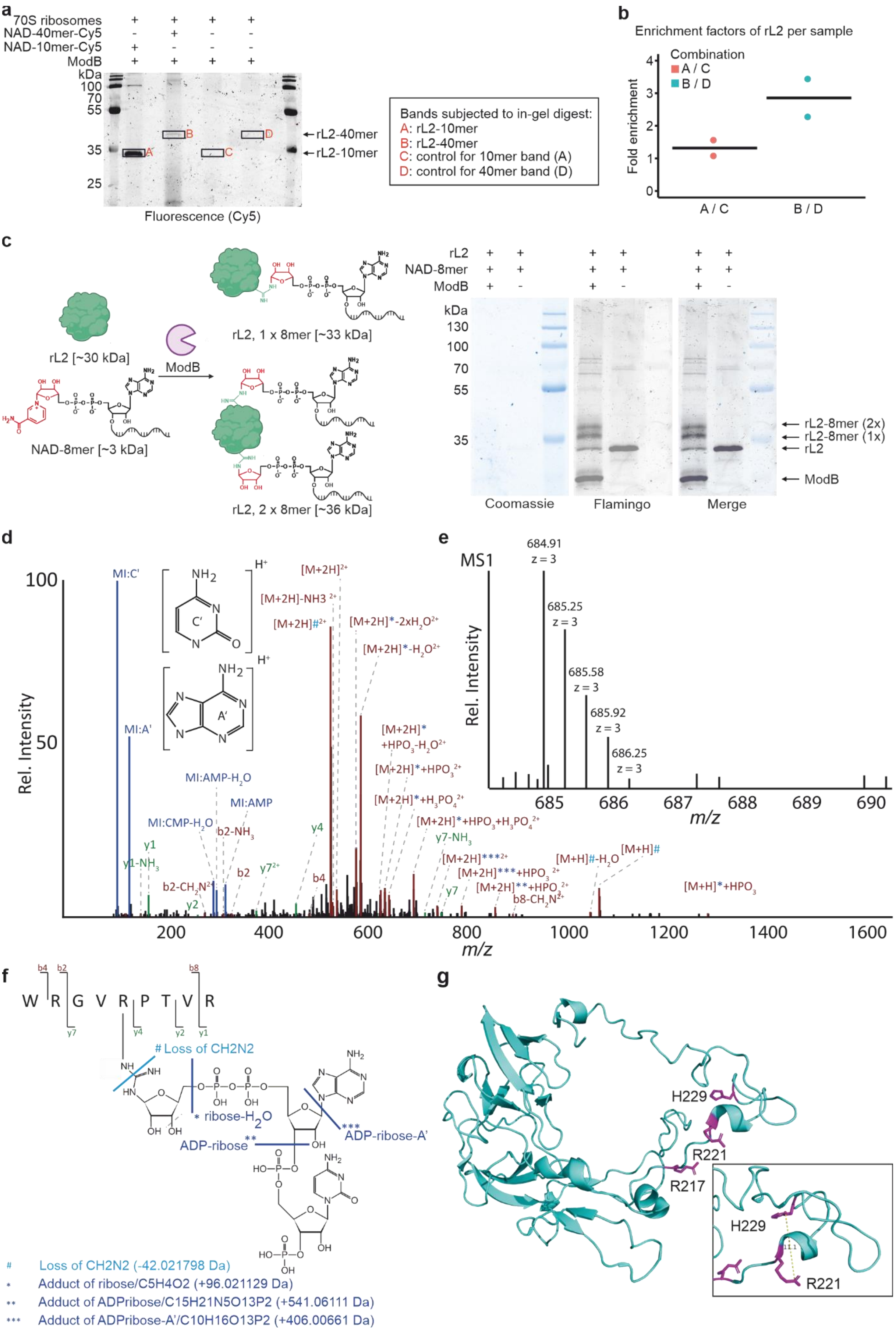

**Extended Data Fig. 10: Scope of ModB RNAylation targets in *E. coli*. a**, RNAylation of *E. coli* ribosomes by ModB. RNAylated protein is shifted upon incubation with NAD-40mer compared to NAD-10mer which itself increases protein weight by approx. 3 kDa. Relative enrichment of RNAylated target protein was assessed by subjecting RNAylated protein bands and respective control bands generated in the absence of RNA to in-gel digest and LC-MS/MS analysis (n=2). **b**, Plot of the enrichment of fractional spectral counts for 50S ribosomal protein L2 (rL2) based on in-gel-digest and LC-MS/MS analysis presented in a. Enrichment is calculated for RNAylation with NAD-10mer (A/C) or NAD-40mer (B/D), relative to the respective, non-RNAylated control bands based on spectral counts from Scaffold (n=2). **c**, Analysis of the *in vitro* RNAylation of rL2 by ModB in the presence of NAD-8mer. RNAylated rL2 proteins have reduced electrophoretic mobility during SDS-PAGE. Protein was visualised by fluorescent protein stain (Flamingo) and protein ladder visualised by Coomassie staining. Signals were quantified using ImageLab indicating that about 80 % of rL2 is RNAylated by ModB *in vitro* (n=3). Band patterns indicated that rL2 can be RNAylated once or even twice *in vitro*. **d-f,** Tandem MS-based identification of RNAylated rL2 peptide. **d,** MS/MS fragment ion spectrum (spectrum ID: 8679) of RNAylated rL2 peptide WRGVRPTVR carrying ADP-ribose plus cytidine-monophosphate and a 3'-phosphate group. The spectrum shows marker ions of adenine (A') and cytosine (C') as well as AMP and CMP. The precursor ion ($[M+xH]^{x+}$) is detected unshifted, shifted by the mass of ADP-ribose (*) and by ADP-ribose with adenine loss (**). Also, precursor ions show a specific loss of 42.021798 Da, which can be explained by a loss of $CH_2N_2$ at the modified arginine. **e,** Isotopic peak pattern of the precursor ion shown in **d**, as detected in the corresponding MS precursor ion scan. **f,** Schematic sequence and RNA adduct representation of the RNAylated peptide shown in **d** and **e** including annotations of fragment ions. The fragmentation products observed in the MS/MS spectrum, shown in **d**, of the ADP-ribose+CMP+3'-phosphate adduct are indicated in the structure by light blue (mass loss) and dark blue (mass adducts) lines. **g,** Selected RNAylated residues of rL2 identified by LC-MS/MS. The catalytically important H229 is 11.1 Å apart from R221. rL2 structure derived from a 1.98 Å cryo-EM structure (7K00) [6].

**Extended Data Table 1: ADP-ribosylation of endogenously His-tagged rS1 during T4 phage infection.** MaxQuant intensities are presented for T4 phage-infected and uninfected samples in triplicates (n=3). R139/R142 located in rS1 domain 2 and R485/R487 in rS1 domain 6 appear as ADP-ribosylation sites on rS1 *in vivo* in all three replicates. The ratio comparing intensity of ADP-ribosylated and unmodified species of the same peptide is computed for each sample and peptide.

| Arginine (domain) | Sequence | Modifications | Intensity uninfected replicate 1 | Intensity uninfected replicate 2 | Intensity uninfected replicate 3 | Intensity T4 infected replicate 1 | Intensity T4 infected replicate 2 | Intensity T4 infected replicate 3 |
|---|---|---|---|---|---|---|---|---|
| R139/R142 (rS1 domain 2) | AFLPGSLVDVRPVRDTLHLEGK | ADP-ribosyl | 0.00E+00 | 0.00E+00 | 0.00E+00 | 2.72E+08 | 2.50E+08 | 3.61E+08 |
| | AFLPGSLVDVRPVRDTLHLEGK | Unmodified | 1.41E+11 | 1.48E+11 | 3.50E+10 | 4.13E+09 | 1.23E+10 | 2.64E+10 |
| | **Ratio: intensity of ADP-ribosylated relative to unmodified peptide** | **ADP-ribosyl/ unmodified** | 0.00E+00 | 0.00E+00 | 0.00E+00 | **6.60E-02** | **2.03E-02** | **1.37E-02** |
| R485/R487 (rS1 domain 6) | ASEASRDRVEDATLVLSVGDEVEAK | ADP-ribosyl | 0.00E+00 | 0.00E+00 | 0.00E+00 | 3.51E+07 | 2.30E+07 | 3.50E+07 |
| | ASEASRDRVEDATLVLSVGDEVEAK | Unmodified | 2.37E+10 | 6.59E+10 | 2.36E+10 | 7.84E+08 | 6.52E+08 | 1.23E+10 |
| | **Ratio: intensity of ADP-ribosylated relative to unmodified peptide** | **ADP-ribosyl/ unmodified** | 0.00E+00 | 0.00E+00 | 0.00E+00 | **4.47E-02** | **3.52E-02** | **2.83E-03** |
| R19 (rS1 domain 1) | EIETRPGSIVR | ADP-ribosyl | 0.00E+00 | 0.00E+00 | 0.00E+00 | 7.73E+06 | 0.00E+00 | 0.00E+00 |
| | EIETRPGSIVR | Unmodified | 5.76E+10 | 7.17E+10 | 7.30E+10 | 4.47E+10 | 3.06E+10 | 2.19E+10 |
| | **Ratio: intensity of ADP-ribosylated relative to unmodified peptide** | **ADP-ribosyl/ unmodified** | 0.00E+00 | 0.00E+00 | 0.00E+00 | **1.73E-04** | **0.00E+00** | **0.00E+00** |

**Extended Data Table 2: ADP-ribosylation of rS1-WT, -R139K and -R139A during T4 phage infection.** MaxQuant intensities are presented for T4 phage-infected and uninfected samples in triplicates (n=3) only for the respective peptide of the R139 mutation site which is expected for the respective rS1 version. ADP-ribosylation of the peptide in rS1 is observed *in vivo* in all three replicates. However, ADP-ribosylation at position 139 is abolished by R139A or R139K mutations (mutation indicated in red). The intensity of ADP-ribosylated peptide relative to the intensity of the corresponding unmodified peptide species is at least 3-fold reduced upon R139 mutation. One may speculate that R142 is nevertheless ADP-ribosylated in the mutated rS1 proteins but overall ADP-ribosylation yield at the peptide may be reduced as the potentially predominant ADP-ribosylation site (R139) is not available for modification.

| rS1 R139K – peptide AFLPGSLVDVKPVRDTLHLEGK | | | | | | | |
|---|---|---|---|---|---|---|---|
| **Sequence** | **Modifications** | **Intensity rS1-R139K uninfected replicate 1** | **Intensity rS1-R139K uninfected replicate 2** | **Intensity rS1-R139K uninfected replicate 3** | **Intensity rS1-R139K T4 infected replicate 1** | **Intensity rS1-R139K T4 infected replicate 2** | **Intensity rS1-R139K T4 infected replicate 3** |
| AFLPGSLVDV**K**PVRDTLHLEGK | ADP-ribosyl | 0.00E+00 | 0.00E+00 | 0.00E+00 | 1.61E+09 | 4.87E+09 | 1.50E+09 |
| AFLPGSLVDV**K**PVRDTLHLEGK | Unmodified | 2.27E+10 | 4.88E+10 | 4.70E+10 | 4.17E+10 | 3.52E+10 | 6.60E+10 |
| **Ratio: intensity of ADP-ribosylation relative to unmodified peptide** | | **0.00E+00** | **0.00E+00** | **0.00E+00** | **3.86E-02** | **1.38E-01** | **2.27E-02** |

| rS1 R139A – peptide AFLPGSLVDVAPVRDTLHLEGK | | | | | | | |
|---|---|---|---|---|---|---|---|
| **Sequence** | **Modifications** | **Intensity rS1-R139A uninfected replicate 1** | **Intensity rS1-R139A uninfected replicate 2** | **Intensity rS1-R139A uninfected replicate 3** | **Intensity rS1-R139A T4 infected replicate 1** | **Intensity rS1-R139A T4 infected replicate 2** | **Intensity rS1-R139A T4 infected replicate 3** |
| AFLPGSLVDV**A**PVRDTLHLEGK | ADP-ribosyl | 0.00E+00 | 0.00E+00 | 0.00E+00 | 5.27E+09 | 8.05E+09 | 9.17E+09 |
| AFLPGSLVDV**A**PVRDTLHLEGK | Unmodified | 8.33E+10 | 9.93E+10 | 7.37E+10 | 7.50E+10 | 7.96E+10 | 7.32E+10 |
| **Ratio: intensity of ADP-ribosylation relative to unmodified peptide** | | **0.00E+00** | **0.00E+00** | **0.00E+00** | **7.02E-02** | **1.01E-01** | **1.25E-01** |

| rS1 WT – peptide AFLPGSLVDVRPVRDTLHLEGK | | | | | | | |
|---|---|---|---|---|---|---|---|
| **Sequence** | **Modifications** | **Intensity rS1-WT uninfected replicate 1** | **Intensity rS1-WT uninfected replicate 2** | **Intensity rS1-WT uninfected replicate 3** | **Intensity rS1-WT T4 infected replicate 1** | **Intensity rS1-WT T4 infected replicate 2** | **Intensity rS1-WT T4 infected replicate 3** |
| AFLPGSLVDV**R**PVRDTLHLEGK | ADP-ribosyl | 0.00E+00 | 1.17E+08 | 0.00E+00 | 1.27E+10 | 1.12E+10 | 5.59E+09 |
| AFLPGSLVDV**R**PVRDTLHLEGK | Unmodified | 1.12E+10 | 3.34E+09 | 2.27E+10 | 6.85E+09 | 2.28E+10 | 1.51E+10 |
| **Ratio: intensity of ADP-ribosylation relative to unmodified peptide** | | **0.00E+00** | **3.49E-02** | **0.00E+00** | **1.85E+00** | **4.91E-01** | **3.71E-01** |

**Supplementary Table 1: MaxQuant Output for LC-MS/MS analysis of endogenously His-tagged rS1 from T4 phage-infected *E. coli* B strain.** Endogenously His-tagged rS1 was isolated from T4 phage-infected (inf) and -uninfected *E. coli* B strain and subjected to LC-MS/MS analysis in biological triplicates (n=3). Intensities from MaxQuant are only shown for rS1 (1A;modificationSpecificPeptides). ADP-ribosylation is detected only for a small subset of rS1 peptides from T4 phage-infected samples whilst absent in uninfected samples. Predominantly, R139/R142 and R485/R487 were identified as ADP-ribosylation sites in all three replicates. ADP-ribosylated peptides are listed by rS1 domain and with the respective arginine residues in 1B; ADPr peptides. Modifications occur at R485/R487 (domain 6) and R139/R142 (domain 2) in all three replicates. Comparing intensities of ADP-ribosylated and unmodified peptides (1C; ADPr vs. unmodified peptides) shows ratios varying from 1.4 % to 6.6 %. Based on this data, one may speculate that R139/R142 and R485/R487 might be major ADP-ribosylation sites on rS1 *in vivo*. MaxQuant parameters for the presented data are presented in 1D; Parameters MaxQuant.

**Supplementary Table 2: MaxQuant Output for LC-MS/MS analysis of His-tagged rS1-WT, -R139A and -R139K mutants from T4 phage-infected *E. coli*.** MaxQuant Output filtered for rS1 protein and sorted according to ADP-ribosylation (ADP-ribosylwoDP) is presented (2A; modificationSpecificPeptides). T4 phage-infected samples (T4 phage) and -uninfected control (LB control) per rS1 version (WT, R139A or R139K mutant) are presented. A total of three biological replicates (n=3) were analysed per rS1 version. Peptides which were found ADP-ribosylated are listed and are assigned to their respective location (rS1 domain) and the modified arginines in the rS1 protein (2B; ADPr peptides). Intensities of ADP-ribosylated peptides are compared to their unmodified counterpart each by dividing the respective intensities (2C; ADPr vs. unmodified peptides). The peptide AFLPGSLVDVR(K/A)PVRDTLHLEGK is found ADP-ribosylated for rS1 WT, rS1 R139A and R139K only in T4 phage-infected samples across all three replicates. It becomes obvious that for the WT peptide high intensities of the ADP-ribosylated peptide relative to the unmodified peptide are detected across all three T4 phage-infected replicates. For the mutant peptides R139A and R139K, these intensities are at least 3-fold lower. Based on this finding, one may speculate that R139 mutation might reduce ADP-ribosylation of the AFLPGSLVDVRPVRDTLHLEGK peptide in rS1. MaxQuant parameters for the presented data are presented in 2D; Parameters MaxQuant.

**Supplementary Table 3: *In vitro* ADP-ribosylation and RNAylation sites in rS1 protein as identified by LC-MS/MS analysis.** Peptide spectrum match (PSM) information for ADP-ribosylated and/or RNAylated rS1 peptides are given in summarised form (pivot) and as complete output from OpenMS tool RNPxl (PSMs). Results were filtered for 1 % FDR on PSM level and q-values (scores) are given. Spectrum IDs, precursor m/z values, charge states, best localisation of modification within the peptide sequence, localisation score and mass errors (in ppm) are provided in "PSMs" sheet.

**Supplementary Table 4: Genes identified to contribute to the RNAylome by RNAylomeSeq.** An excerpt from the counts table is presented. Hits are calculated based on the mean read counts for each gene among T4 WT and R73A, G74A (MUT) samples for each replicate individually. For a hit, the log2 Fold Change (LFC) between WT and MUT sample is to be greater than 1.5 and the log2 transformed mean expression greater than -0.5. Hits are indicated as "+" for individual replicates. Read distribution for hits is presented in column "IGV" and the existence of corresponding NAD-capped transcripts is indicated in column "NAD-capped RNA?". Raw read counts are shown in WT_R1 – MUT_R3. Some hits are present in all replicates, some in one or two replicates only. Importantly, for the majority of protein_coding and ncRNA genes, reads initiate with an adenosine or contain an adenosine no more than 2 nt away from the read start. tRNA and rRNA (which more likely represent the background) hits are more abundant in replicates 2 and 3. Especially in replicate 3, the fraction of RNAI reads varies comparing WT and MUT samples, which may explain this variation from the background.

**Supplementary Table 5: *In vitro* ADP-ribosylation and RNAylation sites in rL2 protein as identified by LC-MS/MS analysis.** Peptide spectrum match (PSM) information for ADP-ribosylated and/or RNAylated rL2 peptides are given in summarised form (pivot) and as complete output from OpenMS tool RNPxl (PSMs). Results were filtered for 1 % FDR on PSM level and q-values (scores) are given. Spectrum IDs, precursor m/z values, charge states, best localisation of modification within the peptide sequence, localisation score and mass errors (in ppm) are provided in "PSMs" sheet.

**Supplementary information:** contains Supplementary Tables 6 – 9.

**Supplementary Table 6: RNAs used in this study**

| RNA | RNA sequence |
|---|---|
| **8mer** | ACAGUAUU |
| **RNAI** | ACAGUAUUUGGUAUCUGCGCUCUGCUGAAGCCAGUUACCUUCGGAAAAAGAG<br>UUGGUAGCUCUUGAUCCGGCAAACAAACCACCGCUGGUAGCGGUGGUUUUU<br>UUGUU |
| **100nt-RNA (Qβ)** | AUCUUGAUACUACCUUUAGUUCGUUUAAACACGUUCUUGAUAGUAUCUUUU<br>UAUUAACCCAACGCGUAAAGCGUUGAAACUUUGGGUCAAUUUGAUCAUG |
| **10mer-Cy5** | ACAGUAUUUG |
| **2nt-5'overhang-Cy5** | ACAGACUUCGGUCU-Cy5 |
| **3'overhang-Cy5** | AGACUUCGGUCUA-Cy5 |
| **5'P-blunt-Cy5** | AGACUUCGGUCU-Cy5 |
| **linear-Cy5** | AGACUUCGAC-Cy5 |
| **40mer-Cy5** | ACAGUAUUUGGUAUCUGCGCUCUGCUGAAGCCAGUUACUU-Cy5 |

**Supplementary Table 7: Genomic DNA sequence of ARTs, rS1 variants and ADP-ribose hydrolases.** Start codon in italic; thrombin cleavage site in bold; mutations in red and bold; restriction sites underlined.

| Gene [5', 3' restriction site] | DNA sequence |
|---|---|
| **Alt [*NcoI*, *XhoI*]** | <u>CC*ATG*G</u>GAGAACTTATTACAGAATTATTTGACGAAGATACTACTCTTCCAATTACAAACTTAT<br>ATCCAAAGAAGAAAATACCGCAAATTTTTTCAGTTCATGTTGATGATGCAATTGAACAACCAG<br>GCTTTCGTTTATGTACCTATACATCTGGAGGTGATACTAATCGTGATTTAAAGATGGGCGATA<br>AAATGATGCATATTGTTCCTTTTACATTAACTGCTAAAGGTTCAATTGCTAAATTAAAAGGTCT<br>TGGTCCAAGCCCAATTAATTATATCAATTCAGTTTTTACTGTTGCAATGCAAACAATGCGCCA<br>GTATAAAATTGATGCCTGTATGCTCCGTATTCTTAAGTCTAAAACTGCTGGCCAAGCTCGACA<br>AATTCAAGTTATTGCTGATAGACTTATCCGTAGTCGTTCAGGTGGTAGATACGTCCTTCTTAA<br>GGAACTCTGGGATTACGATAAAAAGTATGCATATATTCTTATACATCGCAAAAATGTATCACT<br>AGAAGACATTCCAGGAGTTCCGGAAATTAGTACCGAGCTCTTTACTAAAGTTGAATCGAAGG<br>TCGGTGATGTTTATATCAATAAAGATACTGGGGCTCAAGTAACTAAAAATGAGGCAATTGCA<br>GCATCTATTGCGCAAGAAAATGATAAACGTTCTGACCAAGCTGTAATCGTTAAAGTTAAAATT<br>TCCCGTAGAGCAATTGCGCAAAGTCAGTCATTGGAATCTTCTAGATTTGAAACACCAATGTTT<br>CAAAAATTTGAGGCTTCAGCGGCCGAATTAAATAAACCAGCGGACGCGCCTTTAATTTCTGAT<br>TCTAATGAATTAACGGTAATTTCTACTTCAGGATTTGCACTAGAGAATGCTCTTAGCAGTGTT<br>ACAGCTGGGATGGCATTCAGAGAAGCTTCTATAATTCCTGAAGATAAAGAATCCATTATTAAC<br>GCAGAAATAAAAAATAAAGCTTTAGAAAGATTACGAAAAGAATCTATTACTTCAATAAAAAC |

| | |
|---|---|
| | CTTAGAAACTATTGCTTCTATCGTCGATGATACTTTAGAAAAATATAAGGGTGCTTGGTTTGA<br>AAGAAATATTAACAAACATTCGCATTTAAACCAAGATGCTGCAAATGAGTTAGTACAAAATTC<br>TTGGAATGCAATAAAAACAAAGATTATTCGAAGAGAATTACGTGGATATGCTCTTACCGCTG<br>GATGGTCATTACATCCTATAGTCGAAAATAAAGATTCATCTAAATACACACCAGCGCAAAAAC<br>GCGGAATTCGTGAATACGTAGGTTCAGGATATGTAGACATAAATAATGCTCTTTTGGGATTAT<br>ATAATCCAGATGAGCGTACAAGTATTTTGACAGCATCTGACATAGAAAAAGCTATTGATAATT<br>TAGATTCAGCCTTTAAAAATGGTGAACGATTACCAAAAGGTATTACTTTGTATCGTTCACAAC<br>GAATGTTACCTTCAATATACGAAGCAATGGTAAAAAATCGAGTTTTTTATTTTAGAAACTTTGT<br>GTCAACATCATTATATCCAAATATTTTTGGTACTTGGATGACTGATTCATCTATAGGTGTTTTA<br>CCAGACGAAAAGCGTTTAAGCGTTTCTATTGATAAAACTGATGAAGGACTTGTAAATTCTAGC<br>GATAATTTAGTTGGAATTGGATGGGTTATTACTGGGGCTGATAAGGTCAATGTTGTTTTACCC<br>GGTGGAAGTTTAGCGCCTTCAAATGAAATGGAAGTCATTTTGCCACGTGGATTAATGGTCAA<br>AGTTAATAAAATAACCGATGCATCTTACAATGATGGAACAGTTAAAACTAACAACAAGCTTAT<br>TCAAGCTGAAGTTATGACCACAGAAGAACTCACCGAATCGGTAATCTATGACGGAGACCATT<br>TAATGGAAACTGGTGAATTGGTTACAATGACAGGTGATATAGAAGATAGAGTTGACTTTGCA<br>TCATTTGTTTCATCAAATGTTAAACAGAAAGTAGAATCATCTCTTGGAATTATTGCGTCTTGCA<br>TAGATATTGCAAACATGCCTTACAAGTTCGTTCAAGGA**CTGGTGCCGCGCGGCAGC**CTCGAG |
| **ModA [*NcoI*, *XhoI*]** | CC*ATGG*GAAAATACTCAGTAATGCAACTAAAAGATTTTAAAATAAAATCAATGGATGCATCG<br>GTGCGTGCTTCTATTCGTGAAGAATTACTTTCTGAAGGGTTTAATTTATCTGAAATTGAACTTT<br>TAATTCATTGTATTACTAATAAACCAGATGACCATTCTTGGTTAAATGAAATAATCAAATCTCG<br>TTTGGTTCCAAACGATAAACCTCTTTGGAGAGGTGTTCCAGCTGAGACTAAACAAGTATTAAA<br>TCAAGGAATTGATATTATTACATTTGATAAAGTCGTATCAGCTTCATATGATAAAAATATAGC<br>TCTACATTTTGCTTCTGGTTTAGAGTATAACACACAAGTTATTTTTGAATTCAAAGCTCCTATG<br>GTATTCAATTTCCAGGAGTATGCTATAAAAGCTCTACGCTGTAAAGAATACAATCCAAACTTT<br>AAGTTTCCGGATAGTCATCGTTATCGTAATATGGAATTAGTTTCAGATGAACAAGAAGTAATG<br>ATACCAGCTGGAAGTGTATTTAGAATTGCAGATAGATATGAGTATAAAAAGTGTTCAACATA<br>CACTATCTATACTCTTGATTTTGAAGGATTTAATCTA**CTGGTGCCGCGCGGCAGC**CTCGAG |
| **ModB [*NcoI*, *XhoI*]** | CC*ATGG*GAATTATTAATCTTGCAGATGTTGAACAGTTATCTATAAAAGCTGAAAGCGTTGATT<br>TTCAATATGATATGTATAAAAAGGTCTGTGAAAAATTTACTGACTTTGAGCAGTCTGTTCTTTG<br>GCAATGTATGGAAGCCAAAAAGAATGAAGCTCTTCATAAGCATTTAAATGAAATCATTAAAA<br>AGCATTTAACTAAATCGCCTTATCAATTATATCGTGGTATATCAAAATCGACAAAAGAACTCA<br>TTAAAGATTTACAAGTTGGAGAAGTGTTTTCAACGAACAGGGTAGATTCATTTACTACTAGTT<br>TGCATACAGCGTGTTCTTTTTCTTATGCTGAATATTTCACTGAAACAATACTTCGTTTAAAAAC<br>TGATAAAGCTTTTAATTATTCTGACCATATCAGCGATATTATACTTTCTTCTCCTAATACTGAGT<br>TTAAGTACACGTATGAAGATACTGATGGATTAGATTCAGAGCGTACTGATAACTTAATGATG<br>ATTGTGCGTGAACAAGAATGGATGATTCCAATTGGAAAGTATAAAATAACTTCTATTTCAAAA<br>GAAAAATTACACGATTCATTTGGAACATTTAAAGTTTATGATATTGAGGTAGTTGAA**CTGGTG<br>CCGCGCGGCAGC**CTCGAG |
| **ModB R73A, G74A [*NcoI*/*XhoI*]** | *CCATGG*GAATTATTAATCTTGCAGATGTTGAACAGTTATCTATAAAAGCTGAAAGCGTTGATT<br>TTCAATATGATATGTATAAAAAGGTCTGTGAAAAATTTACTGACTTTGAGCAGTCTGTTCTTTG<br>GCAATGTATGGAAGCCAAAAAGAATGAAGCTCTTCATAAGCATTTAAATGAAATCATTAAAA<br>AGCATTTAACTAAATCGCCTTATCAATTATAT**GCGGCA**ATATCAAAATCGACAAAAGAACTCA<br>TTAAAGATTTACAAGTTGGAGAAGTGTTTTCAACGAACAGGGTAGATTCATTTACTACTAGTT |

| | |
|---|---|
| | TGCATACAGCGTGTTCTTTTTCTTATGCTGAATATTTCACTGAAACAATACTTCGTTTAAAAAC<br>TGATAAAGCTTTTAATTATTCTGACCATATCAGCGATATTATACTTTCTTCTCCTAATACTGAGT<br>TTAAGTACACGTATGAAGATACTGATGGATTAGATTCAGAGCGTACTGATAACTTAATGATG<br>ATTGTGCGTGAACAAGAATGGATGATTCCAATTGGAAAGTATAAAATAACTTCTATTTCAAAA<br>GAAAAATTACACGATTCATTTGGAACATTTAAAGTTTATGATATTGAGGTAGTTGAA**CTGGTG**<br>**CCGCGCGGCAGC**CTCGAGCACCACCACCACCACCACTGA |
| **pET28-rS1 [*NcoI*, *XhoI*]** | CC*ATGG*GAACTGAATCTTTGCGGCATGCTCAACTCTTTGAAGAGTCCTTAAAAGAAATCGAA<br>ACCCGCCCGGGTTCTATCGTTCGTGGCGTTGTTGTTGCTATCGACAAAGACGTAGTACTGGTT<br>GACGCTGGTCTGAAATCTGAGTCCGCCATCCCGGCTGAGCAGTTCAAAAACGCCCAGGGCGA<br>GCTGGAAATCCAGGTAGGTGACGAAGTTGACGTTGCTCTGGACGCAGTAGAAGACGGCTTC<br>GGTGAAACTCTGCTGTCCCGTGAGAAAGCTAAACGTCACGAAGCCTGGATCACGCTGGAAAA<br>AGCTTACGAAGATGCTGAAACTGTTACCGGTGTTATCAACGGCAAAGTTAAGGGCGGCTTCA<br>CTGTTGAGCTGAACGGTATTCGTGCGTTCCTGCCAGGTTCTCTGGTAGACGTTCGTCCGGTGC<br>GTGACACTCTGCACCTGGAAGGCAAAGAGCTTGAATTTAAAGTAATCAAGCTGGATCAGAAG<br>CGCAACAACGTTGTTGTTTCTCGTCGTGCCGTTATCGAATCCGAAAACAGCGCAGAGCGCGA<br>TCAGCTGCTGGAAAACCTGCAGGAAGGCATGGAAGTTAAAGGTATCGTTAAGAACCTCACTG<br>ACTACGGTGCATTCGTTGATCTGGGCGGCGTTGACGGCCTGCTGCACATCACTGACATGGCC<br>TGGAAACGCGTTAAGCATCCGAGCGAAATCGTCAACGTGGGCGACGAAATCACTGTTAAAGT<br>GCTGAAGTTCGACCGCGAACGTACCCGTGTATCCCTGGGCCTGAAACAGCTGGGCGAAGATC<br>CGTGGGTAGCTATCGCTAAACGTTATCCGGAAGGTACCAAACTGACTGGTCGCGTGACCAAC<br>CTGACCGACTACGGCTGCTTCGTTGAAATCGAAGAAGGCGTTGAAGGCCTGGTACACGTTTC<br>CGAAATGGACTGGACCAACAAAAACATCCACCCGTCCAAAGTTGTTAACGTTGGCGATGTAG<br>TGGAAGTTATGGTTCTGGATATCGACGAAGAACGTCGTCGTATCTCCCTGGGTCTGAAACAG<br>TGCAAAGCTAACCCGTGGCAGCAGTTCGCGGAAACCCACAACAAGGGCGACCGTGTTGAAG<br>GTAAAATCAAGTCTATCACTGACTTCGGTATCTTCATCGGCTTGGACGGCGGCATCGACGGCC<br>TGGTTCACCTGTCTGACATCTCCTGGAACGTTGCAGGCGAAGAAGCAGTTCGTGAATACAAA<br>AAAGGCGACGAAATCGCTGCAGTTGTTCTGCAGGTTGACGCAGAACGTGAACGTATCTCCCT<br>GGGCGTTAAACAGCTCGCAGAAGATCCGTTCAACAACTGGGTTGCTCTGAACAAGAAAGGC<br>GCTATCGTAACCGGTAAAGTAACTGCAGTTGACGCTAAAGGCGCAACCGTAGAACTGGCTGA<br>CGGCGTTGAAGGTTACCTGCGTGCTTCTGAAGCATCCCGTGACCGCGTTGAAGACGCTACCC<br>TGGTTCTGAGCGTTGGCGACGAAGTTGAAGCTAAATTCACCGGCGTTGATCGTAAAAACCGC<br>GCAATCAGCCTGTCTGTTCGTGCGAAAGACGAAGCTGACGAGAAAGATGCAATCGCAACTGT<br>TAACAAACAGGAAGATGCAAACTTCTCCAACAACGCAATGGCTGAAGCTTTCAAAGCAGCTA<br>AAGGCGAG**CTGGTGCCGCGCGGCAGC**CTCGAG |
| **pET28-rS1 R139A [*NcoI*, *XhoI*]** | CC*ATGG*GAACTGAATCTTTGCGGCATGCTCAACTCTTTGAAGAGTCCTTAAAAGAAATCGAA<br>ACCCGCCCGGGTTCTATCGTTCGTGGCGTTGTTGTTGCTATCGACAAAGACGTAGTACTGGTT<br>GACGCTGGTCTGAAATCTGAGTCCGCCATCCCGGCTGAGCAGTTCAAAAACGCCCAGGGCGA<br>GCTGGAAATCCAGGTAGGTGACGAAGTTGACGTTGCTCTGGACGCAGTAGAAGACGGCTTC<br>GGTGAAACTCTGCTGTCCCGTGAGAAAGCTAAACGTCACGAAGCCTGGATCACGCTGGAAAA<br>AGCTTACGAAGATGCTGAAACTGTTACCGGTGTTATCAACGGCAAAGTTAAGGGCGGCTTCA<br>CTGTTGAGCTGAACGGTATTCGTGCGTTCCTGCCAGGTTCTCTGGTAGACGTT**GCC**CCGGTGC<br>GTGACACTCTGCACCTGGAAGGCAAAGAGCTTGAATTTAAAGTAATCAAGCTGGATCAGAAG<br>CGCAACAACGTTGTTGTTTCTCGTCGTGCCGTTATCGAATCCGAAAACAGCGCAGAGCGCGA<br>TCAGCTGCTGGAAAACCTGCAGGAAGGCATGGAAGTTAAAGGTATCGTTAAGAACCTCACTG<br>ACTACGGTGCATTCGTTGATCTGGGCGGCGTTGACGGCCTGCTGCACATCACTGACATGGCC |

| | |
|---|---|
| | TGGAAACGCGTTAAGCATCCGAGCGAAATCGTCAACGTGGGCGACGAAATCACTGTTAAAGT<br>GCTGAAGTTCGACCGCGAACGTACCCGTGTATCCCTGGGCCTGAAACAGCTGGGCGAAGATC<br>CGTGGGTAGCTATCGCTAAACGTTATCCGGAAGGTACCAAACTGACTGGTCGCGTGACCAAC<br>CTGACCGACTACGGCTGCTTCGTTGAAATCGAAGAAGGCGTTGAAGGCCTGGTACACGTTTC<br>CGAAATGGACTGGACCAACAAAAACATCCACCCGTCCAAAGTTGTTAACGTTGGCGATGTAG<br>TGGAAGTTATGGTTCTGGATATCGACGAAGAACGTCGTCGTATCTCCCTGGGTCTGAAACAG<br>TGCAAAGCTAACCCGTGGCAGCAGTTCGCGGAAACCCACAACAAGGGCGACCGTGTTGAAG<br>GTAAAATCAAGTCTATCACTGACTTCGGTATCTTCATCGGCTTGGACGGCGGCATCGACGGCC<br>TGGTTCACCTGTCTGACATCTCCTGGAACGTTGCAGGCGAAGAAGCAGTTCGTGAATACAAA<br>AAAGGCGACGAAATCGCTGCAGTTGTTCTGCAGGTTGACGCAGAACGTGAACGTATCTCCCT<br>GGGCGTTAAACAGCTCGCAGAAGATCCGTTCAACAACTGGGTTGCTCTGAACAAGAAAGGC<br>GCTATCGTAACCGGTAAAGTAACTGCAGTTGACGCTAAAGGCGCAACCGTAGAACTGGCTGA<br>CGGCGTTGAAGGTTACCTGCGTGCTTCTGAAGCATCCCGTGACCGCGTTGAAGACGCTACCC<br>TGGTTCTGAGCGTTGGCGACGAAGTTGAAGCTAAATTCACCGGCGTTGATCGTAAAAACCGC<br>GCAATCAGCCTGTCTGTTCGTGCGAAAGACGAAGCTGACGAGAAAGATGCAATCGCAACTGT<br>TAACAAACAGGAAGATGCAAACTTCTCCAACAACGCAATGGCTGAAGCTTTCAAAGCAGCTA<br>AAGGCGAG**CTGGTGCCGCGCGGCAGC**CTCGAG |
| **pET28-rS1 R139K [*NcoI*, *XhoI*]** | CC*ATGG*GAACTGAATCTTTGCGGCATGCTCAACTCTTTGAAGAGTCCTTAAAAGAAATCGAA<br>ACCCGCCCGGGTTCTATCGTTCGTGGCGTTGTTGTTGCTATCGACAAAGACGTAGTACTGGTT<br>GACGCTGGTCTGAAATCTGAGTCCGCCATCCCGGCTGAGCAGTTCAAAAACGCCCAGGGCGA<br>GCTGGAAATCCAGGTAGGTGACGAAGTTGACGTTGCTCTGGACGCAGTAGAAGACGGCTTC<br>GGTGAAACTCTGCTGTCCCGTGAGAAAGCTAAACGTCACGAAGCCTGGATCACGCTGGAAAA<br>AGCTTACGAAGATGCTGAAACTGTTACCGGTGTTATCAACGGCAAAGTTAAGGGCGGCTTCA<br>CTGTTGAGCTGAACGGTATTCGTGCGTTCCTGCCAGGTTCTCTGGTAGACGTT**AAA**CCGGTG<br>CGTGACACTCTGCACCTGGAAGGCAAAGAGCTTGAATTTAAAGTAATCAAGCTGGATCAGAA<br>GCGCAACAACGTTGTTGTTTCTCGTCGTGCCGTTATCGAATCCGAAAACAGCGCAGAGCGCG<br>ATCAGCTGCTGGAAAACCTGCAGGAAGGCATGGAAGTTAAAGGTATCGTTAAGAACCTCACT<br>GACTACGGTGCATTCGTTGATCTGGGCGGCGTTGACGGCCTGCTGCACATCACTGACATGGC<br>CTGGAAACGCGTTAAGCATCCGAGCGAAATCGTCAACGTGGGCGACGAAATCACTGTTAAA<br>GTGCTGAAGTTCGACCGCGAACGTACCCGTGTATCCCTGGGCCTGAAACAGCTGGGCGAAG<br>ATCCGTGGGTAGCTATCGCTAAACGTTATCCGGAAGGTACCAAACTGACTGGTCGCGTGACC<br>AACCTGACCGACTACGGCTGCTTCGTTGAAATCGAAGAAGGCGTTGAAGGCCTGGTACACGT<br>TTCCGAAATGGACTGGACCAACAAAAACATCCACCCGTCCAAAGTTGTTAACGTTGGCGATG<br>TAGTGGAAGTTATGGTTCTGGATATCGACGAAGAACGTCGTCGTATCTCCCTGGGTCTGAAA<br>CAGTGCAAAGCTAACCCGTGGCAGCAGTTCGCGGAAACCCACAACAAGGGCGACCGTGTTG<br>AAGGTAAAATCAAGTCTATCACTGACTTCGGTATCTTCATCGGCTTGGACGGCGGCATCGAC<br>GGCCTGGTTCACCTGTCTGACATCTCCTGGAACGTTGCAGGCGAAGAAGCAGTTCGTGAATA<br>CAAAAAAGGCGACGAAATCGCTGCAGTTGTTCTGCAGGTTGACGCAGAACGTGAACGTATCT<br>CCCTGGGCGTTAAACAGCTCGCAGAAGATCCGTTCAACAACTGGGTTGCTCTGAACAAGAAA<br>GGCGCTATCGTAACCGGTAAAGTAACTGCAGTTGACGCTAAAGGCGCAACCGTAGAACTGG<br>CTGACGGCGTTGAAGGTTACCTGCGTGCTTCTGAAGCATCCCGTGACCGCGTTGAAGACGCT<br>ACCCTGGTTCTGAGCGTTGGCGACGAAGTTGAAGCTAAATTCACCGGCGTTGATCGTAAAAA<br>CCGCGCAATCAGCCTGTCTGTTCGTGCGAAAGACGAAGCTGACGAGAAAGATGCAATCGCA |

| | |
|---|---|
| | ACTGTTAACAAACAGGAAGATGCAAACTTCTCCAACAACGCAATGGCTGAAGCTTTCAAAGC<br>AGCTAAAGGCGAG**CTGGTGCCGCGCGGCAGC**CTCGAG |
| **pTAC-rS1 [*XhoI*, *SphI*]** | *ATG*AAGCTTCCTCGAGAGACTGAATCTTTTGCTCAACTCTTTGAAGAGTCCTTAAAAGAAATCGAAACC<br>CGCCCGGGTTCTATCGTTCGTGGCGTTGTTGTTGCTATCGACAAAGACGTAGTACTGGTTGACGCTGGT<br>CTGAAATCTGAGTCCGCCATCCCGGCTGAGCAGTTCAAAAACGCCCAGGGCGAGCTGGAAATCCAGGT<br>AGGTGACGAAGTTGACGTTGCTCTGGACGCAGTAGAAGACGGCTTCGGTGAAACTCTGCTGTCCCGTG<br>AGAAAGCTAAACGTCACGAAGCCTGGATCACGCTGGAAAAAGCTTACGAAGATGCTGAAACTGTTACC<br>GGTGTTATCAACGGCAAAGTTAAGGGCGGCTTCACTGTTGAGCTGAACGGTATTCGTGCGTTCCTGCC<br>AGGTTCTCTGGTAGACGTTCGTCCGGTGCGTGACACTCTGCACCTGGAAGGCAAAGAGCTTGAATTTA<br>AAGTAATCAAGCTGGATCAGAAGCGCAACAACGTTGTTGTTTCTCGTCGTGCCGTTATCGAATCCGAAA<br>ACAGCGCAGAGCGCGATCAGCTGCTGGAAAACCTGCAGGAAGGCATGGAAGTTAAAGGTATCGTTAA<br>GAACCTCACTGACTACGGTGCATTCGTTGATCTGGGCGGCGTTGACGGCCTGCTGCACATCACTGACAT<br>GGCCTGGAAACGCGTTAAGCATCCGAGCGAAATCGTCAACGTGGGCGACGAAATCACTGTTAAAGTGC<br>TGAAGTTCGACCGCGAACGTACCCGTGTATCCCTGGGCCTGAAACAGCTGGGCGAAGATCCGTGGGTA<br>GCTATCGCTAAACGTTATCCGGAAGGTACCAAACTGACTGGTCGCGTGACCAACCTGACCGACTACGG<br>CTGCTTCGTTGAAATCGAAGAAGGCGTTGAAGGCCTGGTACACGTTTCCGAAATGGACTGGACCAACA<br>AAAACATCCACCCGTCCAAAGTTGTTAACGTTGGCGATGTAGTGGAAGTTATGGTTCTGGATATCGACG<br>AAGAACGTCGTCGTATCTCCCTGGGTCTGAAACAGTGCAAAGCTAACCCGTGGCAGCAGTTCGCGGAA<br>ACCCACAACAAGGGCGACCGTGTTGAAGGTAAAATCAAGTCTATCACTGACTTCGGTATCTTCATCGGC<br>TTGGACGGCGGCATCGACGGCCTGGTTCACCTGTCTGACATCTCCTGGAACGTTGCAGGCGAAGAAGC<br>AGTTCGTGAATACAAAAAAGGCGACGAAATCGCTGCAGTTGTTCTGCAGGTTGACGCAGAACGTGAAC<br>GTATCTCCCTGGGCGTTAAACAGCTCGCAGAAGATCCGTTCAACAACTGGGTTGCTCTGAACAAGAAA<br>GGCGCTATCGTAACCGGTAAAGTAACTGCAGTTGACGCTAAAGGCGCAACCGTAGAACTGGCTGACG<br>GCGTTGAAGGTTACCTGCGTGCTTCTGAAGCATCCCGTGACCGCGTTGAAGACGCTACCCTGGTTCTGA<br>GCGTTGGCGACGAAGTTGAAGCTAAATTCACCGGCGTTGATCGTAAAAACCGCGCAATCAGCCTGTCT<br>GTTCGTGCGAAAGACGAAGCTGACGAGAAAGATGCAATCGCAACTGTTAACAAACAGGAAGATGCAA<br>ACTTCTCCAACAACGCAATGGCTGAAGCTTTCAAAGCAGCTAAAGGCGAGTGCATGCACGTAGAG |
| **S1 D1 [*NcoI*, *XhoI*]** | CC*ATGG*AGTCCTTAAAAGAAATCGAAACCCGCCCGGGTTCTATCGTTCGTGGCGTTGTTGTTG<br>CTATCGACAAAGACGTAGTACTGGTTGACGCTGGTCTGAAATCTGAGTCCGCCATCCCGGCT<br>GAGCAGTTCAAAAACGCCCAGGGCGAGCTGGAAATCCAGGTAGGTGACGAAGTTGACGTTG<br>CTCTGGACGCAGTAGAAGACGGCTTCGGTGAAACTCTGCTGTCCCGTGAGAAAGCTAAACGT<br>CACGAAGCC**CTGGTGCCGCGCGGCAGC**CTCGAG |
| **S1 D2 [*NcoI*, *XhoI*]** | CC*ATGG*CCTGGATCACGCTGGAAAAAGCTTACGAAGATGCTGAAACTGTTACCGGTGTTATC<br>AACGGCAAAGTTAAGGGCGGCTTCACTGTTGAGCTGAACGGTATTCGTGCGTTCCTGCCAGG<br>TTCTCTGGTAGACGTTCGTCCGGTGCGTGACACTCTGCACCTGGAAGGCAAAGAGCTTGAAT<br>TTAAAGTAATCAAGCTGGATCAGAAGCGCAACAACGTTGTTGTTTCTCGTCGTGCCGTTATCG<br>AATCCGAAAACAGCGCAGAG**CTGGTGCCGCGCGGCAGC**CTCGAG |
| **S1 D2 R139A [*NcoI*, *XhoI*]** | CC*ATGG*CCTGGATCACGCTGGAAAAAGCTTACGAAGATGCTGAAACTGTTACCGGTGTTATC<br>AACGGCAAAGTTAAGGGCGGCTTCACTGTTGAGCTGAACGGTATTCGTGCGTTCCTGCCAGG<br>TTCTCTGGTAGACGTT**GCC**CCGGTGCGTGACACTCTGCACCTGGAAGGCAAAGAGCTTGAAT<br>TTAAAGTAATCAAGCTGGATCAGAAGCGCAACAACGTTGTTGTTTCTCGTCGTGCCGTTATCG<br>AATCCGAAAACAGCGCAGAG**CTGGTGCCGCGCGGCAGC**CTCGAG |
| **S1 D2 R139K [*NcoI*, *XhoI*]** | CC*ATGG*CCTGGATCACGCTGGAAAAAGCTTACGAAGATGCTGAAACTGTTACCGGTGTTATC<br>AACGGCAAAGTTAAGGGCGGCTTCACTGTTGAGCTGAACGGTATTCGTGCGTTCCTGCCAGG |

| | |
|---|---|
| | TTCTCTGGTAGACGTT**AAA**CCGGTGCGTGACACTCTGCACCTGGAAGGCAAAGAGCTTGAAT<br>TTAAAGTAATCAAGCTGGATCAGAAGCGCAACAACGTTGTTGTTTCTCGTCGTGCCGTTATCG<br>AATCCGAAAACAGCGCAGAG**CTGGTGCCGCGCGGCAGC**CTCGAG |
| **S1 D3 [*NcoI*, *XhoI*]** | CC*ATG*GCCCGCGATCAGCTGCTGGAAAACCTGCAGGAAGGCATGGAAGTTAAAGGTATCGT<br>TAAGAACCTCACTGACTACGGTGCATTCGTTGATCTGGGCGGCGTTGACGGCCTGCTGCACA<br>TCACTGACATGGCCTGGAAACGCGTTAAGCATCCGAGCGAAATCGTCAACGTGGGCGACGA<br>AATCACTGTTAAAGTGCTGAAGTTCGACCGCGAACGTACCCGTGTATCCCTGGGCCTGAAAC<br>AGCTGGGCGAAGATCCG**CTGGTGCCGCGCGGCAGC**CTCGAG |
| **S1 D4 [*NcoI*, *XhoI*]** | CC*ATG*GCCTGGGTAGCTATCGCTAAACGTTATCCGGAAGGTACCAAACTGACTGGTCGCGTG<br>ACCAACCTGACCGACTACGGCTGCTTCGTTGAAATCGAAGAAGGCGTTGAAGGCCTGGTACA<br>CGTTTCCGAAATGGACTGGACCAACAAAAACATCCACCCGTCCAAAGTTGTTAACGTTGGCG<br>ATGTAGTGGAAGTTATGGTTCTGGATATCGACGAAGAACGTCGTCGTATCTCCCTGGGTCTG<br>AAACAGTGCAAAGCTAACCCG**CTGGTGCCGCGCGGCAGC**CTCGAG |
| **S1 D5 [*NcoI*, *XhoI*]** | CC*ATG*GCCTGGCAGCAGTTCGCGGAAACCCACAACAAGGGCGACCGTGTTGAAGGTAAAAT<br>CAAGTCTATCACTGACTTCGGTATCTTCATCGGCTTGGACGGCGGCATCGACGGCCTGGTTCA<br>CCTGTCTGACATCTCCTGGAACGTTGCAGGCGAAGAAGCAGTTCGTGAATACAAAAAAGGCG<br>ACGAAATCGCTGCAGTTGTTCTGCAGGTTGACGCAGAACGTGAACGTATCTCCCTGGGCGTT<br>AAACAGCTCGCAGAAGATCCG**CTGGTGCCGCGCGGCAGC**CTCGAG |
| **S1 D6 [*NcoI*, *XhoI*]** | CC*ATG*GCCTTCAACAACTGGGTTGCTCTGAACAAGAAAGGCGCTATCGTAACCGGTAAAGTA<br>ACTGCAGTTGACGCTAAAGGCGCAACCGTAGAACTGGCTGACGGCGTTGAAGGTTACCTGC<br>GTGCTTCTGAAGCATCCCGTGACCGCGTTGAAGACGCTACCCTGGTTCTGAGCGTTGGCGAC<br>GAAGTTGAAGCTAAATTCACCGGCGTTGATCGTAAAAACCGCGCAATCAGCCTGTCTGTTCG<br>TGCGAAAGACGAAGCTGACGAGAAA**CTGGTGCCGCGCGGCAGC**CTCGAG |
| **S1 domain of PNPase [*NcoI*, *XhoI*]** | CC*ATG*GCAGAAATCGAAGTGGGCCGCGTCTACACTGGTAAAGTGACCCGTATCGTTGACTTT<br>GGCGCATTTGTTGCCATCGGCGGCGGTAAAGAAGGTCTGGTCCACATCTCTCAAATCGCTGA<br>CAAACGCGTTGAGAAAGTGACCGATTACCTGCAGATGGGTCAGGAAGTACCGGTGAAAGTT<br>CTGGAAGTTGATCGCCAGGGCCGTATCCGTCTGAGCATTAAAGAAGCGACTGAGCAGTCTCA<br>ACCTGCTGCA**CTGGTGCCGCGCGGCAGC**CTCGAG |
| **pET28 ARH1 [*NcoI*, *XhoI*]** | CC*ATG*GAAAAATACGTCGCCGCGATGGTTTTGTCAGCTGCTGGCGATGCTTTGGGATATTAT<br>AATGGAAAGTGGGAATTTCTTCAGGACGGGGAGAAAATTCATCGTCAACTGGCTCAATTAGG<br>GGGGCTGGATGCTCTGGACGTTGGCCGTTGGCGTGTGTCTGATGATACTGTCATGCACTTGG<br>CAACAGCCGAGGCTTTGGTCGAGGCCGGAAAGGCTCCAAAACTGACTCAGCTTTATTATTTG<br>TTAGCCAAGCACTATCAGGATTGCATGGAAGATATGGACGGTCGCGCACCCGGGGGTGCGT<br>CTGTACACAACGCGATGCAGCTTAAACCTGGGAAACCGAATGGCTGGCGTATCCCATTTAAC<br>TCGCATGAAGGAGGGTGTGGCGCGGCGATGCGCGCGATGTGTATCGGTTTGCGTTTTCCGC<br>ATCACTCTCAATTAGACACACTGATCCAAGTATCGATCGAGTCAGGACGTATGACCCATCATC<br>ACCCGACAGGGTACCTTGGCGCACTTGCGTCCGCCTTATTCACGGCCTATGCGGTAAATAGCC<br>GCCCTCCATTGCAGTGGGGTAAGGGACTTATGGAGCTTTTGCCAGAGGCTAAAAAATACATT<br>GTCCAATCCGGGTACTTTGTGGAAGAAAATTTACAGCATTGGTCTTATTTTCAAACGAAGTGG<br>GAAAACTATCTTAAACTGCGTGGAATCTTGGACGGCGAGAGTGCTCCAACATTCCCTGAATCT<br>TTTGGCGTTAAAGAGCGCGACCAGTTCTACACTTCGTTGTCATATAGTGGCTGGGGCGGTTC |

| | |
|---|---|
| | ATCTGGGCATGATGCCCCCATGATCGCGTATGACGCGGTGCTGGCGGCGGGAGACTCCTGG<br>AAAGAGCTTGCGCACCGCGCCTTCTTTCACGGAGGTGACTCGGATTCGACCGCAGCCATTGC<br>TGGATGTTGGTGGGGCGTCATGTACGGATTTAAGGGCGTCAGCCCCAGCAACTACGAAAAA<br>TTAGAGTATCGCAATCGCCTTGAGGAAACAGCTCGCGCACTTTACTCGCTGGGTAGTAAAGA<br>AGACACTGTTATCTCGCTG**CTGGTGCCGCGCGGCAGC**CTCGAG |
| **pET ARH1 D55A, D56A [*NcoI, XhoI*]** | CCATGGAAAAATACGTCGCCGCGATGGTTTTGTCAGCTGCTGGCGATGCTTTGGGATATTAT<br>AATGGAAAGTGGGAATTTCTTCAGGACGGGGAGAAAATTCATCGTCAACTGGCTCAATTAGG<br>GGGGCTGGATGCTCTGGACGTTGGCCGTTGGCGTGTGTCT**GCGGCG**ACTGTCATGCACTTGG<br>CAACAGCCGAGGCTTTGGTCGAGGCCGGAAAGGCTCCAAAACTGACTCAGCTTTATTATTTG<br>TTAGCCAAGCACTATCAGGATTGCATGGAAGATATGGACGGTCGCGCACCCGGGGGTGCGT<br>CTGTACACAACGCGATGCAGCTTAAACCTGGGAAACCGAATGGCTGGCGTATCCCATTTAAC<br>TCGCATGAAGGAGGGTGTGGCGCGGCGATGCGCGCGATGTGTATCGGTTTGCGTTTTCCGC<br>ATCACTCTCAATTAGACACACTGATCCAAGTATCGATCGAGTCAGGACGTATGACCCATCATC<br>ACCCGACAGGGTACCTTGGCGCACTTGCGTCCGCCTTATTCACGGCCTATGCGGTAAATAGCC<br>GCCCTCCATTGCAGTGGGGTAAGGGACTTATGGAGCTTTTGCCAGAGGCTAAAAAATACATT<br>GTCCAATCCGGGTACTTTGTGGAAGAAAATTTACAGCATTGGTCTTATTTTCAAACGAAGTGG<br>GAAAACTATCTTAAACTGCGTGGAATCTTGGACGGCGAGAGTGCTCCAACATTCCCTGAATCT<br>TTTGGCGTTAAAGAGCGCGACCAGTTCTACACTTCGTTGTCATATAGTGGCTGGGGCGGTTC<br>ATCTGGGCATGATGCCCCCATGATCGCGTATGACGCGGTGCTGGCGGCGGGAGACTCCTGG<br>AAAGAGCTTGCGCACCGCGCCTTCTTTCACGGAGGTGACTCGGATTCGACCGCAGCCATTGC<br>TGGATGTTGGTGGGGCGTCATGTACGGATTTAAGGGCGTCAGCCCCAGCAACTACGAAAAA<br>TTAGAGTATCGCAATCGCCTTGAGGAAACAGCTCGCGCACTTTACTCGCTGGGTAGTAAAGA<br>AGACACTGTTATCTCGCTG**CTGGTGCCGCGCGGCAGC**CTCGAG |
| **pET rL2 [NcoI, XhoI]** | CCATGGGCGCAGTTGTTAAATGTAAACCGACATCTCCGGGTCGTCGCCACGTAGTTAAAGTG<br>GTTAACCCTGAGCTGCACAAGGGCAAACCTTTTGCTCCGTTGCTGGAAAAAAACAGCAAATC<br>CGGTGGTCGTAACAACAATGGCCGTATCACCACTCGTCATATCGGTGGTGGCCACAAGCAGG<br>CTTACCGTATTGTTGACTTCAAACGCAACAAAGACGGTATCCCGGCAGTTGTTGAACGTCTTG<br>AGTACGATCCGAACCGTTCCGCGAACATCGCGCTGGTTCTGTACAAAGACGGTGAACGCCGT<br>TACATCCTGGCCCCTAAAGGCCTGAAAGCTGGCGACCAGATTCAGTCTGGCGTTGATGCTGC<br>AATCAAACCAGGTAACACCCTGCCGATGCGCAACATCCCGGTTGGTTCTACTGTTCATAACGT<br>AGAAATGAAACCAGGTAAAGGCGGTCAGCTGGCACGTTCCGCTGGTACTTACGTTCAGATCG<br>TTGCTCGTGATGGTGCTTATGTCACCCTGCGTCTGCGTTCTGGTGAAATGCGTAAAGTAGAAG<br>CAGACTGCCGTGCAACTCTGGGCGAAGTTGGCAATGCTGAGCATATGCTGCGCGTTCTGGGT<br>AAAGCAGGTGCTGCACGCTGGCGTGGTGTTCGTCCGACCGTTCGCGGTACCGCGATGAACCC<br>GGTAGACCACCCACATGGTGGTGGTGAAGGTCGTAACTTTGGTAAGCACCCGGTAACTCCGT<br>GGGGCGTTCAGACCAAAGGTAAGAAGACCCGCAGCAACAAGCGTACTGATAAATTCATCGT<br>ACGTCGCCGTAGCAAACTCGAG |

**Supplementary Table 8: Primers used in this study.** Corresponding restriction site in bold, underlined; mutation in bold and red

| Primer | Sequence (5' to 3') |
|---|---|
| Fwd Qβ T7 | TAATACGACTCACTATTATCTTGATACTACCTTTAG |

| | |
|---|---|
| Rev Qβ | CATGATCAAATTGACCCAAAGTTTCAACGCTTTACGCG |
| Fwd RNAI T7 | TAATACGACTCACTATAACAGTATTTGGTATC |
| Rev RNAI | ACAAAAAAACCACCGCTACCAGCGGTGGTTTGTTTGCC |
| Fwd Alt NcoI | ATCGAC**CCATGG**GAGAACTTATTACAGAATTATTTGACG |
| Rev Alt XhoI | ATTCGA**CTCGAG**GCTGCCGCGCGGCACCAGTCCTTGAACGAACTTGTAAGGCATG |
| Fwd ModA NcoI | ATCGA**CCATGG**GAAAATACTCAGTAATGCAACTAAAAG |
| Rev ModA XhoI | ATCGTA**CTCGAG**GCTGCCGCGCGGCACCAGTAGATTAAATCCTTCAAAATCAAG |
| Fwd ModB NcoI | ATCGAC**CCATGG**GAATTATTAATCTTGCAGATGTTG |
| Rev ModB XhoI | ACTTAG**CTCGAG**GCTGCCGCGCGGCACCAGTTCAACTACCTCAATATCATAAAC |
| Fwd rS1 NcoI | ATCGAC**CCATGG**GAACTGAATCTTTTGCTCAACTCTTTGAAGAGTCC |
| Rev rS1 XhoI | ATTCGA**CTCGAG**GCTGCCGCGCGGCACCAGCTCGCCTTTAGCTGCTTTG |
| Fwd rS1-pTAC XhoI | ATGAAGCTTC**CTCGAG**AGACTGAATCTTTTGCTCAACTCTTTGAAGAGTCC |
| Rev rS1-pTAC SphI | CTCTACGT**GCATGC**ACTCGCCTTTAGCTGCTTTGAAAGCTTCAGCC |
| Fwd NcoI rS1 D1 | ATCGAC**CCATGG**AGTCCTTAAAAGAAATCGAAACCCGCCCGGG |
| Rev XhoI rS1 D1 | TGGTG**CTCGAG**GCTGCCGCGCGGCACCAGGGCTTCGTGACGTTTAGCTTTCTCACGGG |
| Fwd NcoI rS1 D2 | ATCGAC**CCATGG**CCTGGATCACGCTGGAAAAAGCTTACGAAGATGCTGAAAC |
| Rev XhoI rS1 D2 | GGTG**CTCGAG**GCTGCCGCGCGGCACCAGCTCTGCGCTGTTTTCGGATTCGATAACGGCAC |
| Fwd NcoI rS1 D3 | ATCGAC**CCATGG**CCCGCGATCAGCTGCTGGAAAACCTGCAGGAAGG |
| Rev XhoI rS1 D3 | TGGTG**CTCGAG**GCTGCCGCGCGGCACCAGCGGATCTTCGCCCAGCTGTTTCAGGCCCAGG |
| Fwd NcoI rS1 D4 | ATCGAC**CCATGG**CCTGGGTAGCTATCGCTAAACGTTATCCGGAAGG |
| Rev XhoI rS1 D4 | TGGTG**CTCGAG**GCTGCCGCGCGGCACCAGCGGGTTAGCTTTGCACTGTTTCAGACCCAGGGAG |
| Fwd NcoI rS1 D5 | ATCGAC**CCATGG**CCTGGCAGCAGTTCGCGGAAACCCACAACAAGGGCGACCGTGTTG |

| | |
|---|---|
| Rev XhoI S1 D5 | TGGTG**CTCGAG**GCTGCCGCGCGGCACCAGCGGATCTTCTGCGAGCTGTTTAACGCCCAGGGAGATACG |
| Fwd NcoI rS1 D6 | ATCGAC**CCATGG**CCTTCAACAACTGGGTTGCTCTGAACAAGAAAGGCGCTATCG |
| Rev XhoI rS1 D6 | TGGTG**CTCGAG**GCTGCCGCGCGGCACCAGTTTCTCGTCAGCTTCGTCTTTCGCACGAACAGACAGG |
| Fwd NcoI PNPase rS1 binding | ATCGAC**CCATGG**CAGAAATCGAAGTGGGCCGCGTCTACACTGGTAAAGTGACCCG |
| Rev XhoI PNPase rS1 binding | TGGTG**CTCGAG**GCTGCCGCGCGGCACCAGTGCAGCAGGTTGAGACTGCTCAGTCGCTTC |
| Fwd ARH1 NcoI | TGCAG**CCATGG**AAAAATACGTCGCCGCGATG |
| Rev ARH1 XhoI | GTGGTG**CTCGAG**GCTGCCGCGCGGCACCAG |
| Fwd rS1 R139A | CTGGTAGACGTT**GCC**CCGGTGCGTGACACTC |
| Fwd rS1 R139K | CTGGTAGACGTT**AAA**CCGGTGCGTGACACTC |
| Rev rS1 R139 | AGAACCTGGCAGGAACGCACGAATACCG |
| Fwd ARH1 D55,56A | GGCCGTTGGCGTGTGTCT**GCGGCG**ACTGTCATGCACTTGGC |
| Rev ARH1 D55,56A | AACGTCCAGAGCATCCAGCCCCCCTAA |
| Fwd ModB R73A | CCTTATCAATTATAT**GCG**GGTATATCAAAATCG |
| Rev ModB R73A | CGATTTAGTTAAATGCTTTTTAATGATTTC |
| Fwd ModB G74A | GACAAAAGAACTCATTAAAGATTTAC |
| Rev ModB G74A | GATTTTGATAT**TGCCGC**ATATAATTGATAAGGCG |
| Fwd ModB DS_SPCas | AATTATATCGGTTTTAGAGCTATGCTGTTTTGAATGGTCC |
| Rev ModB DS_SPCas | GATAAGGCGAGCTAGCACTGTACCTAGGACTGAGC |
| Fwd ModB amplification T4 genome | CCAAGAATGGTCATCTGGTTTATTAG |
| Rev ModB amplification T4 genome | CCGCCTTGGGCTCCCTGG |

| | |
|---|---|
| Fwd sequencing ModB T4 genome | CAGTTATCTATAAAAGCTGAAAG |
| Rev sequencing ModB T4 genome | CTTTCCAATTGGAATCATCCATTC |
| rpsA homologous downstream fwd | TTCTCTGACTCTTCGGGATTTTTATTC |
| rpsA homologous downstream rev | AGGCAAATTAAGCGGCTGCTG |
| Terminator region fwd | TTCTCTGACTCTTCGGGATTTTTATTC |
| Terminator region rev | AGGACGAAACCTGCAATCTGTC |
| FRT pKD4 fwd | TCGGAATAAAAATCCCGAAGAGTCAGAGAAGTCCATATGAATATCCTCCTTAGTTC |
| FRT pKD4 rev | GTTTACTTGACAGATTGCAGGTTTCGTCCTGTGTAGGCTGGAGCTGCTTC |
| 5_70 left rev rpsA | AGGACGAAACCTGCAATCTGTC |
| 5_fwd_rS1 amplification | GGCGTTGATCGTAAAAACCGC |
| Fwd NcoI rL2 | **CCATGG**GCGCAGTTGTTAAATGTAAACCG |
| Rev XhoI rL2 | **CTCGAG**TTTGCTACGGCGACGTACGATG |
| adenylated RNA-3'-adapter | /5rApp/CNNNNNNAGATCGGAAGAGCACACGTCTG/3SpC3/ |
| RT primer | CAGACGTGTGCTCTTCCGAT |
| cDNA anchor fwd | ACACGACGCTCTTCCGATCTGGG |
| cDNA anchor rev | /5Phos/CAGATCGGAAGAGCGTCGTGTCCC/3SpC3/ |
| qPCR acpP fwd | CGTGGTAAGACCTGCCGG |
| qPCR acpP rev | CTCAACGGTGTCAAGAGAATCCAAAAC |
| qPCR gadY fwd | GAGCACAAAGTTTCCCGTGC |
| qPCR gadY rev | AAACCCGGCATAGGGGACC |

| | |
|---|---|
| qPCR mcaS fwd | AAAATAGAGTCTGTCGACATCCGC |
| qPCR mcaS rev | CACCGGCGCAGAGGAGAC |
| qPCR oxyS fwd | AAAAGCGGATCCTGGAGATCC |
| qPCR oxyS rev | GAAACGGAGCGGCACCTC |
| qPCR rnaC fwd | CGTTGCGGCAACCTTGTC |
| qPCR rnaC rev | AAAAATATTGAGTAGCGTCAACTAC |

**Supplementary Table 9: Strains and plasmids used in this study**

| Name | Description | Reference or resource |
|---|---|---|
| ***E. coli* strain B** | *E. coli* strain applied for bacteriophage T4 infection | DMSZ, Escherichia coli (Migula 1895) Castellani and Chalmers 1919 (DSM 613, ATCC 11303) |
| ***E. coli* strain B pTAC rS1** | *E. coli* strain B expressing His-tagged rS1 under the control of *E. coli* RNA polymerase promoter | This study |
| ***E. coli* BL21 (DE3) pET16 RNase E (1-529)** | *E. coli* strain expressing His-tagged catalytic domain of RNase E (1-529) | Plasmid was a kind gift from Prof. Dr. Ben Luisi [7] |
| ***E. coli* BL21 (DE3) pET 28 NudC V157A, E174A, E177A, E178A** | *E. coli* strain expressing His-tagged inactive Mutant of NudC | [8] |
| ***E. coli* BL21 (DE3) pET 28 rS1** | *E. coli* strain expressing His-tagged rS1 | This study |
| ***E. coli* BL21 (DE3) pET 28 rS1 R139K** | *E. coli* strain expressing His-tagged rS1 R139K variant | This study |
| ***E. coli* BL21 (DE3) pET 28 rS1 R139A** | *E. coli* strain expressing His-tagged rS1 R139A variant | This study |
| ***E. coli* BL21 (DE3) pET 28 rS1 D1** | *E. coli* strain expressing His-tagged rS1 D1 | This study |
| ***E. coli* BL21 (DE3) pET 28 rS1 D2** | *E. coli* strain expressing His-tagged rS1 D2 | This study |
| ***E. coli* BL21 (DE3) pET 28 rS1 D2 R139K** | *E. coli* strain expressing His-tagged rS1 D2 R139K | This study |

| | | |
|---|---|---|
| ***E. coli*** **BL21 (DE3) pET 28 rS1 D2 R139A** | *E. coli* strain expressing His-tagged rS1 D2 R139A | This study |
| ***E. coli*** **BL21 (DE3) pET 28 rS1 D3** | *E. coli* strain expressing His-tagged rS1 D3 | This study |
| ***E. coli*** **BL21 (DE3) pET 28 rS1 D4** | *E. coli* strain expressing His-tagged rS1 D4 | This study |
| ***E. coli*** **BL21 (DE3) pET 28 rS1 D5** | *E. coli* strain expressing His-tagged rS1 D5 | This study |
| ***E. coli*** **BL21 (DE3) pET 28 rS1 D6** | *E. coli* strain expressing His-tagged rS1 D6 | This study |
| ***E. coli*** **BL21 (DE3) pET 28 Alt** | *E. coli* strain expressing His-tagged Alt | This study |
| ***E. coli*** **BL21 (DE3) pET 28 ModA** | *E. coli* strain expressing His-tagged ModA | This study |
| ***E. coli*** **BL21 (DE3) pET 28 ModB** | *E. coli* strain expressing His-tagged ModB | This study |
| ***E. coli*** **BL21 (DE3) pET 28 ModB R73A, G74A** | *E. coli* strain expressing His-tagged ModB with point mutations R73A and G74A | This study |
| ***E. coli*** **BL21 (DE3) pET 28 NudC** | *E. coli* strain expressing His-tagged NudC | [8] |
| ***E. coli*** **BL21 (DE3) pET 28 PNPase S1 domain** | *E. coli* strain expressing His-tagged PNPase S1 domain | This study |
| ***E. coli*** **BL21 (DE3) pET 28 ARH1** | *E. coli* strain expressing His-tagged ARH1 | This study |
| ***E. coli*** **BL21 (DE3) pET 28 ARH1 D55A, D56A** | *E. coli* strain expressing His-tagged ARH1 D55A, D56A | This study |
| ***E. coli*** **DHα DS_SPCas_ModB** | *E. coli* strain expressing CRISPR-Cas9 system for cleavage of *modB* | This study |
| ***E. coli*** **DHα DS_SPCas_ModB pET28 ModB R73A, G74A** | *E. coli* strain for editing of *modB* within T4 phage genome | This study |
| ***E. coli*** **BL21 (DE3) pET28 rL2** | *E. coli* strain expressing His-tagged rL2 | This study |

| | | |
|---|---|---|
| ***E. coli* B strain with endogenously His-tagged rS1** | *E. coli* strain with endogenous expression of rS1 with a His-tag fusion at the C-terminus | This study |
| **T4 WT** | Wild-type bacteriophage T4 | Escherichia phage T4, DSM 4505, DSMZ, Braunschweig, Germany) |
| **T4 ModB R73A, G74A** | T4 phage mutant carrying inactive ModB version ModB R73A, G74A | This study |

**References**


1 Jumper, J. *et al.* Highly accurate protein structure prediction with AlphaFold. *Nature* **596**, 583-589 (2021). https://doi.org:10.1038/s41586-021-03819-2

2 MacLean, B. *et al.* Skyline: an open source document editor for creating and analyzing targeted proteomics experiments. *Bioinformatics* **26**, 966-968 (2010). https://doi.org:10.1093/bioinformatics/btq054

3 Winz, M. L. *et al.* Capture and sequencing of NAD-capped RNA sequences with NAD captureSeq. *Nat Protoc* **12**, 122-149 (2017). https://doi.org:10.1038/nprot.2016.163

4 Qureshi, N. S., Bains, J. K., Sreeramulu, S., Schwalbe, H. & Fürtig, B. Conformational switch in the ribosomal protein S1 guides unfolding of structured RNAs for translation initiation. *Nucleic Acids Res* **46**, 10917-10929 (2018). https://doi.org:10.1093/nar/gky746

5 Di Tommaso, P. *et al.* T-Coffee: a web server for the multiple sequence alignment of protein and RNA sequences using structural information and homology extension. *Nucleic Acids Res* **39**, W13-17 (2011). https://doi.org:10.1093/nar/gkr245

6 Watson, Z. L. *et al.* Structure of the bacterial ribosome at 2 A resolution. *Elife* **9**, e60482 (2020). https://doi.org:10.7554/eLife.60482

7 Callaghan, A. J. *et al.* Structure of Escherichia coli RNase E catalytic domain and implications for RNA turnover. *Nature* **437**, 1187-1191 (2005). https://doi.org:10.1038/nature04084

8 Cahova, H., Winz, M. L., Höfer, K., Nübel, G. & Jäschke, A. NAD captureSeq indicates NAD as a bacterial cap for a subset of regulatory RNAs. *Nature* **519**, 374-377 (2015). https://doi.org:10.1038/nature14020
